# Comprehensive volumetric phenotyping of the neonatal brain in Down syndrome

**DOI:** 10.1101/2022.09.30.510205

**Authors:** Abi Fukami - Gartner, Ana A. Baburamani, Ralica Dimitrova, Prachi A. Patkee, Olatz Ojinaga Alfageme, Alexandra F. Bonthrone, Daniel Cromb, Alena Uus, Serena J. Counsell, Joseph V. Hajnal, Jonathan O’Muircheartaigh, Mary A. Rutherford

## Abstract

Down syndrome (DS) is the most common genetic cause of intellectual disability with a wide spectrum of neurodevelopmental outcomes. Magnetic resonance imaging (MRI) has been used to investigate differences in whole and/or regional brain volumes in DS from infancy to adulthood. However, to date, there have been relatively few *in vivo* neonatal brain imaging studies in DS, despite the presence of clearly identifiable characteristics at birth. Improved understanding of early brain development in DS is needed to assess phenotypic severity and identify appropriate time windows for early intervention. In this study, we used *in vivo* brain MRI to conduct a comprehensive volumetric phenotyping of the neonatal brain in DS. Using a robust cross-sectional reference sample of close to 500 preterm- to term-born control neonates, we have performed normative modelling and quantified volumetric deviation from the normative mean in 25 individual infants with DS [postmenstrual age at scan, median (range) = 40.57 (32.43 – 45.57) weeks], corrected for sex, age at scan and age from birth. We found that absolute whole brain volume was significantly reduced in neonates with DS (pFDR <0.0001), as were most underlying absolute tissue volumes, except for the lentiform nuclei and the extracerebral cerebrospinal fluid (eCSF), which were not significantly different, and the lateral ventricles, which were significantly enlarged (pFDR <0.0001). Relative volumes, adjusting for underlying differences in whole brain volume, revealed a dynamic shift in brain proportions in neonates with DS. In particular, the cerebellum, as well as the cingulate, frontal, insular and occipital white matter (WM) segments were significantly reduced in proportion (pFDR <0.0001). Conversely, deep grey matter (GM) structures, such as the thalami and lentiform nuclei, as well as CSF-filled compartments, such as the eCSF and the lateral ventricles were significantly enlarged in proportion (pFDR <0.0001). We also observed proportionally reduced frontal and occipital lobar volumes, in contrast with proportionally enlarged temporal and parietal lobar volumes. Lastly, we noted age-related volumetric differences between neonates with and without a congenital heart defect (CHD), indicating that there may be a baseline brain phenotype in neonates with DS, which is further altered in the presence of CHD. In summary, we provide a comprehensive volumetric phenotyping of the neonatal brain in DS and observe many features that appear to follow a developmental continuum, as noted in older age cohorts. There are currently no paediatric longitudinal neuroimaging investigations in DS, starting from the earliest time points, which greatly impedes our understanding of the developmental continuum of neuroanatomical parameters in DS. Whilst life expectancy of individuals with DS has greatly improved over the last few decades, early interventions may be essential to help improve outcomes and quality of life.

**GRAPHICAL ABSTRACT:** 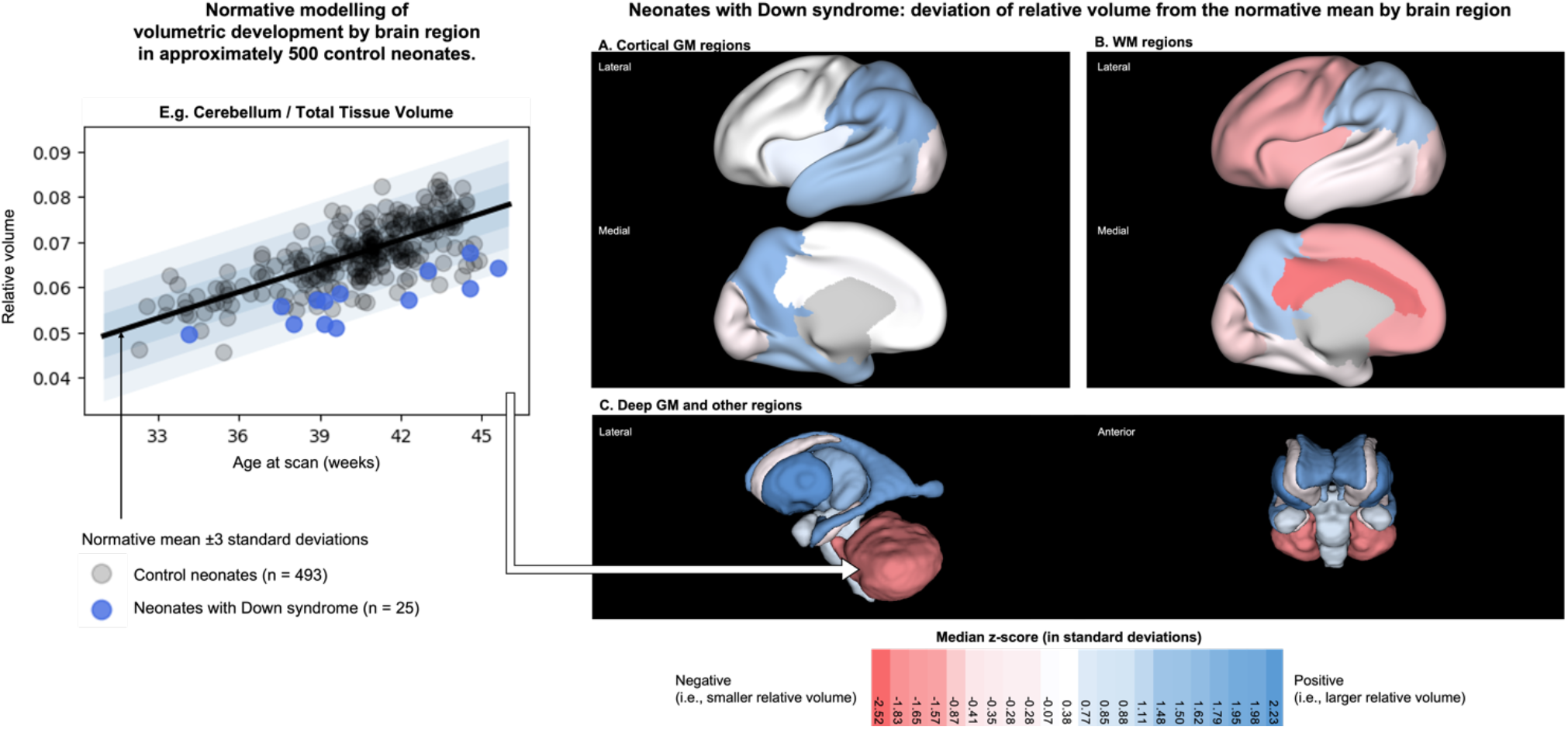

## INTRODUCTION

Down syndrome (DS) is caused by the partial or complete triplication of human chromosome 21 (*Hsa21*; trisomy 21)^1–3^ and is the most common genetic developmental disorder in humans^4,5^. It is estimated that DS affects approximately 1 in 750^6–8^ to 1 in 1000^9^ live births globally. DS is a complex disorder, which affects multiple body systems, in particular the neurological, cardiovascular and musculoskeletal systems^2,3,5^.

DS represents the most common genetic cause of intellectual disability with a wide spectrum of neurodevelopmental outcomes^10^. The majority of individuals are classified as having mild to moderate disability, although a wide and largely unexplained spectrum of intelligence quotients are noted^10–12^. The cognitive profile of individuals with DS demonstrates strengths in visual learning, but weaknesses in expressive language, verbal working memory, and episodic memory^10,12–14^.

Clinical comorbidities observed at birth, include congenital heart defects, gastrointestinal malformations (e.g., duodenal atresia, Hirschsprung’s disease), haematological (e.g., thrombocytopenia, polycythaemia), endocrine (e.g., hypothyroidism), hearing and visual impairments amongst others^15,16^. Characteristic physical features include a brachycephalic cranium with flat occiput, as identified by an increased biparietal to occipitofrontal diameter ratio^17,18^.

Congenital heart defects (CHD) are present in approximately 50% of neonates with DS, the most common of which are atrioventricular septal defects (AVSD, approximately 42% of CHDs in DS), ventricular septal defects (VSD, ~22%) and atrial septal defects (ASD, ~16%), whilst coarctation of the aorta, hypoplastic left heart syndrome, tetralogy of Fallot, and persistent *patent ductus arteriosus* or *patent foramen ovale* are also noted in smaller numbers^19,20^. Children with CHD, with^21^ and without DS^22^, are at risk of neurodevelopmental impairments. Children with DS, with an associated AVSD, have poorer gross motor and cognitive skills, and lower scores in expressive and receptive language, in comparison to children with DS with a structurally normal heart^21,23,24^. Children with CHD, without any known genetic diagnoses, are at increased risk of developmental deficits in domains including executive functioning, speech and language, motor coordination and cognition^25–27^.

Magnetic resonance imaging (MRI) has been used to investigate differences in whole and/or regional brain volumes in DS during toddlerhood^28–30^, early childhood^30–32^, middle childhood^30,33–36^, adolescence and young adulthood^30,37–40^ (see Hamner et al. (2018) for a review of paediatric neuroimaging in DS)^41^. However, to date, there have been relatively few *in vivo* fetal or neonatal brain imaging studies in DS, despite the presence of clearly identifiable characteristics *in utero* and at birth respectively^30,42–44^. Improved understanding of early brain development in DS is needed to assess phenotypic severity, identify appropriate time windows for early intervention, and improve parental counselling. Previous studies in fetuses and neonates with DS showed that whole brain and cerebellar volumes were smaller than age-matched euploid controls from the second trimester (< 28 weeks gestational age, GA)^42^, whilst total cortical grey matter (GM) volume was reduced from the third trimester onwards (> 28 weeks GA)^42,43^. To date, volumetric differences in regional cortical GM, white matter (WM), deep GM and other brain segments have not been examined in detail in neonates with DS. Furthermore, prior group-level analyses have not quantified individual variability within a highly heterogenous cohort.

In this study, we used *in vivo* brain MRI to conduct a comprehensive volumetric phenotyping of the neonatal brain in DS. Using a robust cross-sectional reference sample of close to 500 preterm- to term-born control neonates, we have performed normative modelling and quantified individual volumetric deviation from the normative model mean in 25 neonates with DS, corrected for sex, age at scan and age from birth.

## MATERIAL & METHODS

### Ethical approval

National Research Ethics Committees (REC) based in London (UK) provided ethical approval for the following studies: quantification of fetal brain development using MRI (Trisomy 21 group, denoted ‘T21’) [07/H0707/105], early brain imaging in Down syndrome (‘eBiDS’) [19/LO/0667] and the developing Human Connectome Project (‘dHCP’) [14/LO/1169]. In accordance with the declaration of Helsinki, informed written parental consent was obtained before MRI in all above studies, and before developmental follow-up in the dHCP.

### Participants

#### Neonates with Down syndrome

Neonates with a confirmed postnatal diagnosis of DS were recruited from the neonatal unit and postnatal wards at St Thomas’ Hospital London and invited for a neonatal scan up to 46 weeks post-menstrual age (PMA). Additionally, former fetal scan participants with a confirmed postnatal diagnosis of DS were invited for a neonatal scan if they had consented to be contacted post-delivery^42^. A total of 36 neonates with DS were recruited to the T21 and eBiDS studies between 2014 and 2021. Ten neonates were excluded from analysis as they were scanned with older acquisition protocols. One neonate was excluded from analysis due to an acute infarction/parenchymal haemorrhage. 25 neonates with DS [12 female, 10 preterm births < 37 weeks GA, PMA at scan median (range) = 40.57 (32.43 – 45.57) weeks] (Table S1 and Table 2) were scanned using the same acquisition parameters as controls and were used for analysis. Neonates with clinical comorbidities (as listed in Table S2 and S3) were not excluded.

**Table 1:** Neonatal brain segmentation and relative volume calculation. Each segment of interest is accompanied by a description of tissue segments & structures included, and where applicable, how a relative volume was calculated. The table is categorised into **A**) whole brain volumes, **B**) total GM or WM volumes, **C**) regional volumes, **D.1**) specific tissue volumes, and **D.2**) specific CSF-filled volumes. Left and right brain regions were consolidated for all labels. *[Abbreviations: ICV = intracranial volume, TBV = total brain volume, TTV = total tissue volume, eCSF = extracerebral cerebrospinal fluid, GM = grey matter, WM = white matter.]*

|  | Description of tissue segments & structures included | Relative volume |
| --- | --- | --- |
| **<u>A) Whole Brain Volumes</u>** |  |  |
| Intracranial Volume (ICV) | All brain segments, excluding extracranial background. | - |
| Total Brain Volume (TBV) | All brain segments, excluding extracranial background and eCSF. | - |
| Total Tissue Volume (TTV) | All brain segments, excluding extracranial background, eCSF and lateral ventricles. | - |
| **<u>B) Total GM or WM Volumes</u>** |  |  |
| Total Cortical GM | Frontal Lobe GM, Temporal Lobe GM, Parietal Lobe GM, Occipital Lobe GM, Insula GM, Cingulate GM. | /TTV |
| Total Deep GM | Caudate Nucleus, Lentiform Nucleus, Thalamus, and Intracranial Background. | /TTV |
| Total WM | Frontal Lobe WM, Temporal Lobe WM, Parietal Lobe WM, Occipital Lobe WM, Insula WM, Cingulate WM. | /TTV |
| **<u>C) Regional Volumes</u>** |  |  |
| Total Frontal Lobe | Frontal Lobe GM and WM | /TTV |
| Total Temporal Lobe | Temporal Lobe GM and WM | /TTV |
| Total Parietal Lobe | Parietal Lobe GM and WM | /TTV |
| Total Occipital Lobe | Occipital Lobe GM and WM | /TTV |
| Total Insula | Insula GM and WM | /TTV |
| Total Cingulate | Cingulate GM and WM | /TTV |
| Posterior Fossa | Cerebellum and Brainstem | /TTV |
| Basal Ganglia | Caudate Nucleus and Lentiform Nucleus | /TTV |
| **<u>D.1) Specific Tissue Volumes</u>** |  |  |
|  | Frontal Lobe GM, Frontal Lobe WM, Temporal Lobe GM, Temporal Lobe WM, Parietal Lobe GM, Parietal Lobe WM, Occipital Lobe GM, Occipital Lobe WM, Insula GM, Insula WM, Cingulate GM, Cingulate WM, Cerebellum, Brainstem, Caudate Nucleus, Lentiform Nucleus, Thalamus, Hippocampus, Amygdala. | /TTV |
| **<u>D.2) Specific CSF-Filled Volumes</u>** |  |  |
| eCSF | Extracerebral cerebrospinal fluid (eCSF), including the third and fourth ventricles. | /ICV |
| Lateral Ventricles | Lateral ventricles, including cavum septum pellucidum (if present). | /TBV |

**Table 2:**
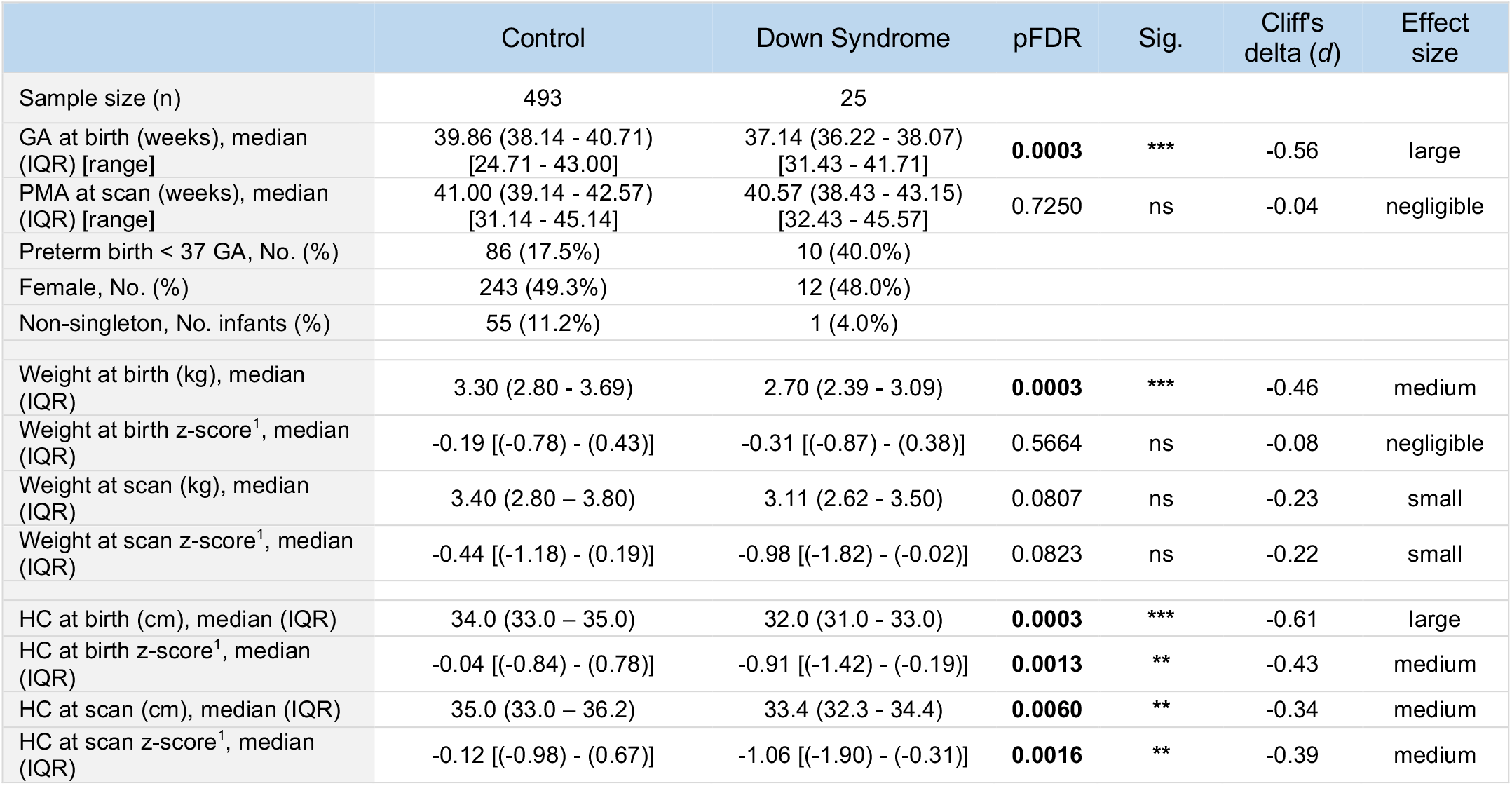
Summary of demographic, weight, and head circumference data. Details of GA at birth, PMA at scan, number of preterm births (< 37 weeks GA), sex and non-singleton neonates (e.g., twins) are provided for both control (n = 493) and Down syndrome (n = 25) groups. Non-parametric Mann-Whitney U tests with Benjamini & Hochberg’s FDR multiple comparison correction (pFDR) were conducted between the two groups. Cliff’s delta (*d*) test was used to assess effect size. pFDR < 0.05 are bolded. (^1^) indicates z-scores calculated using the RCPCH UK-WHO growth charts and not GPR modelling. *[Abbreviations: GA = gestational age at birth, HC = head circumference, IQR = inter-quartile range, PMA = postmenstrual age at scan.]*

#### Preterm- and term-born neonates for normative modelling

A total of 493 preterm- and term-born neonates [243 female, 86 preterm births < 37 weeks GA, of which 33 were born < 32 GA, PMA at scan median (range) = 41.00 (31.14 - 45.14) weeks] were selected from the dHCP (www.developingconnectome.org)^45^ to generate a normative model from 32 to 46 weeks PMA at scan^46–48^ (Table 2). Exclusion criteria included incidental findings on MRI^49^ (detailed in section *MR image review*, below) and Bayley III Scales of Infant Development (BSID-III)^50^ cognitive and motor composite scores (test mean [SD] = 100 [15]) below 70 (> 2 SD below the test mean) at 18 months^48^. No repeat scans from the same neonate were used. Healthy neonates from twin pregnancies were included. Taking all above exclusion criteria into account, this preterm- to term-born reference sample was referred to as a *‘control cohort’* for the purpose of this specific study.

### Clinical information

Weight, head circumference (HC), and clinical details were extracted from clinical notes at the time of scan. Weights (at birth and scan, in kg) and HC (at birth and scan, in cm) were converted into z-scores based on the RCPCH UK-WHO growth charts^51^ using the *‘childsds’* package v0.7.6 in R (Table S1 and Table 2). For neonates with DS, CHD diagnosis (n = 13, 52% of DS group) and additional clinical details, such as gastrointestinal malformations, can be found respectively in Table S2 and S3.

### MRI acquisition and pre-processing

Neonatal MRI data were acquired on a Philips Achieva 3T system using a dedicated 32-channel neonatal head coil and positioning system^52^. Imaging was performed during natural sleep without sedation^52^. T2-weighted scans were acquired with repetition time (TR) = 12000 ms, echo time (TE) = 156 ms, flip angle = 90°, SENSE factor = 2.11 / 2.58 (axial / sagittal). The resultant in-plane resolution was 0.8 x 0.8 mm with a slice thickness of 1.6 mm and a slice overlap of 0.8 mm. Images were motion-corrected^53^ and super-resolution reconstructed^54^ resulting in a 0.5 mm isotropic pixel resolution.

### MR image review

All MRI scans were examined by a neonatal neuroradiologist. Exclusion criteria for the dHCP scans were incidental findings with possible or likely significance for clinical outcome and/or imaging analysis, including acute infarction or parenchymal haemorrhage, and major lesions within white matter, cortical grey matter, cerebellum, or basal ganglia^49^. However, we did not exclude neonates with < 10 punctate white matter lesions (PWML), small subependymal cysts, small subdural haemorrhages, or haemorrhages in the caudothalamic notch, as these are common findings, especially within the preterm population^49^. One DS scan was excluded from analysis due to an acute infarction/parenchymal haemorrhage.

### MR image segmentation

Motion-corrected and reconstructed T2-weighted images^54^ were corrected for bias-field inhomogeneities, brain extracted and segmented using the dHCP structural pipeline (https://github.com/BioMedIA/dhcp-structural-pipeline; Accessed 15 November 2021)^55^, an automated tissue structure segmentation algorithm optimised for the neonatal brain^55–57^. T2-weighted images were segmented into seven main tissue classes: the extracerebral cerebrospinal fluid (eCSF), lateral ventricles, cortical GM, WM, deep GM, cerebellum, and brainstem. The eCSF included the third and fourth ventricles but excluded the lateral ventricles. The lateral ventricles included the *cavum septum pellucidum*, a transient fluid-filled cavity located in the midline of the brain, between the left and right anterior horns, which, if present, typically closes in the neonatal period^58,59^. Cortical GM, deep GM and WM were further automatically segmented into the specific tissue segments summarised in Table 1^56,57^. Tissue segments were visually inspected for accuracy, and where appropriate, any mislabelled voxels were manually corrected using ITK-SNAP (version 3.8.0)^60^. Tissue segments were used to calculate absolute (cm^3^) and relative (proportion) volumes. Relative volumes were calculated as the proportion of each tissue volume over total tissue volume (TTV), except for the lateral ventricles, which were calculated as a proportion of total brain volume (TBV), and eCSF as a proportion of intracranial volume (ICV) (as defined in Table 1).

### Normative modelling using Gaussian process regression

Gaussian process regression (GPR) was used to model absolute and relative volumetric development for each tissue segment (Table 1) based on cross-sectional data from 493 preterm- to term-born control neonates, whilst accounting for sex, PMA at scan (in weeks) and age from birth (in weeks) variables. GPR modelling was implemented using GPy in Python (https://sheffieldml.github.io/GPy/; Accessed 15 November 2021). GPR is a

Bayesian non-parametric regression method that provides a point estimate of the average volume and measures of predictive confidence for every observation, whilst accounting for modelled covariates^61^. The difference between predicted and observed values normalised by the predictive confidence (standard deviation, SD) represents the deviation of a data point from the expected mean, expressed as a z-score in units of SD^47,48,61,62^. Absolute and relative volumetric z-scores, for each tissue segment, were extracted from GPR modelling for an independent sample of 25 neonates with DS.

### Statistical analyses

Absolute and relative volumetric z-scores were used in statistical analyses. Shapiro-Wilk and Kolmogorov-Smirnov tests were used to test normality. In general, non-parametric tests such as the Mann-Whitney U test and Kruskal-Wallis one-way test of variance were used to test statistical difference between groups, including DS vs control, or DS neonates with vs without CHD. Cliff’s delta (*d*, ranging from −1 to 1) was used to assess effect size using the *‘effsize’* package v 0.8.1 in R. Effect sizes were categorised as negligible (< ± 0.147), small (< ± 0.33), medium (< ± 0.474), or large (≥ ± 0.474)^63,64^. Extreme deviations in volume were taken as a z-score ≤ - 2.6 or ≥ + 2.6 SD, representing the top and bottom 0.5% of the control population^47^. Simple linear regressions were used to model the relationship between volumetric z-scores and PMA at scan (in weeks). Spearman’s rank correlation coefficient (Rho, *ρ*) was used to assess correlation, which was considered very weak from 0 < *ρ ≤* 0.19, weak from 0.20 *≤ ρ ≤* 0.39, moderate from 0.40 *≤ ρ ≤* 0.59, strong from 0.60 *≤ ρ ≤* 0.79 and very strong from 0.80 *≤ ρ ≤* 1.00. R^2^ (the coefficient of determination) was used to indicate goodness of linear model fit. The extra sum-of-squares F test (GraphPad Prism v9.1.1.) was used to test for differences in the slope or intercept (i.e., elevation) parameters of two separate linear regressions (e.g., DS vs control, or DS neonates with vs without CHD). This test compares whether a combined model or two separate linear models provide a better goodness-of-fit. The result is expressed as an F ratio, from which a P value is calculated. For all above analyses, Benjamini & Hochberg’s False Discovery Rate was applied to correct for multiple comparisons (reported as pFDR) and statistical significance was set at pFDR < 0.05. For the extra sum-of-squares F test and the Spearman’s rank correlation test, the uncorrected P value was also presented alongside pFDR as additional information. All analyses and visualisations were performed in GraphPad Prism v9.1.1. or R v4.1.0 and 3D brain visualisations were created using HCP workbench^65^ and/or ITKSNAP^60^.

## RESULTS

### i) Demographic characteristics of participant groups

Demographics of the DS (n = 25, 48.0% female) and control groups (n = 493, 49.3% female) are summarised in Table 2. PMA at scan was not significantly different between DS and control groups (pFDR = 0.73, *d* = −0.04). However, GA at birth was significantly earlier in the DS group (pFDR = 0.0003, *d* = −0.56).

As a cohort, neonates with DS weighed less at birth (median = 2.70 kg, *d* = −0.46, pFDR = 0.0003). However, after correcting for sex and age differences using z-scores derived from the RCPCH UK-WHO growth charts^51^, DS birth weight (median z-score = −0.31 SD, *d* = −0.08, pFDR = 0.57) and scan weight (median z-score = −0.98 SD, *d* = −0.22, pFDR = 0.082) were not significantly different from control. Neonates with DS had a smaller absolute head circumference (HC) at birth (median = 32.0 cm, *d* = −0.61, pFDR = 0.0003) and at scan (median = 33.4 cm, *d* = −0.34, pFDR = 0.006), which remained smaller than control even after correcting for sex and age differences using z-scores (HC median z-score at birth = −0.91 SD, *d* = −0.43, pFDR = 0.0013; HC median z-score at scan = −1.06 SD, *d* = - 0.39, pFDR = 0.0016) (Table 2).

### ii) Whole brain volumes were significantly smaller in neonates with DS, as were most underlying tissue volumes

Absolute volumetric z-scores for neonates with DS were extracted from GPR normative modelling (Figures 2, 3 and S1) and used to compute groupwise comparisons by tissue segment (Table 3 and Figure 4).

**Figure 2:**
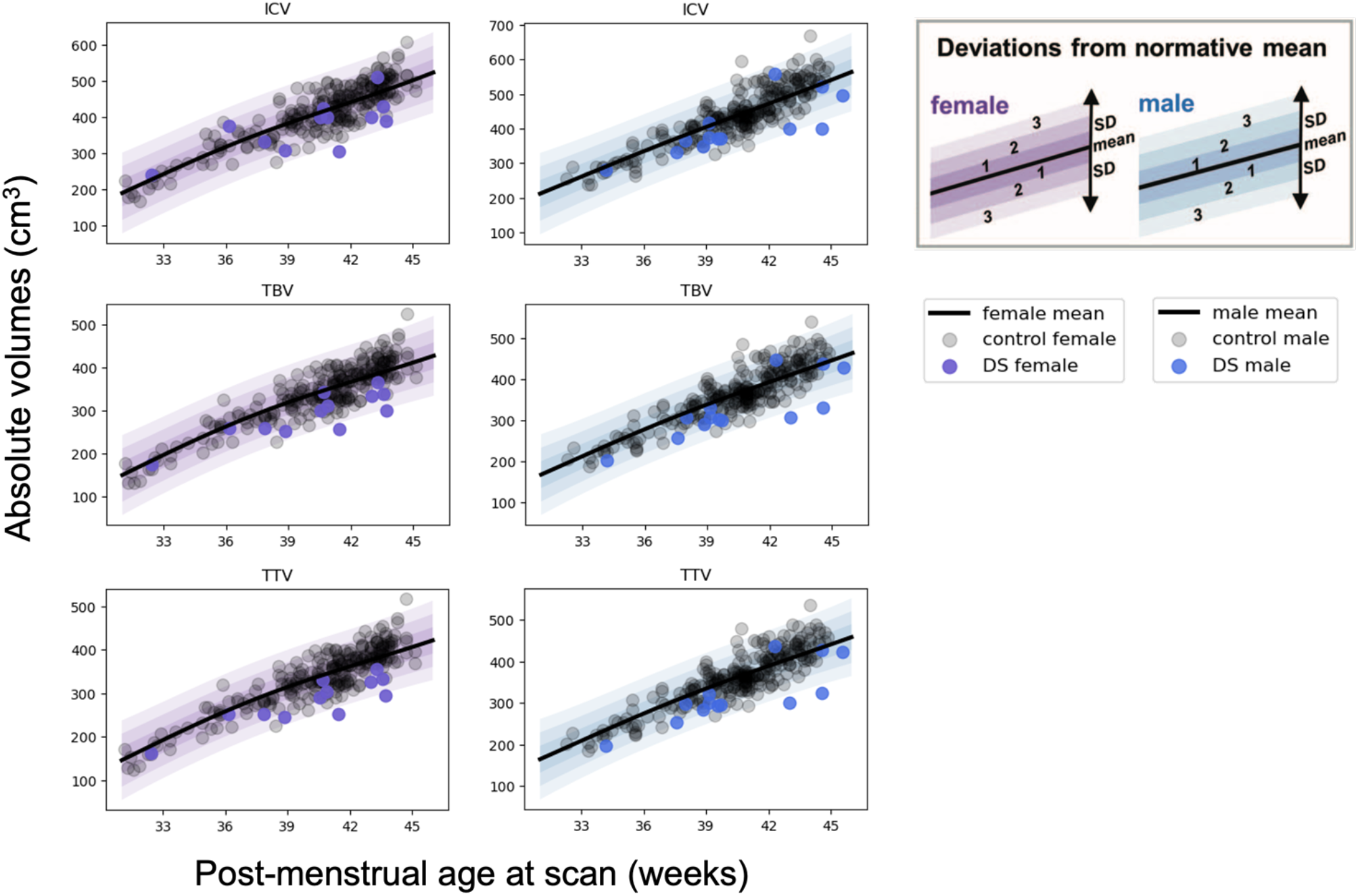
Normative modelling of absolute whole brain volumes from 32 to 46 weeks PMA. GPR plots of absolute whole brain volumes for females (in purple) and males (in blue) from 32 to 46 weeks PMA. Detailed descriptions of intracranial volume (ICV), total brain volume (TBV), and total tissue volume (TTV) can be found in Table 1. The normative mean appears as a bolded black curve, whilst shaded areas represent ±1, 2 and 3 standard deviations (SD) from the normative mean. Transparent grey dots represent control neonates (n = 243 females and n = 250 males). Data for the DS cohort (n = 25) is shown for females (purple dots, n = 12) and males (blue dots, n = 13). GPR plots for all specific tissue segments can be found in Figure S1.

**Figure 3:**
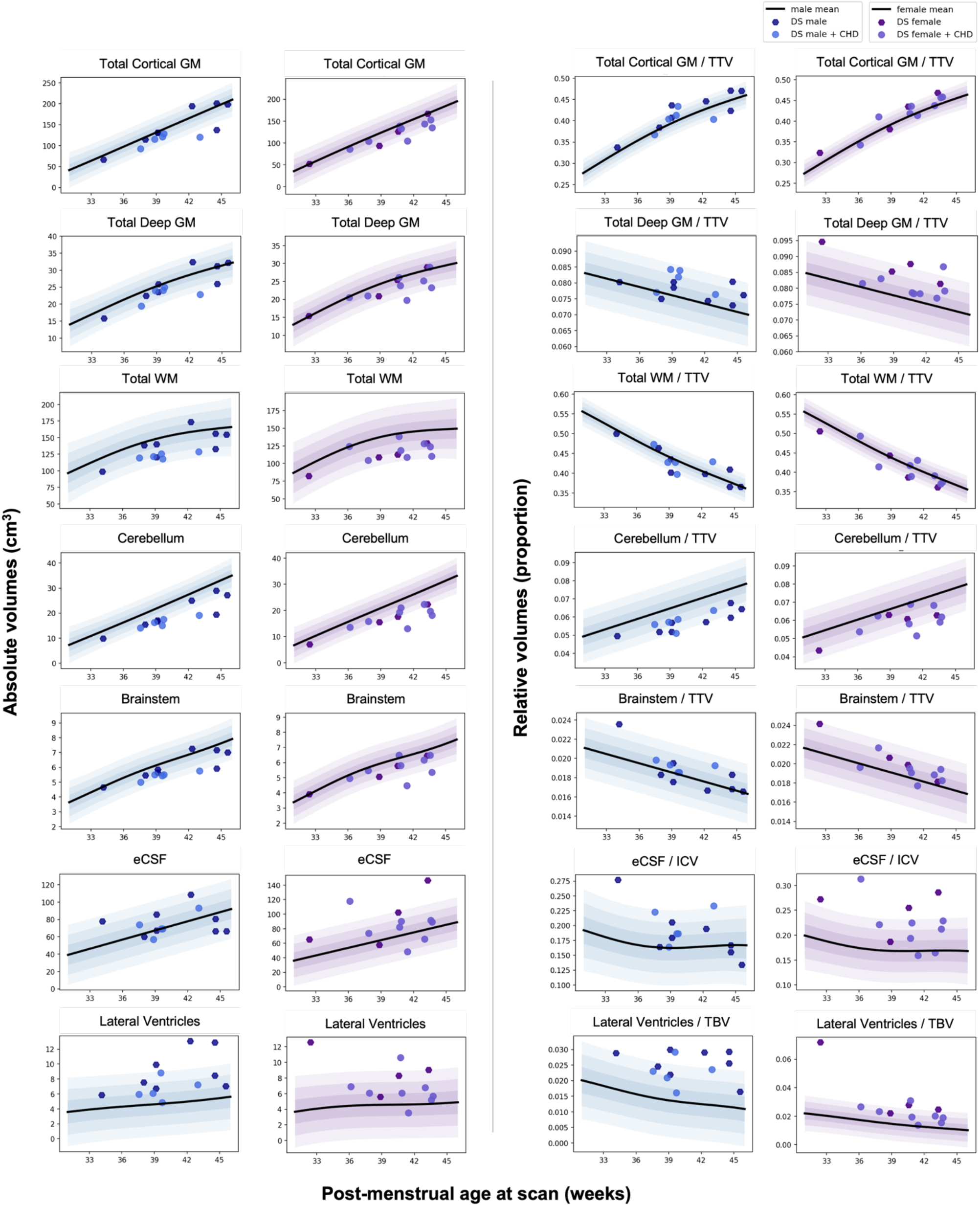
Normative modelling of absolute and relative main tissue volumes from 32 to 46 weeks PMA. GPR plots of the main tissue classes of the brain from 32 to 46 weeks PMA. Plots for *absolute volumes* (in cm^3^) appear on the left side and plots for *relative volumes* (i.e., proportion of TTV, TBV or ICV) appear on the right side. The normative mean appears as a bolded black curve, whilst shaded areas represent ±1, 2 and 3 standard deviations from the normative mean. Control neonates are not shown for ease of view. Data for DS neonates (n = 25) is shown for females (purple dots, n = 12) and males (blue dots, n = 13). Lighter dots for both females and males indicate DS neonates with a congenital heart defect (CHD) (n = 5 males, 8 females). GPR plots for all specific tissue segments can be found in Figure S1.

**Figure 4:**
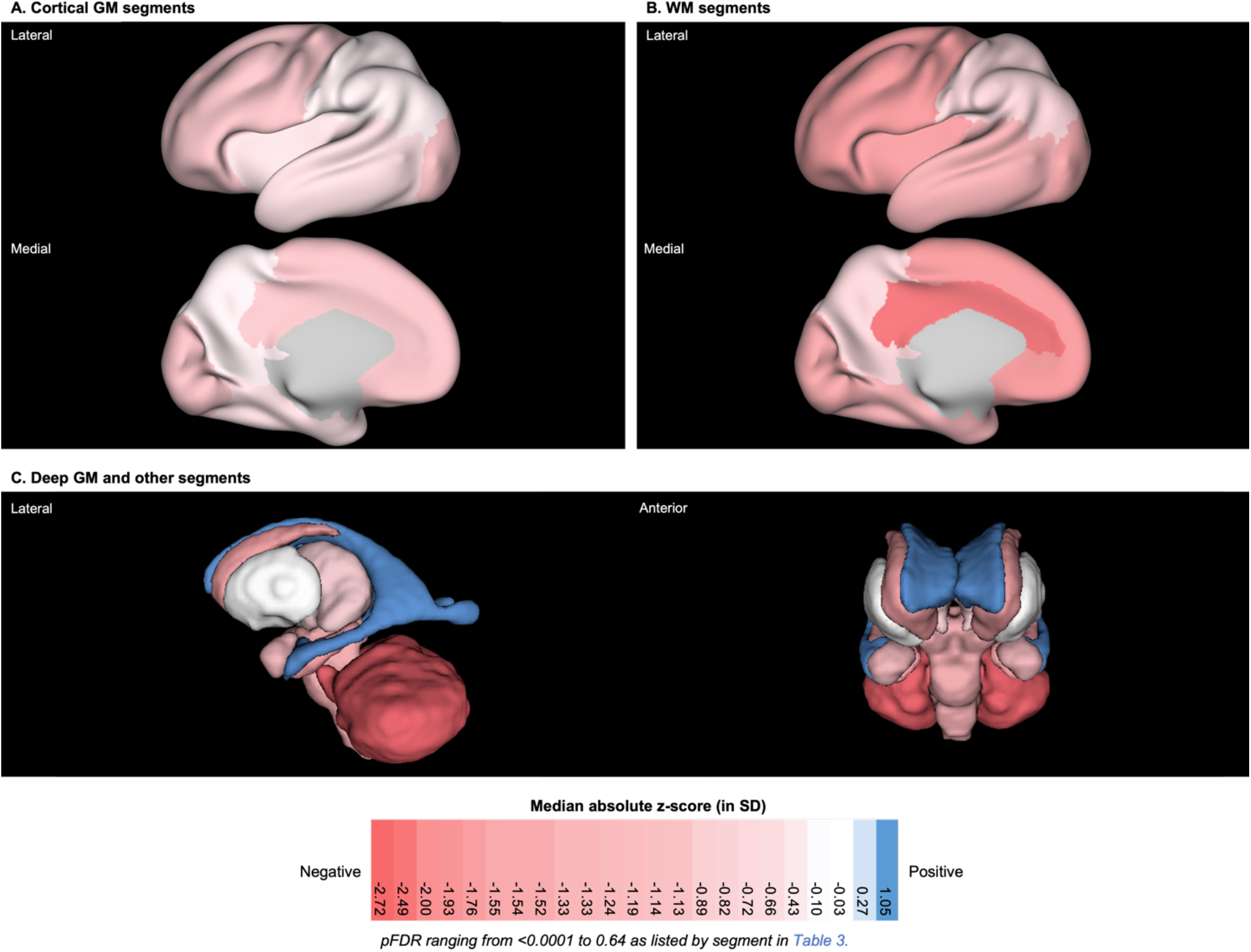
3D brain visualisation of the median absolute z-score by tissue segment for neonates with DS. 3D brain visualisation indicating the median absolute z-score (in standard deviations, SD) by tissue segment for neonates with DS. pFDR values can be found listed in Table 3. The median z-score is indicated by a colour scale, whereby red indicates a negative deviation from the normative mean (z < 0, i.e., a smaller volume than control), white indicates no significant deviation (z ~ 0), and blue indicates a positive deviation (z > 0, i.e., a larger volume than control). 3D brain visualisation for **A**) cortical GM segments, **B**) WM segments and **C**) Deep GM and other segments.

**Table 3:** Groupwise comparison (DS vs Control) of absolute volumetric z-scores by tissue segment. Table of median z-scores and interquartile range (IQR) derived from *absolute volumes* (see Figure S1) for DS (n = 25) and control (n = 493) groups. The table is organised into **A**) whole brain volumes, **B**) total GM or WM volumes, **C**) regional volumes and **D**) specific tissue volumes (including CSF-filled volumes). A non-parametric Kruskal-Wallis test with Benjamini & Hochberg’s FDR multiple comparison correction (pFDR) was performed for each tissue segment. Cliff’s delta (*d*) was used to assess the effect size. A colour scale has been applied, whereby red indicates a negative deviation from the normative mean (z < 0, i.e., a smaller volume than control), white indicates no significant deviation (z ~ 0), and blue indicates a positive deviation (z > 0, i.e., a larger volume than control).

|  | DS<br>median<br>(z-score) | DS IQR<br>(z-score) | Control<br>median<br>(z-score) | Control<br>IQR<br>(z-score) | Kruskal<br>Wallis<br>(pFDR) | Sig. | Cliff's<br>delta | Effect size |
| --- | --- | --- | --- | --- | --- | --- | --- | --- |
| **A) Whole Brain Volumes** |  |  |  |  |  |  |  |  |
| ICV | -1.05 | -2.21, -0.34 | -0.01 | -0.67, 0.55 | <0.0001 | **** | -0.50 | large |
| TBV | -1.71 | -2.27, -0.76 | -0.05 | -0.69, 0.58 | <0.0001 | **** | -0.72 | large |
| TTV | -1.76 | -2.34, -0.95 | -0.03 | -0.67, 0.58 | <0.0001 | **** | -0.76 | large |
| **B) Total GM or WM Volumes** |  |  |  |  |  |  |  |  |
| Total White Matter | -1.91 | -2.56, -1.07 | -0.03 | -0.67, 0.59 | <0.0001 | **** | -0.80 | large |
| Cortical Grey Matter | -1.22 | -2.07, -0.50 | -0.05 | -0.61, 0.58 | <0.0001 | **** | -0.59 | large |
| Deep Grey Matter | -1.04 | -1.86, -0.47 | 0.03 | -0.68, 0.58 | <0.0001 | **** | -0.57 | large |
| **C) Regional Volumes** |  |  |  |  |  |  |  |  |
| Posterior Fossa | -2.43 | -3.26, -1.54 | -0.05 | -0.67, 0.67 | <0.0001 | **** | -0.93 | large |
| Total Cingulate | -1.98 | -2.89, -1.35 | -0.06 | -0.64, 0.58 | <0.0001 | **** | -0.83 | large |
| Total Frontal Lobe | -1.92 | -2.67, -1.01 | -0.05 | -0.66, 0.57 | <0.0001 | **** | -0.81 | large |
| Total Insula | -1.62 | -3.01, -0.91 | -0.02 | -0.66, 0.67 | <0.0001 | **** | -0.79 | large |
| Total Occipital Lobe | -1.52 | -2.04, -0.98 | -0.02 | -0.66, 0.62 | <0.0001 | **** | -0.78 | large |
| Total Temporal Lobe | -1.08 | -2.22, -0.40 | -0.07 | -0.67, 0.59 | <0.0001 | **** | -0.57 | large |
| Total Parietal Lobe | -0.65 | -1.73, 0.00 | -0.02 | -0.71, 0.61 | 0.0005 | *** | -0.41 | medium |
| Basal Ganglia | -0.56 | -1.37, -0.31 | -0.02 | -0.67, 0.62 | 0.0003 | *** | -0.43 | medium |
| **D) Specific Tissue Volumes** |  |  |  |  |  |  |  |  |
| Cingulate WM | -2.72 | -3.46, -1.89 | -0.05 | -0.62, 0.59 | <0.0001 | **** | -0.95 | large |
| Cerebellum | -2.49 | -3.51, -1.65 | -0.04 | -0.64, 0.65 | <0.0001 | **** | -0.95 | large |
| Frontal Lobe WM | -2.00 | -2.72, -1.24 | -0.04 | -0.66, 0.62 | <0.0001 | **** | -0.86 | large |
| Insula WM | -1.93 | -2.75, -1.20 | -0.02 | -0.68, 0.65 | <0.0001 | **** | -0.87 | large |
| Occipital Lobe WM | -1.76 | -1.96, -1.14 | -0.03 | -0.76, 0.61 | <0.0001 | **** | -0.79 | large |
| Temporal Lobe WM | -1.55 | -2.41, -0.69 | -0.03 | -0.66, 0.58 | <0.0001 | **** | -0.68 | large |
| Caudate Nucleus | -1.54 | -2.00, -0.69 | -0.09 | -0.69, 0.57 | <0.0001 | **** | -0.71 | large |
| Hippocampus | -1.52 | -2.46, -0.65 | -0.11 | -0.7, 0.56 | <0.0001 | **** | -0.66 | large |
| Intra Cranial Background | -1.33 | -2.17, -0.66 | -0.03 | -0.68, 0.64 | <0.0001 | **** | -0.54 | large |
| Brainstem | -1.33 | -1.95, -0.30 | -0.02 | -0.65, 0.60 | <0.0001 | **** | -0.62 | large |
| Occipital Lobe GM | -1.24 | -1.93, -0.67 | -0.03 | -0.64, 0.59 | <0.0001 | **** | -0.68 | large |
| Frontal Lobe GM | -1.19 | -2.06, -0.61 | -0.13 | -0.61, 0.58 | <0.0001 | **** | -0.68 | large |
| Amygdala | -1.14 | -2.23, -0.57 | -0.02 | -0.71, 0.64 | <0.0001 | **** | -0.59 | large |
| Cingulate GM | -1.13 | -2.21, -0.43 | -0.07 | -0.64, 0.59 | <0.0001 | **** | -0.54 | large |
| Parietal Lobe WM | -0.89 | -1.59, -0.27 | -0.05 | -0.70, 0.65 | 0.0001 | *** | -0.46 | medium |
| Thalamus | -0.82 | -1.85, -0.48 | -0.02 | -0.63, 0.66 | <0.0001 | **** | -0.53 | large |
| Temporal Lobe GM | -0.72 | -2.14, 0.15 | -0.06 | -0.67, 0.57 | 0.0003 | *** | -0.44 | medium |
| Insula GM | -0.66 | -2.05, -0.07 | -0.07 | -0.66, 0.66 | 0.0002 | *** | -0.44 | medium |
| Parietal Lobe GM | -0.43 | -1.92, 0.00 | -0.01 | -0.64, 0.56 | 0.0039 | ** | -0.35 | medium |
| Lentiform Nucleus | -0.03 | -0.72, 0.57 | -0.05 | -0.64, 0.61 | 0.6425 | ns | -0.05 | negligible |
| eCSF | 0.27 | -0.91, 1.78 | -0.02 | -0.72, 0.60 | 0.1977 | ns | 0.16 | small |
| Lateral Ventricles | 1.05 | -0.06, 2.54 | -0.18 | -0.71, 0.52 | <0.0001 | **** | 0.52 | large |

Absolute whole brain volumes were significantly smaller in neonates with DS compared to control with large effect sizes (intracranial volume (ICV) median z-score = −1.05 SD, *d* = −0.50; total tissue volume (TTV) median z-score = −1.76 SD, *d* = −0.76; pFDR < 0.0001) (Table 3.A). As whole brain volumes were significantly smaller in DS, most underlying tissue segments were also significantly smaller than control in absolute volume (Table 3.D). This was clearly evidenced by the large number of segments in red (indicating a median z < 0, i.e., a smaller volume than control) listed in Table 3.D and seen visually in Figure 4. Only the lentiform nuclei (comprising the putamen and pallidum) (median z-score = −0.03 SD, *d* = −0.05, ns) and the eCSF (median z-score = 0.27, *d* = +0.16, ns) were not significantly different from control, whilst the lateral ventricles were significantly enlarged (median z-score = +1.05 SD, *d* = +0.52, pFDR < 0.0001).

Of particular interest, we observed that the total cerebral WM (median z-score = −1.91 SD, *d* = −0.80, pFDR < 0.0001) was smaller compared to control than total cortical GM (median z-score = −1.22 SD, *d* = −0.59, pFDR < 0.0001) (Table 3.B). Lastly, from a regional perspective, the parietal lobe (median z-score = −0.65 SD, *d* = −0.41, pFDR = 0.0005), and the basal ganglia (comprising the caudate and lentiform nuclei) (median z-score = −0.56 SD, *d* = −0.43, pFDR = 0.0003) were less deviated from the normative mean than other regions in absolute volume (Table 3.C).

### iii) Relative volumes demonstrated a dynamic shift in tissue proportions across the brain in neonates with DS

Although, most tissue segments were significantly smaller than control in absolute volume (Table 3), the use of relative volumes, adjusting for differences in whole brain volume (as per Table 1), revealed that there was a dynamic shift in tissue proportions across the brain in neonates with DS compared to controls. This scaling exercise^66^ showed that some relative tissue volumes were not significantly different in proportion from control, with small to negligible effect sizes (i.e., approximately *isometric*), whilst other segments were significantly smaller or larger in proportion with medium to large effect sizes (i.e., *allometric*) as indicated by a colour scale in Table 4 and visualised in Figure 5.

**Table 4.** Groupwise comparison (DS vs Control) of relative volumetric z-scores by tissue segment. Table of median z-scores and interquartile range (IQR) derived from *relative volumes* (see Figure S1) for DS (n = 25) and control (n = 493) groups. The table is organised into **A**) total GM or WM volumes, **B**) regional volumes and **C**) specific tissue volumes (including CSF-filled volumes). A non-parametric Kruskal-Wallis test with Benjamini & Hochberg’s FDR multiple comparison correction (pFDR) was performed for each tissue label. Cliff’s delta (*d*) tests were used to assess the effect size. A colour scale has been applied, whereby red indicates a negative deviation from the normative mean (z < 0, ‘allometric smaller’, i.e., a smaller proportion of the whole brain than control), white indicates no significant deviation (z ~ 0, approximately ‘isometric’), and blue indicates a positive deviation (z > 0, ‘allometric larger’, i.e., a larger proportion of the whole brain than control).

|  | DS<br>median<br>(z-score) | DS IQR<br>(z-score) | Control<br>median<br>(z-score) | Control<br>IQR<br>(z-score) | Kruskal<br>Wallis<br>(pFDR) | Sig. | Cliff's<br>delta | Effect<br>size |
| --- | --- | --- | --- | --- | --- | --- | --- | --- |
| **A) Total GM or WM Volumes** |  |  |  |  |  |  |  |  |
| Total WM / TTV | -0.59 | -2.30, 0.54 | -0.02 | -0.59, 0.57 | 0.0119 | * | -0.32 | small |
| Total Deep GM / TTV | 1.27 | 0.56, 2.31 | 0.03 | -0.66, 0.59 | <0.0001 | **** | 0.65 | large |
| Total Cortical GM / TTV | 1.37 | -0.05, 2.40 | -0.06 | -0.64, 0.55 | 0.0001 | *** | 0.46 | medium |
| **B) Regional Volumes** |  |  |  |  |  |  |  |  |
| Total Cingulate / TTV | -1.57 | -2.04, -0.82 | -0.08 | -0.71, 0.69 | <0.0001 | **** | -0.66 | large |
| Posterior Fossa / TTV | -1.31 | -2.40, -1.06 | 0.04 | -0.64, 0.61 | <0.0001 | **** | -0.76 | large |
| Total Frontal Lobe / TTV | -1.03 | -1.89, -0.10 | 0.01 | -0.69, 0.75 | <0.0001 | **** | -0.48 | large |
| Total Insula / TTV | -0.77 | -2.07, 0.19 | -0.02 | -0.69, 0.69 | 0.0033 | ** | -0.36 | medium |
| Total Occipital Lobe / TTV | -0.71 | -1.34, 0.09 | 0.00 | -0.64, 0.67 | 0.0018 | ** | -0.40 | medium |
| Total Temporal Lobe / TTV | 0.64 | -0.04, 2.04 | 0.03 | -0.65, 0.65 | 0.0008 | *** | 0.43 | medium |
| Basal Ganglia / TTV | 1.31 | 0.46, 1.87 | -0.01 | -0.66, 0.59 | <0.0001 | **** | 0.64 | large |
| Total Parietal Lobe / TTV | 2.03 | 1.25, 2.66 | -0.06 | -0.65, 0.75 | <0.0001 | **** | 0.86 | large |
| **C) Specific Tissue Volumes** |  |  |  |  |  |  |  |  |
| Cingulate WM / TTV | -2.52 | -3.02, -1.67 | -0.09 | -0.63, 0.60 | <0.0001 | **** | -0.94 | large |
| Cerebellum / TTV | -1.83 | -2.75, -1.34 | 0.02 | -0.63, 0.63 | <0.0001 | **** | -0.85 | large |
| Frontal Lobe WM / TTV | -1.65 | -2.55, 0.15 | 0.01 | -0.74, 0.65 | 0.0001 | *** | -0.48 | large |
| Insula WM / TTV | -1.57 | -2.60, -0.46 | 0.01 | -0.69, 0.62 | <0.0001 | **** | -0.60 | large |
| Occipital Lobe WM / TTV | -0.87 | -1.60, -0.42 | 0.05 | -0.66, 0.64 | <0.0001 | **** | -0.52 | large |
| Hippocampus / TTV | -0.41 | -1.19, 0.40 | -0.07 | -0.71, 0.68 | 0.1861 | ns | -0.10 | negligible |
| Occipital Lobe GM / TTV | -0.35 | -1.01, 0.81 | -0.04 | -0.65, 0.65 | 0.4247 | ns | -0.10 | negligible |
| Caudate Nucleus / TTV | -0.28 | -1.03, 0.25 | -0.08 | -0.63, 0.61 | 0.1198 | ns | -0.20 | small |
| Temporal Lobe WM / TTV | -0.28 | -0.85, 0.99 | 0.00 | -0.61, 0.60 | 0.9167 | ns | -0.02 | negligible |
| Cingulate GM / TTV | -0.07 | -0.72, 1.11 | -0.06 | -0.72, 0.69 | 0.8384 | ns | 0.03 | negligible |
| Frontal Lobe GM / TTV | 0.38 | -0.31, 0.84 | -0.02 | -0.66, 0.61 | 0.2984 | ns | 0.13 | negligible |
| Amygdala / TTV | 0.77 | -0.40, 1.36 | -0.04 | -0.66, 0.67 | 0.0543 | ns | 0.23 | small |
| Insula GM / TTV | 0.85 | -0.23, 1.68 | 0.02 | -0.64, 0.67 | 0.0092 | ** | 0.33 | small |
| Brainstem / TTV | 0.88 | 0.18, 1.58 | -0.01 | -0.66, 0.61 | 0.0001 | *** | 0.43 | medium |
| eCSF / ICV | 1.11 | -0.08, 2.97 | 0.00 | -0.62, 0.56 | <0.0001 | **** | 0.51 | large |
| Parietal Lobe WM / TTV | 1.48 | 0.57, 1.92 | -0.05 | -0.63, 0.60 | <0.0001 | **** | 0.64 | large |
| Thalamus / TTV | 1.50 | 0.42, 2.30 | 0.00 | -0.67, 0.65 | <0.0001 | **** | 0.59 | large |
| Temporal Lobe GM / TTV | 1.62 | 0.09, 2.42 | 0.03 | -0.64, 0.65 | <0.0001 | **** | 0.52 | large |
| Parietal Lobe GM / TTV | 1.95 | 0.54, 2.42 | -0.11 | -0.65, 0.59 | <0.0001 | **** | 0.67 | large |
| Lateral Ventricles / TBV | 1.98 | 1.11, 3.03 | -0.19 | -0.68, 0.49 | <0.0001 | **** | 0.78 | large |
| Lentiform Nucleus / TTV | 2.23 | 1.42, 2.87 | -0.02 | -0.63, 0.62 | <0.0001 | **** | 0.82 | large |

**Figure 5:**
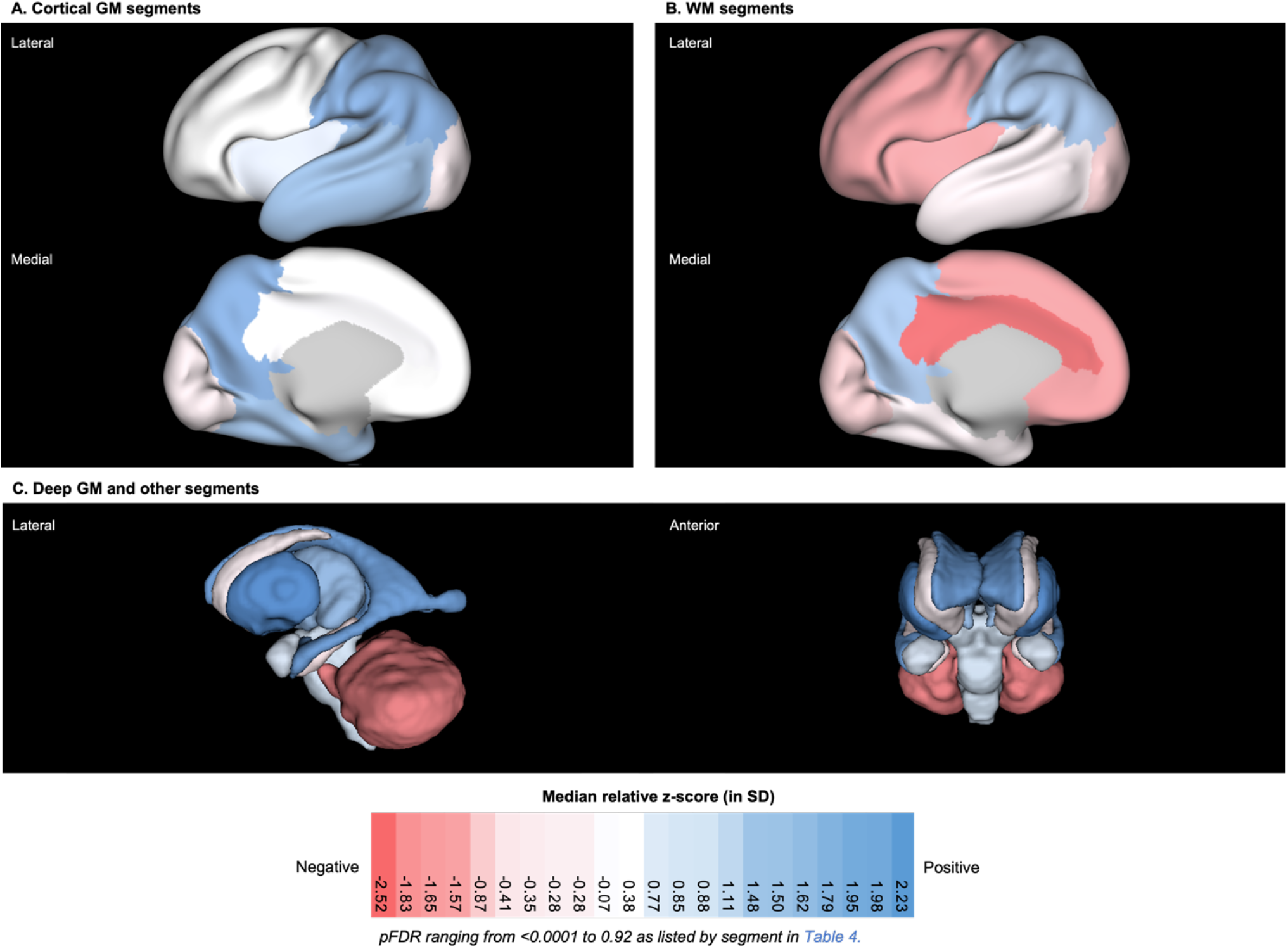
3D brain visualisation of the median relative z-score by tissue segment for neonates with DS. 3D brain visualisation indicating the median relative z-score (in standard deviations, SD) by tissue segment for neonates with DS. pFDR values can be found listed in Table 4. The median z-score is indicated by a colour scale, whereby red indicates a negative deviation from the normative mean (z < 0, ‘allometric smaller’, i.e., a smaller proportion of the whole brain than control), white indicates no significant deviation (z ~ 0, approximately ‘isometric’), and blue indicates a positive deviation (z > 0, ‘allometric larger’, i.e., a larger proportion of the whole brain than control). 3D brain visualisations for **A**) cortical GM segments, **B**) WM segments and **C**) Deep GM and other segments.

Firstly, we observed that five tissue segments (i.e., the cerebellum, as well as the cingulate, frontal, insular and occipital WM segments) were significantly smaller in relative volume (pFDR < 0.0001, *d* = −0.52 to −0.94), and thus occupied smaller proportions of the whole brain in neonates with DS compared to control, as indicated by darker red tones in Table 4.C and in Figure 5.

Total cortical GM was significantly enlarged (median z-score = +1.37 SD, *d* = +0.46, pFDR = 0.0001), whilst conversely, total cerebral WM was significantly reduced in relative volume compared to control (median z-score = −0.59 SD, *d* = −0.32, pFDR = 0.012) (Table 4.A). We observed a dynamic shift in regional proportions of cortical GM and WM segments in neonates with DS compared to control. Cortical GM segments were approximately isometric (i.e., occipital, cingulate, frontal GM segments; *d* = −0.10 to +0.13, pFDR = 0.30 to 0.84) or significantly enlarged in proportion (i.e., insular, temporal, and parietal GM segments; *d* = +0.33 to +0.67, pFDR = 0.0092 to < 0.0001). Comparatively, most WM segments were significantly reduced in proportion (i.e., cingulate, frontal, insular and occipital WM segments; *d* = −0.94 to −0.52, pFDR < 0.0001), except the temporal WM, which was approximately isometric (median z-score = −0.28 SD, *d* = −0.02, pFDR = 0.92) and the parietal WM, which was significantly enlarged in proportion (median z-score = +1.48 SD, *d* = +0.64, pFDR < 0.0001) (Table 4.C). Thus, from a total lobar perspective (i.e., GM + WM), we noticed a regional pattern, whereby the temporal and parietal lobes were enlarged, whilst the fronto-occipital lobes were reduced in relative volume compared to control (Table 4.B).

Within the deep GM, the lentiform nuclei, which were not significantly different from control in absolute volume (Table 3.D), were disproportionately enlarged in relative volume (median z-score = +2.23 SD, *d =* +0.82, pFDR< 0.0001) (Table 4.C). The thalami were also proportionally larger (median z-score = +1.50 SD, *d =* +0.59, pFDR < 0.0001), whilst the caudate nuclei were approximately isometric (median z-score = −0.28 SD, *d =* −0.20, pFDR = 0.12). Separately, the hippocampus and the amygdala were both approximately isometric with small to negligible effect sizes (*d* = −0.10 to +0.23, pFDR = 0.054 to 0.19).

Within the posterior fossa, the cerebellum and the brainstem exhibited different dynamics. The cerebellum was significantly and disproportionately small with a very large effect size (median z-score = −1.83 SD, *d* = −0.85, pFDR < 0.0001), whilst conversely, the brainstem was significantly enlarged in relative volume with a medium effect size (median z-score = 0.88 SD, *d* = +0.43, pFDR = 0.0001).

Lastly, CSF-filled volumes were significantly enlarged in relative volume. The lateral ventricles were disproportionately enlarged with a very large effect size (median z-score = +1.98 SD, *d =* +0.78, pFDR < 0.0001). The eCSF, which was not significantly different from control in absolute volume (Table 3.D), was significantly enlarged in relative volume with a large effect size (median z-score = +1.11 SD, *d =* +0.51, pFDR < 0.0001) (Table 4.C).

### iv) Age-related deviation in volume from the control mean in neonates with DS

The neonatal period from 32 to 46 weeks PMA at scan was marked by a phase of rapid brain development. Absolute and relative volumetric development in the preterm- to term-born control cohort was characterised in detail and can be found as supplementary information in Figure S2 and Table S4. Simple linear regressions were used to appreciate age-related change in volumetric z-scores in neonates with DS, as well as age-related deviation from the control mean (Figures 6). Linear regressions for the control group were always characterised by a flat line at z = 0. Results for the extra sum-of-squares F tests are summarised in Table S5 and the Spearman’s rank correlation tests in Table S6.

**Figure 6:**
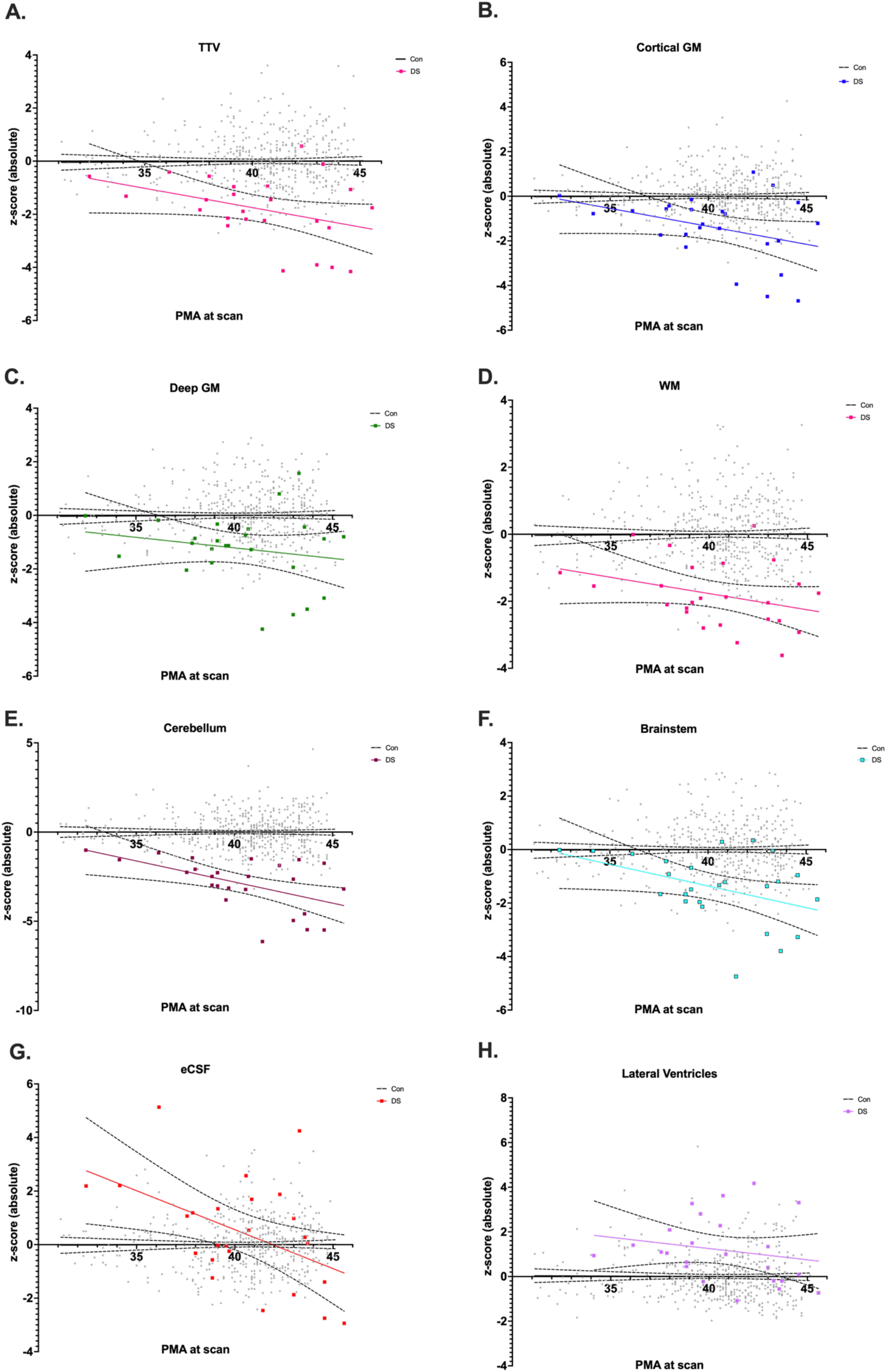
Simple linear regression of absolute z-scores against PMA at scan for whole brain and main tissue volumes. Simple linear regression plots of absolute z-scores against PMA at scan from 32 to 46 weeks for **A**. the whole brain (i.e., TTV) and main tissue classes of the brain, including **B**. the total cortical GM, **C**. the total deep GM, **D**. the total WM, **E**. the cerebellum, **F**. the brainstem, **G**. the eCSF and **H**. the lateral ventricles. Dots for individual control neonates (n = 493, females and males consolidated) appear in grey, and simple linear regressions appear as flat black lines at z = 0 with 95% confidence intervals as dotted lines. Dots for individual neonates with DS (n = 25, females and males) appear in colour, and the simple linear regressions appear as coloured lines with 95% confidence intervals as dotted lines. Linear regression plots for all other specific tissue segments can be found in Figure S3 and a table of results for F-tests in Table S5. Plots for z-scores derived from relative volumes can be found in Figure S5.

The linear regressions for whole brain volume (i.e., ICV and TTV) presented negative slopes that were significantly different from control (ICV, F ratio = 11.45, pFDR = 0.0021; TTV, F ratio = 5.42, pFDR = 0.034, Table S5), indicating a gradual deviation in whole brain volume from the control mean with advancing PMA at scan in neonates with DS. ICV was also moderately negatively correlated with advancing PMA at scan, although this correlation was no longer significant after multiple comparison correction (ICV, Spearman’s *ρ =* −0.52, R^2^ = 0.24, uncorrected P-value = 0.008, pFDR = 0.08, Table S6). As such, whole brain volumes were closer to the control mean for neonates born and scanned preterm (i.e., < 36 weeks PMA) but were markedly smaller than control for neonates scanned after term age (approximately 37 to < 46 weeks PMA) (Figures 6.A & S3). There was pronounced individual variability in neonates scanned at later ages, where there appeared to be a bimodal distribution of z-scores, whereby some neonates displayed typical whole brain volumes for DS, whilst others displayed extreme negative deviations (i.e., four neonates with TTV, z ≤ - 2.6 SD). The topic of individual variability in whole brain volume, particularly at later ages at scan, is covered again in section (vi) pertaining to neonates with CHD.

As per whole brain volume, most underlying absolute tissue volumes also displayed a gradual deviation from the control mean with advancing PMA (Figure 6.B-H, Table S5), including the total cortical GM (F ratio = 6.32, pFDR = 0.022), the brainstem (F ratio = 6.36, pFDR = 0.022), and the cerebellum (F ratio = 13.15, pFDR = 0.0009). The cerebellum (Figure 6.E) also displayed a moderate negative correlation with PMA (Spearman’s *ρ =* −0.54, R^2^ = 0.29, uncorrected P-value = 0.0058, pFDR = 0.08, Table S6), although this correlation was no longer significant after multiple comparison correction. It is worth noting that cerebellar deviation was particularly pronounced in neonates with DS, with twelve neonates displaying extreme negative deviations (z ≤ −2.6 SD).

The slopes for the total deep GM and total WM linear regressions (Figures 6.C & D) were not significantly different from control (deep GM, F ratio = 1.63, pFDR = 0.27; WM, F ratio = 2.52, pFDR = 0.16). However, their intercepts (i.e., *elevations*) were significantly different from control (deep GM, F ratio = 35.72, pFDR = 0.0003; WM, F ratio = 78.07, pFDR = 0.0003) (Table S5). Individual z-scores for total WM were more consistently below the control mean (z < 0) across all ages at scan.

The slope for eCSF (Figure 6.G) was significantly different from control (F ratio = 19.05, pFDR = 0.0003, Table S5). Individual z-scores for the eCSF were markedly larger than the control mean (i.e., +5.1 > z > +2 SD) for neonates born and scanned preterm, indicating a tendency for excessive eCSF at these ages. However, these gradually decreased by later ages at scan, although individual variability remained high across neonates.

Lastly, the lateral ventricles (Figure 6.H) tended to be consistently larger than the control mean at all ages. After removing an outlier with an extreme positive deviation (z = +5.8, scanned at 32.4 weeks PMA), possibly due to very early preterm birth, the slope for the lateral ventricles was not significantly different from control (F ratio = 1.61, pFDR = 0.21). However, the elevation was significantly different (F ratio = 29.40, pFDR = 0.0003) indicating the presence of consistently enlarged lateral ventricles for sex and age in neonates with DS (Table S5).

Linear regression plots for all specific tissue segments can be found for absolute z-scores (in Figure S3), and relative z-scores (in Figure S5). Results for the extra sum-of-squares F test are summarised in Tables S5 & S7, and the Spearman’s rank correlation test in Tables S6 & S8. Of note, the slope and intercept for the linear regression of the lentiform nuclei (Figure S3.D) were not different from control (slope, F ratio = 0.03, pFDR = 0.86; intercept, F ratio = 0.54, pFDR = 0.49). Thus, the lentiform nuclei represented the only brain segment in neonates with DS to develop entirely in line with control neonates.

Lastly, after correcting for differences in whole brain volume, most linear regressions no longer displayed a slope that was different from control using relative volumes (Figure S5 and Table S7). This is because the gradual deviation of whole brain z-score from the control mean with advancing PMA represented the main effect for most underlying tissue segments. However, the slope for relative eCSF (F ratio = 22.11, pFDR = 0.0003), relative cingulate GM (F ratio = 12.14, pFDR = 0.0014) and relative cingulate WM (F ratio = 6.85, pFDR = 0.02) remained significantly different from control.

### v) CHD+ and CHD-neonates did not show any statistically significant groupwise differences after multiple comparison correction

To assess the possible impact of CHD on neonatal brain volumes in DS, neonates were further categorised into subgroups with CHD (CHD+, n = 13) and without CHD (CHD-, n = 12) (Table S2). Overall, CHD+ and CHD-neonates did not show any statistically significant groupwise differences after FDR multiple comparison correction on a Kruskal-Wallis one-way test of variance (Table S9). This was most likely due to low statistical power, as subgroup sizes were small, and due to the large number of multiple comparisons.

However, certain underlying trends were observed from the uncorrected P-values (Table S9). In particular, the caudate nuclei (uncorrected P-value = 0.0356, pFDR = 0.25, d = −0.60, large effect) were smaller in CHD+ compared to CHD-neonates prior to multiple comparison correction. Furthermore, two GM segments, the temporal GM (uncorrected P-value = 0.0064, pFDR = 0.246, d = −0.59, large effect) and the parietal GM (uncorrected P-value = 0.0476, pFDR = 0.246, d = −0.49, large effect) were also smaller in CHD+ compared to CHD-neonates prior to multiple comparison correction.

### vi) Occipital WM volume was significantly reduced in CHD+ neonates compared to CHD-neonates with DS by later ages at scan

Absolute volumetric z-scores were plotted against PMA at scan for each tissue segment for CHD+ and CHD-neonates to observe age-related volumetric differences between subgroups. Linear regressions can be found in Figures 7 and S4, as well as results for the extra sum-of-squares F test (in Table S10) and the Spearman’s rank correlation test (in Table S11).

**Figure 7:**
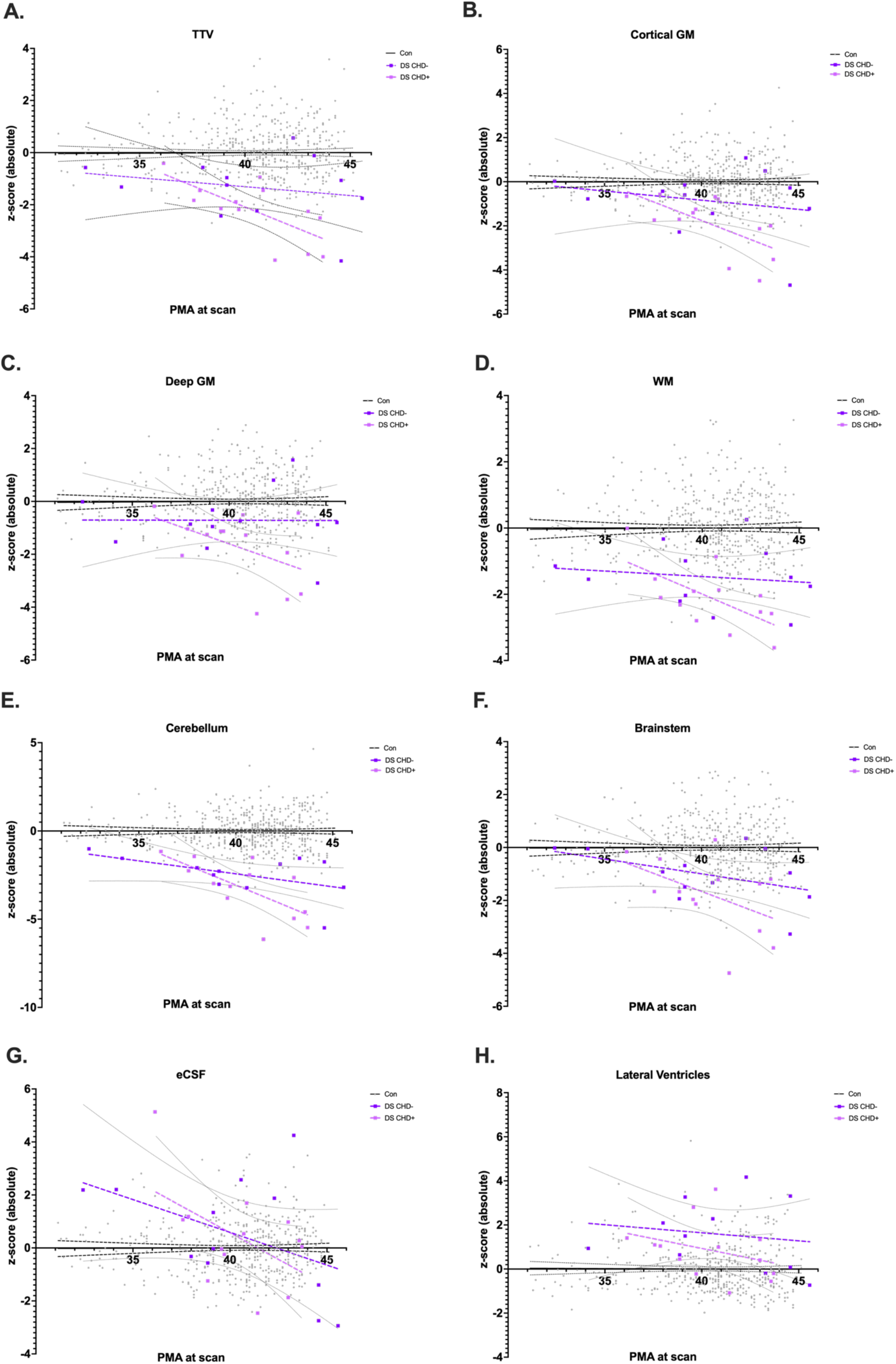
Simple linear regression of absolute z-scores against PMA at scan for whole brain and main tissue volumes in CHD+ and CHD-neonates with DS. Simple linear regression plots of absolute z-scores against PMA at scan (in weeks) appear as coloured lines with 95% confidence intervals as dotted lines for CHD+ (n = 13, in pink) and CHD-neonates (n = 12, in purple) with DS. Dots for individual control neonates (n = 493) appear in grey, and simple linear regressions appear as flat black lines at z = 0 with 95% confidence intervals as dotted lines. Plots for **A**. the whole brain (i.e., total tissue volume, TTV) and main tissue classes of the brain, including **B**. the total cortical GM, **C**. the total deep GM, **D**. the total WM, **E**. the cerebellum, **F**. the brainstem, **G**. the eCSF and **H**. the lateral ventricles. CHD+ and CHD-linear regression plots for all other specific tissue segments can be found in Figure S4 and a table of results for F-tests in Table S10 and Spearman’s correlations in Table S11.

Only one tissue segment, the occipital WM (Figure S4.C.), showed a statistically significant difference in slope after multiple comparison correction between CHD+ and CHD-neonates (F ratio = 18.21, uncorrected P-value = 0.0003, pFDR = 0.0084) (Table S10). In CHD+ neonates, the occipital WM segment also displayed a significant and very strong negative correlation with PMA (*ρ* = −0.89, R^2^ = 0.78, uncorrected P-value < 0.0001, pFDR = 0.0027), which was not the case for CHD-neonates (*ρ* = +0.15, R^2^ = 0.05, uncorrected P-value = 0.63, pFDR = 0.85) (Table S11). Thus, absolute occipital WM volume was significantly reduced in CHD+ neonates compared to CHD-neonates by later ages at scan (from approximately 40 weeks PMA).

Although several other tissue segments displayed differences between CHD+ and CHD-neonates on the F test and/or the Spearman’s test (uncorrected P-value < 0.05), these did not survive multiple comparison correction (pFDR < 0.05) (Tables S10 & S11). This was most likely due to low statistical power, as subgroup sizes were small, and due to the large number of multiple comparisons. However, we report certain underlying trends, which were observed from the uncorrected P-values for completeness of information, as this is a rare clinical cohort.

Firstly, whole brain volume in CHD+ neonates tended to be smaller than CHD-neonates by later ages at scan. ICV and TTV in CHD+ neonates displayed strong negative correlations with advancing PMA at scan (ICV, *ρ* = −0.67, R^2^ = 0.48, uncorrected P-value = 0.015, pFDR = 0.05; TTV, *ρ* = −0.73, R^2^ = 0.48, uncorrected P-value = 0.0059, pFDR = 0.05), which was not the case for CHD-neonates (e.g., TTV, *ρ* = −0.18, R^2^ = 0.05, uncorrected P-value = 0.58, pFDR = 0.85) (Figures 7 & S4, Table S10). Upon examining individual volumetric z-scores and detailed CHD information (see Table S2), we noted that three out of four neonates with extreme negative deviations in TTV (i.e., z ≤ −2.6 SD) had several cardiac defects in addition to being scanned late (after 41 weeks PMA). These neonates also tended to display low oxygen saturation scores at time of scan. One neonate had an AVSD and Tetralogy of Fallot, another displayed an AVSD with coarctation of the aorta, whilst the third had an ASD with persistent *patent foramen ovale*. Although the fourth neonate did not have a CHD, according to clinical notes at time of scan, this neonate suffered from thrombocytopenia and particularly poor feeding in the first few weeks of life. Thus, in certain neonates, it is possible that CHD and low oxygen saturation may be leading to a gradual deviation in whole brain volume from the DS baseline with advancing age at scan.

Much like whole brain volume, several underlying tissue segments were negatively correlated with advancing PMA prior to multiple comparison correction in CHD+ neonates, and not in CHD-neonates. These segments were the cerebellum, the frontal, parietal and occipital GM segments, the parietal WM, the hippocampi, and the thalami (*ρ* = −0.59 to −0.71, R^2^ = 0.35 to 0.49, uncorrected P-value = 0.036 to 0.008, pFDR = 0.07 to 0.05) (Table S11).

Lastly, Figure 8 displays the tissue segments for which the slope or intercept of CHD+ and CHD-linear regressions were significantly different prior to multiple comparison correction (P uncorrected < 0.05) using the F test (Table S10). These were the caudate nuclei, the temporal GM & WM, the parietal WM, and the insular WM (F ratio = 4.89 to 8.09, uncorrected P-value = 0.038 to 0.0095, pFDR = 0.19 for all segments). These segments were particularly clustered in the lateral and posterior regions of the brain, aside from the caudate nucleus. In future, the brain regions with significant age-related volumetric differences between CHD+ and CHD-neonates may become more evident with larger subgroup sizes and higher statistical power.

**Figure 8:**
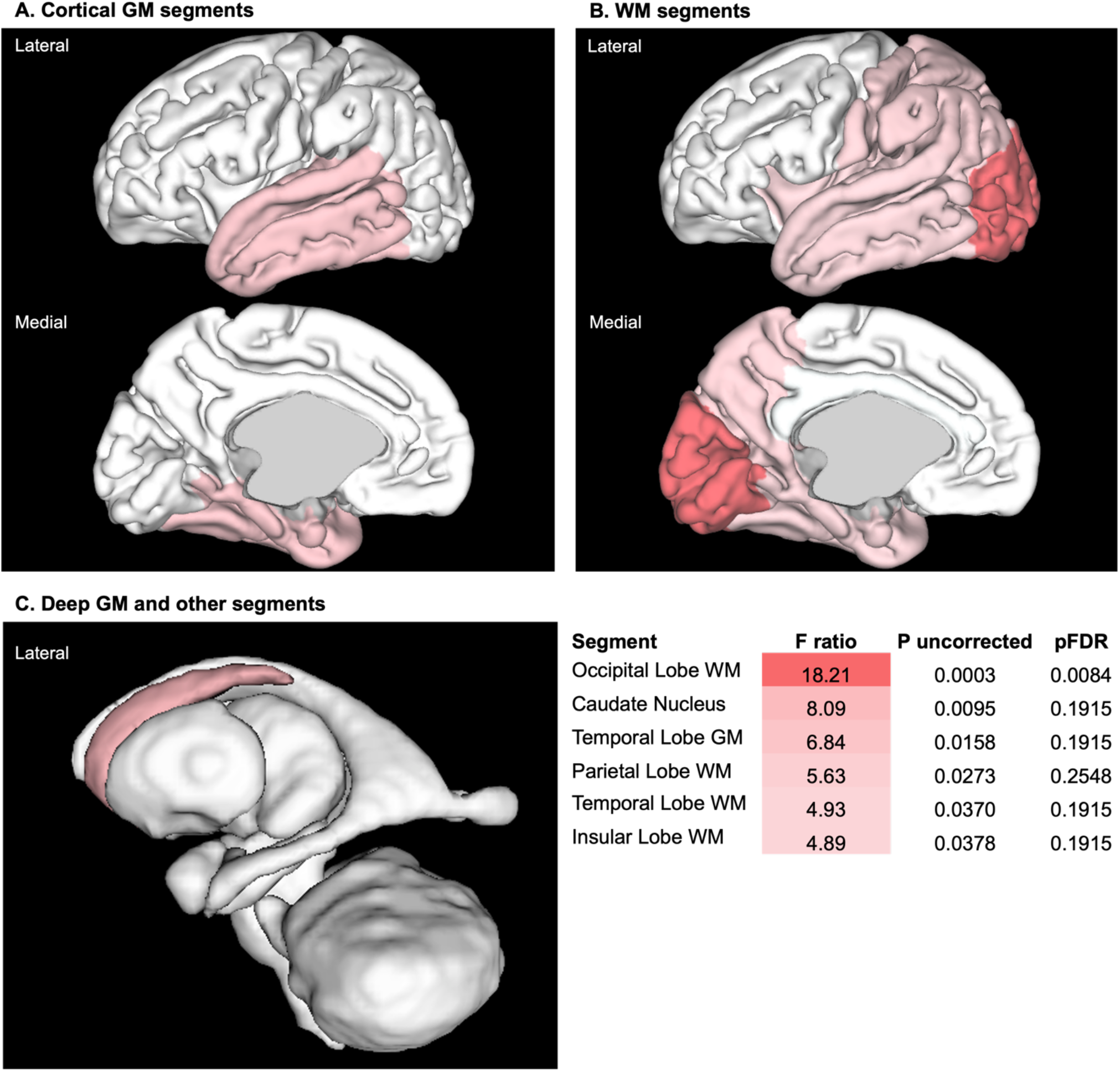
Tissue segments with a difference in slope or intercept between CHD+ and CHD-linear regressions. 3D brain visualisation of the F ratio (from the extra sum-of-squares F test) for tissue segments, in which the slope or intercept of CHD+ and CHD-linear regressions were significantly different prior to multiple comparison correction (P uncorrected < 0.05). Importantly, only the F test for the occipital WM remained significant after multiple comparison correction (pFDR < 0.05) (see Table S10).

### viii) Assessing the impact of other clinical comorbidities on neonatal brain volumes in DS

The possible impact of other clinical comorbidities on neonatal brain volumes were also assessed. Postnatal clinical details for neonates with DS can be found listed in Table S3. Gastrointestinal (GI) malformations, including duodenal atresia and Hirschsprung’s disease were present in 8 out of 25 (32%) of neonates with DS. The same analyses as per CHD+ vs CHD-subgroups were trialled with GI+ and GI-subgroups, as well as other clinically defined subgroups, but these did not show any significant results (data not shown). This was most likely due to small subgroup size, as well as multiple comorbidities. For example, four out of eight neonates with a GI malformation also had CHD. A compounded risk factor for multiple comorbidities was trialled (data not shown), but this did not yield any statistically significant results either.

## DISCUSSION

In this study, we conducted a comprehensive volumetric phenotyping of the neonatal brain in Down syndrome. To the best of our knowledge, the *early brain imaging in DS* study represents the largest dataset of *in vivo* brain imaging in neonates with DS. Robust normative modelling allowed individual inference of volumetric deviation from the normative mean for a given sex, age at scan and age from birth. Although we had a small sample size of neonates with DS, the use of individualised z-scores for each brain segment greatly improved the sensitivity of our analysis compared to traditional volumetry using raw absolute volumes (*data not shown*).

Neuroimaging has successfully identified volumetric differences between DS and appropriate controls from toddlerhood to adolescence^28–41^. However, to date, there have been relatively few *in vivo* brain imaging studies in neonates with DS^30,42^, owing most likely to the specialist nature of MRI at this time point^52,67^. In our study, we corroborated prior neuroimaging findings from fetuses with DS^42,43^, and identified novel differences that have not yet been documented in the developing brain in DS. Moreover, we were able to observe age-related volumetric differences between neonates with and without CHD, indicating that there may be a baseline brain phenotype in neonates with DS, which is further altered in the presence of CHD. The following sections discuss how our key findings relate to other *in vivo* neuroimaging studies in DS, with a focus on the paediatric developmental continuum, as well as results from prior post-mortem tissue analyses of the developing brain in DS.

### i) Several features of the neonatal brain in DS appear to follow a developmental continuum

We observed several volumetric features of the brain in neonates with DS, that were consistent with observations in older cohorts, and which appeared to follow a developmental continuum. Firstly, whole brain volume was significantly smaller in neonates with DS, in line with other paediatric neuroimaging studies ranging from fetal (< 28 gestational weeks, GW)^42^ to adolescent stages^30–32,35,37–40,43^. Whole brain volume also remained smaller in adults with DS^68–70^, although it is important not to confound the effects of early onset Alzheimer’s disease, for which the mean age of diagnosis is 55 years^71,72^. Thus, it appears as though a smaller whole brain volume is likely to begin antenatally and is a lifelong feature in DS, although there is a wide range of individual variation and overlap with the typical population at all stages^41^. Additionally, post-mortem biometric examinations of the developing brain in DS reported a significantly lower total brain weight and occipitofrontal diameter in trisomic fetuses as early as 15 GW^73^.

Despite the presence of a significantly smaller whole brain volume, CSF-filled compartments tended to be enlarged in neonates with DS. The lateral ventricles were previously found to be enlarged in fetuses from 28 GW onwards^42^. Hydrocephalus^74,75^, ventriculomegaly^76^ (especially in very low birth weight infants < 1500g)^77^, and enlargement of the third ventricle^78^ have all been noted in small numbers in DS. McCann et al. have demonstrated continued enlargement of the lateral ventricles throughout childhood into young adulthood with large effect sizes^30^. Consistent with this, in DS murine models (e.g., Ts1Cje, Ts1Rhr), overexpression of purkinje cell protein 4 (pcp4) impaired ciliary function in ependymal cells of the choroid plexus resulting in ventriculomegaly^79^, a process which may potentially underlie ventricular enlargement in human DS amongst other mechanisms.

Although total eCSF volume was not significantly different from control in absolute volume, as observed in prior fetal and neonatal neuroimaging^42^, it was significantly enlarged as a proportion of total intracranial volume. Interestingly, neonates born and scanned preterm (< 37 weeks PMA) exhibited larger relative eCSF volumes, which gradually decreased by later ages at scan. Increased eCSF at term has been associated with prematurity in non-DS infants^80,81^. In children with DS, relative eCSF volume was enlarged from 0 to 5 years old, but no longer after this age bracket^30^. As such, eCSF enlargement appears to be a feature of prematurity in our cohort, but it is not likely to be a continuous feature in DS.

Decreased cerebellar volume, often referred to as cerebellar hypoplasia, is a cardinal and lifelong feature of the brain in DS^5,41^. Smaller cerebellar volume has been evidenced in fetuses (< 28 GW)^42,43^, toddlers^28,30^, children^30,35^, adolescents^30,37,40^ and adults with DS^82^. We found that relative cerebellar volume was significantly and disproportionately small in neonates with DS, representing one of the smallest structures in comparison to control neonates. We also noted that absolute cerebellar volume continued to deviate from the normative mean with advancing age, during this neonatal phase of rapid cerebellar development in control neonates. Post-mortem examinations have evidenced reduced cerebellar transversal diameter from as early as 15 GW^73^, and significant hypocellularity in all cerebellar layers, which is likely due to impaired proliferation of cerebellar precursor cells in DS^83^.

Volumetric features of the deep GM, cortical GM and WM also appear to follow a developmental continuum and are discussed in dedicated sections below. Lastly, regarding the hippocampus and amygdala, we found that, although these were significantly smaller in absolute volume, they were not significantly different from control in relative volume in our neonatal sample. In children^36^ and non-demented adults with DS^84^, hippocampal volume was significantly smaller than control in relative volume, but the amygdala was approximately isometric. From a histological perspective, post-mortem analyses have shown evidence of decreased cell proliferation and increased apoptotic cell death in the hippocampi of fetuses with DS from 17 to 21 GW^85^.

### ii) The deep GM may be “selectively preserved” in DS

We found that the caudate nucleus and thalamus were significantly smaller in absolute volume and were respectively isometric and proportionally enlarged in relative volume. Most surprisingly, the lentiform nucleus (comprising the putamen and pallidum), was not significantly different from control in absolute volume, and thus, disproportionately enlarged in relative volume in neonates with DS. This allometric scaling of the lentiform nucleus and thalamus has been observed in several other paediatric neuroimaging studies in DS. In toddlers with DS, Gunbey et al.^28^ found that the bilateral caudate and thalami were significantly smaller, whilst the putamen and pallidum (particularly the right sides) were not significantly different from control in absolute volume. Moreover, McCann et al.^30^ demonstrated that the putamen, more so than the pallidum, was enlarged as a proportion of total intracranial volume from childhood to young adulthood in DS. In non-demented adults with DS, relative putamen volume was also significantly enlarged^86^, illustrating how this is likely to be a lifelong feature in DS, with an early developmental origin. Pinter et al.^35^ found that adjusted subcortical GM volume was significantly enlarged and “selectively preserved” in a cohort of children and young adults with DS. They eloquently discuss how *“the relatively large size or preservation of [the deep GM] structures in children with DS in the context of significantly smaller overall cerebral volumes suggests that there is a temporal dissociation for the development of cortical versus subcortical [deep GM] regions”*^35^.

During embryonic brain development, the thalamus develops as part of the diencephalon, and regions such as the dorsal thalamus and the pre-thalamus are well established by approximately 40 days post-conception (Carnegie stage 17)^87,88^. Soon after, the basal ganglia (including the caudate, putamen and pallidum) derived from the ganglionic eminences of the ventral telencephalon, can be recognised by approximately 56 days postconception (Carnegie stage 23)^87,88^. It is possible that the thalamus and basal ganglia develop relatively normally in DS throughout embryonic development (up to 8 weeks post-conception), before the onset of major alterations in cortical GM and WM development throughout the second and third trimesters of gestation. Some notable abnormalities in cortical GM development reported in DS, include alterations to neuronal density, laminar organisation, dendritic arborisation and synaptogenesis^89–91^. Thus, it would be of great interest to better understand which morphogenetic processes are most affected during embryonic and fetal brain development in DS, and how these may be differentially impacted by the triplication of *Hsa21*. However, it is important to note that although the basal ganglia and thalami appear to be “selectively preserved” in relative volume, this does not exclude possible dysfunction, and could still reflect underlying issues, such as insufficient programmed cell death during development or aberrant neurocircuitry^35^.

### iii) Regional differences in cortical GM and WM volumes may be linked with brachycephaly, reduced cortical folding, and altered fetal WM development in DS

A brachycephalic cranium with a flat occiput is commonly observed in DS^17,18^. Brachycephaly can be identified on neuroimaging by an increased biparietal diameter (BPD) to occipitofrontal diameter (OFD) ratio approaching the 95^th^ percentile (cephalic index = BPD/OFD × 100)^17^. Post-mortem biometric examinations have been able to identify brachycephaly in fetuses with DS as early as 15 GW, evidenced by a larger BPD width and BPD to head circumference ratio^73^.

Although it is not possible to infer cause-and-effect, in our study we observed regional dynamics in cortical GM and WM lobar volumes, which may be associated with brachycephaly in DS. In particular, the parietal lobe was significantly enlarged in relative volume, in line with an increased BPD reported in DS, whilst the frontal and occipital lobes were significantly reduced in relative volume, in line with a shorter occipitofrontal diameter^17,73,89,90^.

Our findings in the neonate were consistent with observations from several other paediatric neuroimaging studies in DS (spanning from approximately 3 to 20 years). In particular, many of these studies found significantly larger absolute and brain volume-adjusted parietal GM^31,35,38^ and WM volumes^38^. Similarly, adjusted temporal GM and WM^35,38,41^ were enlarged in certain studies. Conversely, adjusted frontal GM was not significantly different, whilst adjusted frontal WM was significantly smaller than control^38,41^. Finally, occipital WM^31,38,41^ was significantly smaller in DS in absolute volume^38,41^. As such, it is likely that the regional dynamics in cortical GM and WM lobar volumes that we have observed in neonates with DS follow a developmental continuum and may be associated with brachycephaly in DS.

Regional alterations in cortical thickness, surface area and folding have also been observed in DS^38,44,92^. Lower average sulcal depth and gyrification has been observed across the brain in fetuses with DS^44^. In children with DS (from 0 to 5 years), cortical thickness was increased, whilst its variability was decreased, indicating abnormal maturation of GM in several regions of the brain^92^. In youth with DS (from 5 to 24 years), cortical thickness was increased throughout much of the frontal, superior parietal and occipital lobes, whilst surface area was reduced in frontal and temporal lobes^38^. As such, it is possible that reduced cortical folding and surface area may underlie the reduced absolute cortical GM volume observed in neonates with DS, whilst increased cortical thickness may be offsetting this reduction in certain regions. It is therefore vital to observe both volumetric and morphometric data in tandem to accurately ascertain alterations to cortical GM in DS.

From a histological perspective, post-mortem analyses have reported a reduction in total neuronal number, as well as a delayed and disorganised lamination of the cortical GM in DS^89,90,93,94^. More specifically, Guidi et al. identified decreased neurogenesis and increased cell death in certain GM regions, such as the hippocampus^85^, the fusiform gyrus and the inferior temporal gyrus^95^ from 17 to 21 GW. Decreased neurogenesis of interneurons has also been reported in DS^96^. As such, greatly reduced neuronal cell proliferation, and increased cell death, may account for the reduced cortical GM volumes noted in DS.

Lastly, we found that the cingulate, frontal, insular and occipital WM segments were significantly reduced in relative volume in neonates with DS. These findings may be supported by a growing body of post-mortem research evidencing aberrant fetal WM development in DS. A postnatal delay in myelination has been noted in DS since the 1980s^89,97^. More recently, Olmos-serrano et al.^98^ conducted a multi-region transcriptomic analysis of DS and euploid brain tissue spanning from the mid-fetal stage to adulthood. They uncovered the co-dysregulation of genes associated with oligodendrocyte differentiation and myelination, which they further validated across species in the Ts65Dn mouse model of DS^98^. Several other studies have reported glial disturbances in the developing brain in DS, including significantly reduced radial glial progenitors^99^, and an imbalance in astro-^100,101^ and oligodendroglial cells^101–103^. This glial imbalance may be due to the altered expression of essential transcription factors for oligodendroglial differentiation^98,103,104^. Thus, it is possible that the cellular, axonal, and perhaps, extracellular matrix compositions of the fetal WM are altered in DS, giving rise to the significantly reduced volumes observed in this study.

### iv) The impact of congenital heart defects on neonatal brain volumes in DS

In our study, the occipital WM displayed a significant difference in age-related volume between CHD+ and CHD-neonates with DS. Occipital WM volumes were significantly smaller in CHD+ neonates by later ages at scan (from approximately 40 weeks PMA). Interestingly, reduced occipital regional volume has been associated with later visual difficulties in preterm infants without DS^105^. It is widely recognised that children with DS have a broad range and high prevalence of visual deficits^106^. Thus, in future, it would be of great interest to associate neonatal occipital WM volume and visual outcomes in infants with DS.

We observed other age-related volumetric differences between CHD+ and CHD-neonates with DS, although these were not statistically significant after multiple comparison correction, most likely due to small subgroup sizes and low statistical power. Fundamentally, whole brain volume tended to be smaller in CHD+ neonates compared to CHD-neonates by later ages at scan. Many underlying specific tissue segments presented with the same trend. Importantly, a high proportion (three out of four neonates) with extreme negative deviations in whole brain volume were scanned at later PMA, had several cardiac defects, and presented low oxygen saturation scores at the time of their neonatal scan. All CHD+ neonates in this study were scanned prior to any cardiac surgery or intervention, although most required surgery within 6 months of life (see supplementary Table S2 for details). Interestingly, body weight at scan, corrected for sex and age differences, was not different between CHD+ and CHD-subgroups, implying that there may be impaired brain growth, over body growth, in neonates with CHD during this neonatal period.

Taken together, our findings may indicate that there is a baseline brain phenotype in neonates with DS, which is further altered in the presence of an associated CHD. We hypothesise that the volumetric differences and trends observed in CHD+ neonates may be due to compromised cardiac function, and reduced cerebral oxygenation in early postnatal life^107–109^. It is also possible that there may be an additional genetic mechanism present in neonates with CHD, as noted in detailed gene mapping experiments using the Dp1Tyb mouse model of DS^110^. In order to better understand the underlying mechanisms affecting neonates with CHD, we could utilise phase contrast angiography to assess cerebral oxygen delivery^48^ and longitudinal follow-up scanning of the same infants over time. Earlier interventions to improve cerebral oxygen delivery may help promote early brain growth and improve developmental outcomes in DS infants with CHD^21,23,24^.

### v) Limitations

The primary limitation of this study was the small sample size of neonates with DS, particularly for clinically defined subgroups (e.g., CHD+ vs CHD-). In the UK, the estimated live birth prevalence for DS was approximately 1.16 in 1000 in 2018, in line with the rest of Europe^9^, whilst an estimated 85.2% of antenatal diagnoses were terminated that year (DSMIG, www.dsmig.org.uk). Our neonatal recruitment is primarily conducted through one site, St Thomas’ Hospital (London), and thus, multi-site recruitment within the UK, or globally, would be needed to significantly increase sample size. In future, retrospective data harmonisation may also be possible, depending on the type of analyses sought. Nevertheless, to the best of our knowledge, our cohort represents the largest dataset of neonates with DS scanned in a prospective study, which is unaffected by variability in acquisition parameters^30,43,44^, in line with a robust neonatal control population^45^.

In our study, participants were each imaged once as a neonate (up to 46 weeks PMA), and we do not have any follow-up scans during the neonatal period. In future, it could be particularly beneficial to conduct follow-up imaging to monitor the continued impact of CHD on neonatal and infant brain measures. Although we have used a well-validated segmentation pipeline optimised for the neonatal brain^55,56^, some current limitations include the lack of automated segmentation for the *cavum septum pellucidum*, the third and fourth ventricles, and cerebellar sub-structures. Structural MRI, and volumetric quantification, cannot tell us detailed information about underlying microstructure and neurocircuitry. In future, we hope to associate data derived from diffusion MRI to complement our volumetric findings. Lastly, although these have not been discussed in this paper, neurodevelopmental outcomes are being collected from participants in our study. This data will be essential to understand if any features of the neonatal brain may serve as early biomarkers for later developmental outcomes in DS.

### Conclusion

In conclusion, we conducted a comprehensive volumetric phenotyping of the neonatal brain in DS. We have demonstrated volumetric groupwise differences, as well as age-related differences in volume, across multiple brain segments between neonates with DS and control. For the first time, we have also identified age-related volumetric differences between neonates with and without CHD in DS.

We observed several volumetric features of the neonatal brain, which appear to follow a developmental continuum in DS, including a reduced whole brain volume; relatively reduced frontal and occipital lobar volumes, in contrast with relatively enlarged temporal and parietal lobar volumes; a “selective preservation”^35^ of the deep GM volume, particularly the lentiform nuclei and thalami; a decreased cerebellar volume; and a tendency for enlargement of the lateral ventricles.

There is a relative scarcity of knowledge pertaining to *in vivo* neonatal brain development in DS. Currently, there are no paediatric longitudinal neuroimaging investigations in DS, starting from the earliest time points, which greatly impedes our understanding of the developmental continuum of neuroanatomical and cognitive parameters in DS. In future, this field of research could greatly benefit from long-term longitudinal imaging and larger sample sizes, which could be delivered by collaborative multi-site investigations. Whilst life expectancy of individuals with DS has greatly improved over the last few decades^5,7^, early interventions may be essential to help support and improve outcomes and quality of life in DS.

## Acknowledgements

We thank the parents and children who participated in this study. The authors gratefully acknowledge staff from the Centre for the Developing Brain (CDB) at King’s College London and the Neonatal Intensive Care Unit (NICU) at St. Thomas’ Hospital. In particular, the research radiologists, radiographers, clinicians, neonatal nurses, midwives, and the administrative teams. In addition, we wish to thank all our obstetric and fetal medicine colleagues from our patient identification sites who have referred participants to us.

