## Supplementary Information for "Comprehensive volumetric phenotyping of the neonatal brain in Down syndrome"

### Index

#### Supplementary Tables

- **Table S1**: Demographic, weight, and head circumference data for neonates with DS.
- **Table S2:** Congenital heart defects (CHD) in neonates with DS.
- **Table S3:** Additional clinical details for neonates with DS.
- **Table S4:** Volumetric brain development in control cohort from 32 to < 46 weeks PMA.
- **Table S5:** Table of results for the extra sum-of-squares F tests comparing DS and control simple linear regressions by brain segment (using absolute z-scores).
- **Table S6:** Table of results for Spearman's rank correlation tests (using absolute z-scores).
- **Table S7:** Table of results for the extra sum-of-squares F tests comparing DS and control simple linear regressions by brain segment (using relative z-scores).
- **Table S8:** Table of results for Spearman's rank correlation tests (using relative z-scores).
- **Table S9:** Group comparison of absolute z-scores between CHD+ and CHD- neonates.
- **Table S10:** Table of results for the extra sum-of-squares F tests comparing simple linear regressions for CHD+ vs. CHD- neonates with DS (using absolute z-scores).
- **Table S11:** Table of results for Spearman's rank correlation tests for CHD+ and CHD- neonates with DS (using absolute z-scores).

#### Supplementary Figures

- **Figure S1:** GPR plots for all segments.
- **Figure S2:** Volumetric brain development in control cohort from 32 to < 46 weeks PMA.
- **Figure S3:** DS and control simple linear regressions by brain segment (using absolute z-scores).
- **Figure S4:** CHD+ and CHD- simple linear regressions by brain segment (using absolute z-scores).
- **Figure S5:** DS and control simple linear regressions by brain segment (using relative z-scores).

### SUPPLEMENTARY TABLES

#### **Table S1:** Demographic, weight, and head circumference (HC) data for neonates with DS.

Data extracted from clinical records on date of scan. HC at birth was missing for 3 neonates with DS.

| ID | Sex | Birth | GA at birth (weeks) | PMA at scan (weeks) | Age from birth (weeks) | Birth Weight (kg) | Birth HC (cm) | Scan Weight (kg) | Scan HC (cm) |
|---|---|---|---|---|---|---|---|---|---|
| **NT21_6** | F | Early Preterm | 31.43 | 43.29 | 11.86 | 1.56 | 28.0 | 3.45 | 36.8 |
| **NT21_8** | F | Late Preterm | 36.43 | 43.00 | 6.57 | 2.28 | - | 3.31 | 33.4 |
| **NT21_9** | M | Late Preterm | 36.57 | 39.14 | 2.57 | 2.58 | 33.0 | 3.19 | 35.3 |
| **NT21_10** | F | Term | 37.57 | 40.57 | 3.00 | 3.13 | 31.5 | 3.40 | 33.6 |
| **NT21_11** | F | Early Preterm | 32.29 | 36.14 | 3.85 | 2.50 | 32.0 | 2.81 | 33.9 |
| **NT21_12** | F | Term | 38.14 | 41.43 | 3.29 | 2.69 | 31.2 | 2.62 | 30.8 |
| **NT21_13** | M | Term | 41.71 | 43.00 | 1.29 | 3.38 | 33.2 | 3.54 | 32.7 |
| **NT21_16** | F | Term | 37.71 | 43.57 | 5.86 | 3.17 | 32.0 | 3.83 | 34.6 |
| **NT21_17** | M | Term | 38.43 | 39.57 | 1.14 | 2.90 | 33.0 | 2.90 | 33.5 |
| **NT21_18** | M | Late Preterm | 36.71 | 44.57 | 7.86 | 3.06 | 33.0 | 3.27 | 33.1 |
| **NT21_19** | M | Term | 37.57 | 44.57 | 7.00 | 2.95 | 31.0 | 4.56 | 36.2 |
| **NT21_21** | F | Early Preterm | 32.00 | 32.43 | 0.43 | 1.86 | 29.5 | 1.86 | 30.0 |
| **NT21_22** | M | Term | 37.00 | 37.57 | 0.57 | 2.50 | 31.5 | 2.40 | 32.3 |
| **NT21_23** | M | Late Preterm | 35.29 | 38.86 | 3.57 | 2.00 | - | 2.20 | 31.5 |
| **NT21_24** | F | Term | 37.00 | 37.86 | 0.86 | 3.01 | 32.0 | 3.01 | 32.7 |
| **NT21_25** | M | Term | 37.14 | 38.00 | 0.86 | 2.33 | 32.0 | 2.26 | 32.2 |
| **NT21_26** | M | Early Preterm | 31.71 | 34.14 | 2.43 | 1.66 | 29.0 | 1.80 | 29.9 |
| **NT21_29** | M | Term | 38.43 | 39.14 | 0.71 | 3.36 | 34.0 | 3.38 | 34.2 |
| **NT21_30** | F | Term | 37.57 | 38.86 | 1.29 | 2.89 | 30.8 | 2.98 | 31.8 |
| **NT21_31** | F | Term | 39.86 | 40.86 | 1.00 | 3.55 | 34.5 | 3.75 | 33.6 |
| **NT21_32** | F | Term | 38.00 | 43.71 | 5.71 | 2.52 | 32.4 | 2.61 | 33.0 |
| **NT21_33** | M | Late Preterm | 36.29 | 42.29 | 6.00 | 2.70 | - | 4.08 | 37.5 |
| **NT21_34** | F | Late Preterm | 36.14 | 40.71 | 4.57 | 2.44 | 31.0 | 3.11 | 34.0 |
| **NT21_35** | M | Term | 38.57 | 45.57 | 7.00 | 3.11 | 33.2 | 3.60 | 36.5 |
| **NT21_36** | M | Term | 37.43 | 39.71 | 2.29 | 2.77 | 32.0 | 2.93 | 33.0 |

**Table S2:** Congenital heart defects (CHD) in neonates with DS.

Data extracted from clinical records on date of scan. **Columns:** in blue = data from clinical records on day of scan, prior to surgery; in yellow = details of later surgery derived from EPR; in green = categorical labels.

| ID | CHD type(s) | CHD details | Evidence of low SPO2 | Age at surgery (months) (EPR) | Surgery details (EPR) | CHD | Cyanotic/ Acyanotic | Severity Classification |
|---|---|---|---|---|---|---|---|---|
| **NT21_6** | | | | | | | | |
| **NT21_8** | AVSD, RAA | Complete AVSD, Bi-directional shunt, Right-sided aortic arch. | Yes (below 90%) | 2.87 | AVSD repair | Yes | Cyanotic | Serious |
| **NT21_9** | | | | | | | | |
| **NT21_10** | | | | | | | | |
| **NT21_11** | Small VSD, PVS, RAA | Small VSD, Pulmonary valve stenosis, Right-sided aortic arch, PDA. | No evidence | N/A | No cardiac surgery. | Yes | Acyanotic | Significant |
| **NT21_12** | AVSD, ToF | AVSD (L to R shunt), ToF (tetralogy canal defect), PDA. | Yes (below 90%) | 4.30 | Elective AVSD closure and PDA ligation | Yes | Cyanotic | Serious |
| **NT21_13** | Small ASD, Persistent PFO | Small ASD, Persistent PFO. | No evidence | N/A | No cardiac surgery. | Yes | Acyanotic | Significant |
| **NT21_16** | AVSD, Small LV, CoA/HAA | Complete unbalanced AVSD, Small LV, Coarctation of the aorta / Hypoplastic aortic arch, PDA. | No evidence (high 90s %) | 3.33 | CoA repair and PA banding | Yes | Acyanotic | Serious |
| **NT21_17** | AVSD, Small LV | Complete AVSD, Small LV, Mild right AV valve regurgitation. | No evidence (high 90s %) | 3.27 | AVSD repair | Yes | Acyanotic | Serious |
| **NT21_18** | | | | | | | | |
| **NT21_19** | | | | | | | | |
| **NT21_21** | | | | | | | | |
| **NT21_22** | AVSD, CoA | Complete AVSD, Coarctation of the aorta. | Yes (below 90%) | 0.37 | CoA repair and PA banding | Yes | Cyanotic | Critical |
| **NT21_23** | Small ASD | Small ASD. | No evidence | N/A | No cardiac surgery. | Yes | Acyanotic | Non-significant |
| **NT21_24** | AVSD | Complete AVSD. | Yes (below 90%) | 5.80 | AVSD repair | Yes | Cyanotic | Serious |
| **NT21_25** | | | | | | | | |
| **NT21_26** | | | | | | | | |
| **NT21_29** | | | | | | | | |
| **NT21_30** | | | | | | | | |
| **NT21_31** | Small VSD, Small ASD | Small VSD, Small secundum ASD. | No evidence | 4.07 | Closure of VSD and ASD | Yes | Acyanotic | Serious |
| **NT21_32** | AVSD, CoA | Complete AVSD, Coarctation of the aorta, Bi-directional shunt. | Yes (below 90%) | 0.13 | CoA repair and PA banding | Yes | Cyanotic | Critical |
| **NT21_33** | | | | | | | | |
| **NT21_34** | Small ASD | Small ASD, Large PDA (L to R shunt). | No evidence | 5.80 | Attempted PDA occlusion by cardiac catheterisation | Yes | Acyanotic | Serious |
| **NT21_35** | | | | | | | | |
| **NT21_36** | AVSD, HLH, HAA | Complete unbalanced AVSD [large VSD (inlet), L-to-R shunt, Primum ASD, Dominant right ventricle and right AV valve, PFO), Hypoplastic left heart, Hypoplastic aortic arch. | Yes (below 90%) | 1.03 | PA banding | Yes | Cyanotic | Critical |

**Abbreviations:** *ASD = Atrial Septal Defect; AVSD = Atrioventricular Septal Defect; CoA = Coarctation of the Aorta; HAA = Hypoplastic Aortic Arch; HLH = Hypoplastic Left Heart Syndrome; LV = Left Ventricle; PDA = Patent Ductus Arteriosus; PFO = Patent Foramen Ovale; PVS = Pulmonary valve stenosis; RAA: Right-sided Aortic Arch; SPO2 = oxygen saturation; ToF = Tetralogy of Fallot; VSD = Ventricular Septal Defect.*

### **Table S3:** Additional clinical details for neonates with DS.

Data extracted from clinical records on date of scan.

| ID | GI issue | GI detail | GI surgery | Infection | Antibiotics | Haematological issue | Metabolic issue | Jaundice / Phototherapy |
|---|---|---|---|---|---|---|---|---|
| **NT21_6** | | | | | | | Hypoglycaemia | Yes |
| **NT21_8** | | | | sepsis suspected | Yes | | | Yes |
| **NT21_9** | Yes | Duodenal Atresia. | Yes | | | Polycythaemia | | Yes |
| **NT21_10** | Yes | Duodenal Atresia. | Yes | | | | | |
| **NT21_11** | Yes | Duodenal stricture. Imperforate anus. | Yes | sepsis suspected | Yes | | Hypoglycaemia | Yes |
| **NT21_12** | Yes | Duodenal Atresia. | Yes | | | | | |
| **NT21_13** | Yes | Hirschsprung Disease. | Yes - later | sepsis confirmed | Yes | Thrombocytopenia (platelet transfusion) | | |
| **NT21_16** | | | | sepsis confirmed | Yes | | Hypoglycaemia | |
| **NT21_17** | | | | | | | | |
| **NT21_18** | | | | sepsis suspected | Yes | Thrombocytopenia | | |
| **NT21_19** | | | | | | | | |
| **NT21_21** | | | | sepsis confirmed | Yes | | | |
| **NT21_22** | | | | | | | | |
| **NT21_23** | Yes | Hirschsprung Disease. | Yes - later | sepsis confirmed | Yes | | | Yes |
| **NT21_24** | | | | | | | | |
| **NT21_25** | | | | | | | | |
| **NT21_26** | Yes | Duodenal Atresia. | Yes | | | | | Yes |
| **NT21_29** | | *Silent aspiration (minor).* | | sepsis suspected | Yes | | | Yes |
| **NT21_30** | Yes | Duodenal Atresia. | Yes | | | Thrombocytopenia, Polycythaemia | | Yes |
| **NT21_31** | | *Abdominal distention (minor)* | | sepsis suspected | Yes | | Raised TSH and/or Hypothyroidism | |
| **NT21_32** | | | | | | | Raised TSH and/or Hypothyroidism, Hypoglycaemia | |
| **NT21_33** | | *Exomphalos resolved* | | | | | | |
| **NT21_34** | | | | sepsis suspected | Yes | | Hypoglycaemia | Yes |
| **NT21_35** | | | | sepsis suspected | Yes | Polycythaemia | | Yes |
| **NT21_36** | | | | | | | Hyponatremia | |

**Abbreviations:** *GI = Gastrointestinal.*
GI surgery: 'Yes' = prior to neonatal scan. 'Yes later' = at a later date, after to neonatal scan.

### **Table S4:** Volumetric brain development in control cohort from 32 to < 46 weeks PMA.

Table detailing mean absolute volume (in $cm^3$) and mean relative volume (in %) for i) whole brain, ii) main tissues, iii) cortical GM segments, iv) WM segments, v) deep GM and other segments for the control cohort at 32- and 45-weeks PMA. Male and female controls neonates have been consolidated. Total % change, as well as an estimated % change per week (pw) are provided.

| | Mean absolute volume (cm3) 32 weeks PMA | Mean absolute volume (cm3) 45 weeks PMA | Total % Change | Est. % Change per week | Mean relative volume (%) 32 weeks PMA | Mean relative volume (%) 45 weeks PMA | Total % Change | Est. % Change per week |
|---|---|---|---|---|---|---|---|---|
| **i) Whole Brain** | | | | | | | | |
| ICV | 234.3 | 539.9 | 130% | 10.0% | - | - | - | - |
| TBV | 187.7 | 444.6 | 137% | 10.5% | - | - | - | - |
| TTV | 183.1 | 437.8 | 139% | 10.7% | - | - | - | - |
| **ii) Main Tissues** | | | | | | | | |
| Cortical Grey Matter | 54.8 | 197.9 | 261% | 20.1% | 28.8% | 45.1% | 56.8% | 4.4% |
| Cerebellum | 9.6 | 33.7 | 252% | 19.4% | 5.2% | 7.7% | 49.0% | 3.8% |
| eCSF | 43.4 | 92.5 | 113% | 8.7% | 20.0% | 16.5% | -17.7% | -1.4% |
| Deep Grey Matter | 15.2 | 31.6 | 108% | 8.3% | 8.5% | 7.3% | -14.7% | -1.1% |
| Brainstem | 3.9 | 7.4 | 93% | 7.1% | 2.2% | 1.7% | -20.6% | -1.6% |
| White Matter | 98.8 | 163.7 | 66% | 5.0% | 54.6% | 37.4% | -31.4% | -2.4% |
| Lateral Ventricles | 3.6 | 5.8 | 60% | 4.6% | 2.4% | 1.3% | -47.4% | -3.6% |
| **iii) Cortical GM** | | | | | | | | |
| Parietal Lobe GM | 12.4 | 47.9 | 285% | 22.0% | 6.5% | 10.9% | 66.4% | 5.1% |
| Occipital Lobe GM | 8.4 | 30.9 | 266% | 20.5% | 4.5% | 7.0% | 56.9% | 4.4% |
| Frontal Lobe GM | 18.2 | 66.2 | 264% | 20.3% | 9.6% | 15.0% | 56.0% | 4.3% |
| Temporal Lobe GM | 11.4 | 41.1 | 262% | 20.1% | 6.0% | 9.3% | 54.8% | 4.2% |
| Insula GM | 1.6 | 4.4 | 174% | 13.4% | 0.9% | 1.0% | 16.2% | 1.2% |
| Cingulate GM | 2.8 | 7.4 | 161% | 12.4% | 1.6% | 1.7% | 9.0% | 0.7% |
| **iv) WM** | | | | | | | | |
| Cingulate WM | 3.2 | 5.8 | 82% | 6.3% | 1.7% | 1.3% | -23.5% | -1.8% |
| Temporal Lobe WM | 18.2 | 32.0 | 76% | 5.8% | 10.1% | 7.4% | -27.0% | -2.1% |
| Frontal Lobe WM | 38.3 | 62.6 | 63% | 4.9% | 21.3% | 14.4% | -32.2% | -2.5% |
| Parietal Lobe WM | 23.1 | 37.5 | 63% | 4.8% | 12.9% | 8.6% | -33.4% | -2.6% |
| Insula WM | 3.7 | 6.0 | 61% | 4.7% | 2.0% | 1.4% | -33.5% | -2.6% |
| Occipital Lobe WM | 10.6 | 16.1 | 52% | 4.0% | 5.9% | 3.6% | -39.1% | -3.0% |
| **v) Deep GM and Other** | | | | | | | | |
| Lentiform Nucleus | 3.1 | 7.7 | 147% | 11.3% | 1.7% | 1.8% | 1.6% | 0.1% |
| Thalamus | 5.1 | 11.1 | 117% | 9.0% | 2.9% | 2.5% | -11.0% | -0.8% |
| Amygdala | 0.5 | 1.1 | 111% | 8.5% | 0.3% | 0.3% | -13.8% | -1.1% |
| Caudate Nucleus | 2.1 | 4.3 | 103% | 7.9% | 1.2% | 1.0% | -17.6% | -1.4% |
| Hippocampus | 1.0 | 1.8 | 76% | 5.9% | 0.6% | 0.4% | -29.8% | -2.3% |

#### Supplementary text:

This table is complementary to Figure S2. The period from 32 up to < 46 weeks PMA at scan was a phase of rapid brain expansion (*section i*). The average intracranial volume (ICV) in control neonates grew more than 2-fold (+130%, an estimated +10% per week, pw) from 234.3 $cm^3$ at 32 weeks to 539.9 $cm^3$ by 45 weeks (males and females consolidated). Similarly, the average total tissue volume (TTV), which represents ICV minus CSF-filled structures (i.e., eCSF and lateral ventricles) grew +139% (estimated +10.7% pw) from 183.1 $cm^3$ to 437.8 $cm^3$.

Looking at the main tissue classes of the brain (*section ii*), we found that the average total cortical GM grew approximately 3.5-fold (+261%, +20.1% pw) from 54.8 $cm^3$ to 197.9 $cm^3$, representing the fastest growing tissue type in the brain during this period. This was composed of, in order from fastest to slowest growing GM segments, the parietal, occipital, frontal, temporal, insular and cingulate GM, which grew between +285% and +161% during this period (*section iii*). Another fast-growing tissue type was the cerebellum, which grew 3.5-fold during this period (+252%, +19.4% pw) from 9.6 to 33.7 $cm^3$. Total deep GM showed a relatively moderate growth of +108% (+8.3% pw), comprised of the lentiform nuclei (+ 147%), the thalami (+117%) and the caudate nuclei (+103%) (*section v*). The brainstem grew relatively moderately compared to other tissue at +93% (+ 7.1% pw, from 3.9 to 7.4 $cm^3$). Total WM only grew +66% (+5.0% pw, from 98.8 to 163.7 $cm^3$) during this period, representing the slowest-growing major tissue type during this period. This was composed of, in order from fastest to slowest growing WM segments, the cingulate, temporal, frontal, parietal, insular and occipital WM, which grew between +82% (+6.3% pw) and +52% (+4.0% pw) during this period (*section iv*). Finally, the eCSF grew moderately (+113%), whilst the lateral ventricles grew the least from 3.6 to 5.8 $cm^3$ (+60%) over this period.

The relative volume of a tissue type indicated its share of the whole brain (as detailed in Table 1 in main text). Total cortical GM grew its relative share of TTV the fastest from 28.8% at 32 weeks to 45.1% at 45 weeks (+ 56.8% change), followed by the cerebellum, which grew from 5.2% to 7.7% of TTV (+ 49.0% change). In contrast, the share of total WM declined from 54.6% to 37.4% of TTV (-31.4% change) during this period. Relative volumes for the brainstem (-20.6%), deep GM structures (-14.7%), eCSF (-17.7%) and lateral ventricles (-47.4%) all declined during this period as they were outcompeted by faster-growing tissue types.

**Table S5:** Table of results for the extra sum-of-squares F tests comparing DS and control simple linear regressions by brain segment (using absolute z-scores).

Table of results for the extra sum-of-squares F tests comparing the parameters (slope and intercept) of DS and control simple linear regressions by brain segment using z-scores derived from absolute volumes. All corresponding simple linear regression plots can be found in Figure S3. 'F' = F ratio, 'DFn' = degree of freedom for the numerator of the F ratio, 'DFd' = degree of freedom for the denominator of the F ratio, 'Sig.' = significance level. The uncorrected P-value, as well as the FDR-corrected P-value are shown. A cell highlighted in green indicates a P-value < 0.05.

| | Extra Sum-of-Squares F Test — DS All (n = 25) vs Control (n = 493) | | | | | | | | | | | | | |
|---|---|---|---|---|---|---|---|---|---|---|---|---|---|---|
| | *Are the slopes equal?* | | | | | | | *Are the elevations or intercepts equal?* | | | | | | |
| | F ratio | DFn | DFd | P value (uncorrected) | Sig | pFDR | Sig | F ratio | DFn | DFd | P value (uncorrected) | Sig | pFDR | Sig |
| **Whole Brain Volumes** | | | | | | | | | | | | | | |
| ICV | 11.45 | 1 | 514 | 0.0008 | *** | 0.0021 | ** | - | - | - | - | - | - | - |
| TTV | 5.42 | 1 | 514 | 0.0203 | * | 0.0338 | * | - | - | - | - | - | - | - |
| **Main Tissue Volumes** | | | | | | | | | | | | | | |
| Cortical GM | 6.32 | 1 | 514 | 0.0122 | * | 0.0222 | * | - | - | - | - | - | - | - |
| Deep GM | 1.63 | 1 | 514 | 0.2026 | ns | 0.2701 | ns | 35.72 | 1 | 515 | <0.0001 | **** | 0.0003 | *** |
| WM | 2.52 | 1 | 514 | 0.1130 | ns | 0.1614 | ns | 78.07 | 1 | 515 | <0.0001 | **** | 0.0003 | *** |
| Cerebellum | 13.15 | 1 | 514 | 0.0003 | *** | 0.0009 | *** | - | - | - | - | - | - | - |
| Brainstem | 6.36 | 1 | 514 | 0.0120 | * | 0.0222 | * | - | - | - | - | - | - | - |
| eCSF | 19.05 | 1 | 514 | <0.0001 | **** | 0.0003 | *** | - | - | - | - | - | - | - |
| Lateral Ventricles | 10.01 | 1 | 514 | 0.0016 | ** | 0.0038 | ** | - | - | - | - | - | - | - |
| Lateral Ventricles (ex-outlier) | 1.61 | 1 | 513 | 0.2054 | ns | | ns | 29.40 | 1 | 514 | <0.0001 | **** | 0.0003 | *** |
| **Cortical GM segments** | | | | | | | | | | | | | | |
| Temporal Lobe GM | 1.52 | 1 | 514 | 0.2189 | ns | 0.2736 | ns | 26.96 | 1 | 515 | <0.0001 | **** | 0.0003 | *** |
| Frontal Lobe GM | 10.10 | 1 | 514 | 0.0016 | ** | 0.0038 | ** | - | - | - | - | - | - | - |
| Parietal Lobe GM | 4.17 | 1 | 514 | 0.0417 | * | 0.0642 | ns | - | - | - | - | - | - | - |
| Occipital Lobe GM | 6.16 | 1 | 514 | 0.0134 | * | 0.0233 | * | - | - | - | - | - | - | - |
| Cingulate Lobe GM | 7.45 | 1 | 514 | 0.0066 | ** | 0.0132 | * | - | - | - | - | - | - | - |
| Insular Lobe GM | 0.34 | 1 | 514 | 0.5589 | ns | 0.5732 | ns | 25.05 | 1 | 515 | <0.0001 | **** | 0.0003 | *** |
| **WM segments** | | | | | | | | | | | | | | |
| Temporal Lobe WM | 0.80 | 1 | 514 | 0.3707 | ns | 0.4119 | ns | 52.16 | 1 | 515 | <0.0001 | **** | 0.0003 | *** |
| Frontal Lobe WM | 3.65 | 1 | 514 | 0.0567 | ns | 0.0840 | ns | 100.80 | 1 | 515 | <0.0001 | **** | 0.0003 | *** |
| Parietal Lobe WM | 0.89 | 1 | 514 | 0.3455 | ns | 0.3949 | ns | 16.71 | 1 | 515 | <0.0001 | **** | 0.0003 | *** |
| Occipital Lobe WM | 1.06 | 1 | 514 | 0.3035 | ns | 0.3679 | ns | 64.97 | 1 | 515 | <0.0001 | **** | 0.0003 | *** |
| Cingulate Lobe WM | 0.92 | 1 | 514 | 0.3386 | ns | 0.3949 | ns | 181.60 | 1 | 515 | <0.0001 | **** | 0.0003 | *** |
| Insular Lobe WM | 2.36 | 1 | 514 | 0.1247 | ns | 0.1720 | ns | 93.57 | 1 | 515 | <0.0001 | **** | 0.0003 | *** |
| **Deep GM & Other** | | | | | | | | | | | | | | |
| Hippocampus | 8.11 | 1 | 514 | 0.0046 | ** | 0.0097 | ** | - | - | - | - | - | - | - |
| Amygdala | 0.69 | 1 | 514 | 0.4071 | ns | 0.4401 | ns | 35.50 | 1 | 515 | <0.0001 | **** | 0.0003 | *** |
| Caudate Nucleus | 1.57 | 1 | 514 | 0.2114 | ns | 0.2728 | ns | 52.97 | 1 | 515 | <0.0001 | **** | 0.0003 | *** |
| Lentiform Nucleus | 0.03 | 1 | 514 | 0.8580 | ns | 0.8580 | ns | 0.54 | 1 | 515 | 0.4642 | ns | 0.4886 | ns |
| Thalamus | 4.80 | 1 | 514 | 0.0290 | * | 0.0464 | * | - | - | - | - | - | - | - |

### **Table S6:** Table of results for Spearman's rank correlation tests (using absolute z-scores).

Table of results for Spearman's rank correlation tests assessing the correlation of absolute z-scores and PMA at scan (in weeks). All corresponding simple linear regression plots can be found in Figure S3. The uncorrected P-value, as well as the FDR-corrected P-value are shown. A cell highlighted in green indicates a P-value < 0.05. The $R^2$ value is provided to assess linear model goodness of fit.

| | ***Correlation of absolute z-scores and PMA at scan*** | | | | | |
|---|---|---|---|---|---|---|
| | **DS All (n = 25)** | | | | | |
| | **Spearman's Rho** | **uncorrected P value** | **Sig.** | **pFDR** | **Sig.** | **$R^2$** |
| **Whole Brain Volumes** | | | | | | |
| ICV | -0.52 | 0.0077 | ** | 0.0824 | ns | 0.24 |
| TTV | -0.36 | 0.0755 | ns | 0.1853 | ns | 0.15 |
| **Main Tissue Volumes** | | | | | | |
| Cortical GM | -0.29 | 0.1536 | ns | 0.2680 | ns | 0.13 |
| Deep GM | -0.12 | 0.5838 | ns | 0.6369 | ns | 0.04 |
| WM | -0.31 | 0.1263 | ns | 0.2436 | ns | 0.11 |
| Cerebellum | -0.54 | 0.0058 | ** | 0.0824 | ns | 0.29 |
| Brainstem | -0.34 | 0.0945 | ns | 0.1963 | ns | 0.17 |
| eCSF | -0.44 | 0.0279 | * | 0.1256 | ns | 0.23 |
| Lateral Ventricles | -0.39 | 0.0540 | ns | 0.1620 | ns | 0.18 |
| **Cortical GM segments** | | | | | | |
| Temporal Lobe GM | -0.11 | 0.6133 | ns | 0.6369 | ns | 0.03 |
| Frontal Lobe GM | -0.38 | 0.0642 | ns | 0.1733 | ns | 0.20 |
| Parietal Lobe GM | -0.19 | 0.3636 | ns | 0.4675 | ns | 0.08 |
| Occipital Lobe GM | -0.47 | 0.0177 | * | 0.0956 | ns | 0.22 |
| Cingulate Lobe GM | -0.41 | 0.0440 | * | 0.1485 | ns | 0.19 |
| Insular Lobe GM | 0.11 | 0.5914 | ns | 0.6369 | ns | 0.01 |
| **WM segments** | | | | | | |
| Temporal Lobe WM | -0.20 | 0.3299 | ns | 0.4454 | ns | 0.03 |
| Frontal Lobe WM | -0.42 | 0.0386 | * | 0.1485 | ns | 0.15 |
| Parietal Lobe WM | -0.26 | 0.2084 | ns | 0.3210 | ns | 0.04 |
| Occipital Lobe WM | -0.35 | 0.0883 | ns | 0.1963 | ns | 0.06 |
| Cingulate Lobe WM | -0.21 | 0.3178 | ns | 0.4454 | ns | 0.05 |
| Insular Lobe WM | -0.17 | 0.4162 | ns | 0.5108 | ns | 0.08 |
| **Deep GM & Other** | | | | | | |
| Hippocampus | -0.49 | 0.0122 | * | 0.0824 | ns | 0.24 |
| Amygdala | -0.11 | 0.5991 | ns | 0.6369 | ns | 0.02 |
| Caudate Nucleus | -0.26 | 0.2140 | ns | 0.3210 | ns | 0.05 |
| Lentiform Nucleus | -0.02 | 0.9229 | ns | 0.9229 | ns | 0.00 |
| Thalamus | -0.29 | 0.1588 | ns | 0.2680 | ns | 0.11 |

**Table S7:** Table of results for the extra sum-of-squares F tests comparing DS and control simple linear regressions by brain segment (using relative z-scores).

Table of results for the extra sum-of-squares F tests comparing the parameters (slope and intercept) of DS and control simple linear regressions by brain segment using z-scores derived from relative volumes. All corresponding simple linear regression plots can be found in Figure S5. ‘F’ = F ratio, ‘DFn’ = degree of freedom for the numerator of the F ratio, ‘DFd’ = degree of freedom for the denominator of the F ratio, ‘Sig.’ = significance level. The uncorrected P-value, as well as the FDR-corrected P-value are shown. A cell highlighted in green indicates a P-value < 0.05.

| | Extra Sum of Squares F Test | | | | | | | | | | | | | |
|---|---|---|---|---|---|---|---|---|---|---|---|---|---|---|
| | DS All (n = 25) vs Control (n = 493) | | | | | | | | | | | | | |
| | *Are the slopes equal?* | | | | | | | *Are the elevations or intercepts equal?* | | | | | | |
| | F ratio | DFn | DFd | P-value (uncorrected) | Sig | pFDR | Sig | F ratio | DFn | DFd | P-value (uncorrected) | Sig | pFDR | Sig |
| **Main Tissue Volumes** | | | | | | | | | | | | | | |
| Cortical GM | 1.87 | 1 | 514 | 0.1720 | ns | 0.2386 | ns | 25.20 | 1 | 515 | <0.0001 | **** | 0.0003 | *** |
| Deep GM | 0.65 | 1 | 514 | 0.4218 | ns | 0.5038 | ns | 47.16 | 1 | 515 | <0.0001 | **** | 0.0003 | *** |
| WM | 2.01 | 1 | 514 | 0.1565 | ns | 0.2243 | ns | 11.68 | 1 | 515 | 0.0007 | *** | 0.0019 | ** |
| Cerebellum | 3.07 | 1 | 514 | 0.0806 | ns | 0.1444 | ns | 92.78 | 1 | 515 | <0.0001 | **** | 0.0003 | *** |
| Brainstem | 2.41 | 1 | 514 | 0.1212 | ns | 0.1861 | ns | 20.61 | 1 | 515 | <0.0001 | **** | 0.0003 | *** |
| eCSF | 22.11 | 1 | 514 | <0.0001 | **** | 0.0003 | *** | - | - | - | - | - | - | - |
| Lateral Ventricles | 38.68 | 1 | 514 | <0.0001 | **** | 0.0003 | *** | - | - | - | - | - | - | - |
| Lateral Ventricles (ex-outlier) | 3.29 | 1 | 513 | 0.0703 | ns | 0.1444 | ns | 81.15 | 1 | 514 | <0.0001 | **** | 0.0003 | **** |
| **Cortical GM segments** | | | | | | | | | | | | | | |
| Temporal Lobe GM | 1.10 | 1 | 514 | 0.2939 | ns | 0.3830 | ns | 40.35 | 1 | 515 | <0.0001 | **** | 0.0003 | *** |
| Frontal Lobe GM | 4.98 | 1 | 514 | 0.0261 | * | 0.0591 | ns | - | - | - | - | - | - | - |
| Parietal Lobe GM | 0.20 | 1 | 514 | 0.6576 | ns | 0.6981 | ns | 73.34 | 1 | 515 | <0.0001 | **** | 0.0003 | *** |
| Occipital Lobe GM | 0.03 | 1 | 514 | 0.8599 | ns | 0.8804 | ns | 0.42 | 1 | 515 | 0.5192 | ns | 0.5875 | |
| Cingulate Lobe GM | 12.14 | 1 | 514 | 0.0005 | *** | 0.0014 | ** | - | - | - | - | - | - | - |
| Insular Lobe GM | 4.78 | 1 | 514 | 0.0292 | * | 0.0628 | ns | - | - | - | - | - | - | - |
| **WM segments** | | | | | | | | | | | | | | |
| Temporal Lobe WM | 0.88 | 1 | 514 | 0.3499 | ns | 0.4376 | ns | 0.19 | 1 | 515 | 0.6656 | ns | 0.6981 | ns |
| Frontal Lobe WM | 0.23 | 1 | 514 | 0.6287 | ns | 0.6932 | ns | 39.33 | 1 | 515 | <0.0001 | **** | 0.0003 | *** |
| Parietal Lobe WM | 0.85 | 1 | 514 | 0.3562 | ns | 0.4376 | ns | 44.58 | 1 | 515 | <0.0001 | **** | 0.0003 | *** |
| Occipital Lobe WM | 3.22 | 1 | 514 | 0.0735 | ns | 0.1374 | ns | 23.27 | 1 | 515 | <0.0001 | **** | 0.0003 | *** |
| Cingulate Lobe WM | 6.85 | 1 | 514 | 0.0092 | ** | 0.0220 | * | - | - | - | - | - | - | - |
| Insular Lobe WM | 1.17 | 1 | 514 | 0.2810 | ns | 0.3776 | ns | 43.69 | 1 | 515 | <0.0001 | **** | 0.0003 | *** |
| **Deep GM & Other** | | | | | | | | | | | | | | |
| Hippocampus | 2.49 | 1 | 514 | 0.1155 | ns | 0.1839 | ns | 2.64 | 1 | 515 | 0.1050 | ns | 0.1737 | ns |
| Amygdala | 2.28 | 1 | 514 | 0.1314 | ns | 0.1948 | ns | 4.52 | 1 | 515 | 0.0341 | * | 0.0698 | ns |
| Caudate Nucleus | 0.00 | 1 | 514 | 0.9468 | ns | 0.9468 | ns | 2.71 | 1 | 515 | 0.1002 | ns | 0.1723 | ns |
| Lentiform Nucleus | 3.45 | 1 | 514 | 0.0639 | ns | 0.1249 | ns | 104.70 | 1 | 515 | <0.0001 | **** | 0.0003 | *** |
| Thalamus | 0.48 | 1 | 514 | 0.4894 | ns | 0.5688 | ns | 42.83 | 1 | 515 | <0.0001 | **** | 0.0003 | *** |

**Table S8:** Table of results for Spearman's rank correlation tests (using relative z-scores).

Table of results for Spearman's rank correlation tests assessing the correlation of relative z-scores and PMA at scan (in weeks). All corresponding simple linear regression plots can be found in Figure S5. The uncorrected P-value, as well as the FDR-corrected P-value are shown. A cell highlighted in green indicates a P-value < 0.05. The $R^2$ value is provided to assess linear model goodness of fit.

| | ***Correlation of relative z-scores and PMA at scan*** | | | | | |
|---|---|---|---|---|---|---|
| | **DS All (n = 25)** | | | | | |
| | **Spearman's Rho** | **uncorrected P value** | **Sig** | **pFDR** | **Sig** | **$R^2$** |
| **Main Tissue Volumes** | | | | | | |
| Cortical GM | -0.02 | 0.9185 | ns | 0.9795 | ns | 0.03 |
| Deep GM | 0.22 | 0.2911 | ns | 0.7278 | ns | 0.02 |
| WM | 0.11 | 0.5978 | ns | 0.9795 | ns | 0.03 |
| Cerebellum | -0.49 | 0.0137 | * | 0.1838 | ns | 0.14 |
| Brainstem | -0.18 | 0.3858 | ns | 0.7881 | ns | 0.10 |
| eCSF | -0.48 | 0.0147 | * | 0.1838 | ns | 0.31 |
| Lateral Ventricles | -0.41 | 0.0423 | * | 0.2115 | ns | 0.33 |
| **Cortical GM segments** | | | | | | |
| Temporal Lobe GM | 0.19 | 0.3646 | ns | 0.7881 | ns | 0.03 |
| Frontal Lobe GM | -0.08 | 0.6914 | ns | 0.9795 | ns | 0.10 |
| Parietal Lobe GM | -0.02 | 0.9185 | ns | 0.9795 | ns | 0.00 |
| Occipital Lobe GM | 0.04 | 0.8493 | ns | 0.9795 | ns | 0.00 |
| Cingulate Lobe GM | -0.33 | 0.1100 | ns | 0.3438 | ns | 0.18 |
| Insular Lobe GM | 0.38 | 0.0616 | ns | 0.2346 | ns | 0.16 |
| **WM segments** | | | | | | |
| Temporal Lobe WM | 0.14 | 0.4900 | ns | 0.8750 | ns | 0.03 |
| Frontal Lobe WM | -0.02 | 0.9403 | ns | 0.9795 | ns | 0.00 |
| Parietal Lobe WM | 0.00 | 0.9898 | ns | 0.9898 | ns | 0.03 |
| Occipital Lobe WM | 0.05 | 0.8193 | ns | 0.9795 | ns | 0.12 |
| Cingulate Lobe WM | 0.44 | 0.0271 | * | 0.2031 | ns | 0.20 |
| Insular Lobe WM | 0.37 | 0.0657 | ns | 0.2346 | ns | 0.02 |
| **Deep GM & Other** | | | | | | |
| Hippocampus | -0.08 | 0.6874 | ns | 0.9795 | ns | 0.09 |
| Amygdala | 0.17 | 0.4098 | ns | 0.7881 | ns | 0.05 |
| Caudate Nucleus | -0.04 | 0.8536 | ns | 0.9795 | ns | 0.00 |
| Lentiform Nucleus | 0.26 | 0.2112 | ns | 0.5867 | ns | 0.13 |
| Thalamus | -0.08 | 0.7159 | ns | 0.9795 | ns | 0.02 |

### **Table S9:** Group comparison of absolute z-scores between CHD+ and CHD- neonates

Table of median absolute z-scores for the DS neonates with CHD+ (n = 13) and without CHD- (n = 12). The table is organised into the following sub-sections: **A)** whole brain volumes, **B)** total tissue volumes, **C)** regional volumes and **D)** specific tissue volumes (including CSF-filled volumes). A non-parametric Kruskal-Wallis test with FDR multiple comparison correction (pFDR) was performed for each tissue label. Uncorrected P-values are also displayed for information. Cliff's delta (*d*) test was used to assess the effect sizes. A colour scale has been applied, whereby red indicates a negative deviation from the normative mean (z < 0, i.e., a smaller volume than norm), white indicates no significant deviation (z = 0), and blue indicates a positive deviation (z > 0, i.e., a larger volume than norm). A cell highlighted in green indicates a P-value < 0.05 or a large effect size.

| **A) Whole Brain Volumes** | **CHD+ (n = 13)** | **CHD- (n = 12)** | **Kruskal Wallis (uncorrected P-value)** | **Kruskal Wallis (pFDR)** | **Sig.** | **Cliff's delta** | **Effect size** |
|---|---|---|---|---|---|---|---|
| ICV | -1.83 | -0.62 | 0.1078 | 0.2668 | ns | -0.35 | medium |
| TBV | -2.08 | -1.02 | 0.0684 | 0.2668 | ns | -0.46 | medium |
| TTV | -2.15 | -1.16 | 0.0785 | 0.2668 | ns | -0.44 | medium |
| **B) Total Tissue Volumes** | | | | | | | |
| Total White Matter | -2.10 | -1.52 | 0.2025 | 0.3159 | ns | -0.40 | medium |
| Cortical Grey Matter | -1.71 | -0.51 | 0.0367 | 0.2455 | ns | -0.53 | large |
| Deep Grey Matter | -1.26 | -0.83 | 0.1058 | 0.2668 | ns | -0.46 | medium |
| **C) Regional Volumes** | | | | | | | |
| Posterior Fossa | -2.74 | -2.08 | 0.4365 | 0.5320 | ns | -0.29 | small |
| Total Cingulate | -2.62 | -1.49 | 0.1367 | 0.2834 | ns | -0.42 | medium |
| Total Frontal Lobe | -2.02 | -1.38 | 0.2293 | 0.3227 | ns | -0.31 | small |
| Total Insula | -2.41 | -1.13 | 0.0631 | 0.2668 | ns | -0.47 | large |
| Total Occipital Lobe | -1.68 | -1.06 | 0.1855 | 0.3014 | ns | -0.45 | medium |
| Total Temporal Lobe | -2.10 | -0.60 | 0.0128 | 0.2455 | ns | -0.58 | large |
| Total Parietal Lobe | -1.13 | -0.44 | 0.1300 | 0.2834 | ns | -0.42 | medium |
| Basal Ganglia | -0.73 | -0.48 | 0.2317 | 0.3227 | ns | -0.29 | small |
| **D) Specific Tissue Volumes** | | | | | | | |
| Cingulate WM | -3.04 | -2.28 | 0.5020 | 0.5594 | ns | -0.33 | medium |
| Cerebellum | -2.98 | -2.18 | 0.4713 | 0.5406 | ns | -0.24 | small |
| Insula WM | -2.65 | -1.52 | 0.1407 | 0.2834 | ns | -0.45 | medium |
| Frontal Lobe WM | -2.36 | -1.79 | 0.4680 | 0.5406 | ns | -0.28 | small |
| Temporal Lobe WM | -2.09 | -0.94 | 0.0504 | 0.2455 | ns | -0.46 | medium |
| Hippocampus | -2.08 | -0.75 | 0.1095 | 0.2668 | ns | -0.38 | medium |
| Caudate Nucleus | -1.98 | -0.82 | 0.0356 | 0.2455 | ns | -0.60 | large |
| Occipital Lobe WM | -1.89 | -1.45 | 0.2267 | 0.3227 | ns | -0.40 | medium |
| Temporal Lobe GM | -1.76 | 0.15 | 0.0064 | 0.2455 | ns | -0.59 | large |
| Cingulate GM | -1.76 | -0.85 | 0.0829 | 0.2668 | ns | -0.41 | medium |
| Occipital Lobe GM | -1.70 | -0.97 | 0.1621 | 0.2927 | ns | -0.36 | medium |
| Brainstem | -1.67 | -0.94 | 0.1821 | 0.3014 | ns | -0.36 | medium |
| Frontal Lobe GM | -1.66 | -0.77 | 0.1453 | 0.2834 | ns | -0.40 | medium |
| Amygdala | -1.42 | -1.11 | 0.3203 | 0.4307 | ns | -0.24 | small |
| Parietal Lobe GM | -1.22 | -0.13 | 0.0476 | 0.2455 | ns | -0.49 | large |
| Parietal Lobe WM | -1.19 | -0.77 | 0.4020 | 0.5057 | ns | -0.26 | small |
| Thalamus | -1.01 | -0.77 | 0.1651 | 0.2927 | ns | -0.28 | small |
| Insula GM | -0.92 | -0.39 | 0.0925 | 0.2668 | ns | -0.46 | medium |
| Lentiform Nucleus | -0.07 | 0.01 | 0.6915 | 0.7097 | ns | -0.09 | negligible |
| eCSF | 0.27 | 0.65 | 0.7726 | 0.7726 | ns | -0.08 | negligible |
| Lateral Ventricles | 1.00 | 1.80 | 0.6902 | 0.7097 | ns | -0.33 | medium |

**Table S10:** Table of results for the extra sum-of-squares F tests comparing simple linear regressions for CHD+ vs. CHD- neonates with DS (using absolute z-scores).

Table of results for the extra sum-of-squares F tests comparing the parameters (slope and intercept) of simple linear regressions using z-scores derived from absolute volumes for CHD+ vs CHD- neonates with DS. All corresponding simple linear regression plots can be found in Figure S4. ‘F’ = F ratio, ‘DFn’ = degree of freedom for the numerator of the F ratio, ‘DFd’ = degree of freedom for the denominator of the F ratio, ‘Sig.’ = significance level. The uncorrected P-value, as well as the FDR-corrected P-value are shown. A cell highlighted in green indicates a P-value < 0.05.

| | Extra Sum-of-Squares F Test | | | | | | | | | | | | | |
|---|---|---|---|---|---|---|---|---|---|---|---|---|---|---|
| | CHD+ (n = 13) vs CHD- (n = 12) | | | | | | | | | | | | | |
| | *Are the slopes equal?* | | | | | | | *Are the elevations or intercepts equal?* | | | | | | |
| | F ratio | DFn | DFd | P-value (uncorrected) | Sig | pFDR | Sig | F ratio | DFn | DFd | P-value (uncorrected) | Sig | pFDR | Sig |
| **Whole Brain Volumes** | | | | | | | | | | | | | | |
| ICV | 2.14 | 1 | 21 | 0.1584 | ns | 0.2977 | ns | 2.01 | 1 | 22 | 0.1705 | ns | 0.2243 | ns |
| TTV | 3.07 | 1 | 21 | 0.0945 | ns | 0.2646 | ns | 3.86 | 1 | 22 | 0.0621 | ns | 0.1915 | ns |
| **Main Tissue Volumes** | | | | | | | | | | | | | | |
| Cortical GM | 2.05 | 1 | 21 | 0.1670 | ns | 0.2977 | ns | 3.68 | 1 | 22 | 0.0680 | ns | 0.1915 | ns |
| Deep GM | 2.27 | 1 | 21 | 0.1470 | ns | 0.2977 | ns | 3.84 | 1 | 22 | 0.0628 | ns | 0.1915 | ns |
| WM | 3.27 | 1 | 21 | 0.0851 | ns | 0.2646 | ns | 2.84 | 1 | 22 | 0.1059 | ns | 0.1915 | ns |
| Cerebellum | 3.75 | 1 | 21 | 0.0664 | ns | 0.2646 | ns | 2.43 | 1 | 22 | 0.1331 | ns | 0.1915 | ns |
| Brainstem | 1.03 | 1 | 21 | 0.3208 | ns | 0.4251 | ns | 2.43 | 1 | 22 | 0.1334 | ns | 0.1915 | ns |
| eCSF | 0.36 | 1 | 21 | 0.5573 | ns | 0.5790 | ns | 0.01 | 1 | 22 | 0.9200 | ns | 0.9200 | ns |
| Lateral Ventricles | 0.05 | 1 | 21 | 0.8303 | ns | 0.8303 | ns | 2.84 | 1 | 22 | 0.1063 | ns | 0.1915 | ns |
| **Cortical GM segments** | | | | | | | | | | | | | | |
| Temporal Lobe GM | 2.02 | 1 | 21 | 0.1701 | ns | 0.2977 | ns | 6.84 | 1 | 22 | 0.0158 | * | 0.1915 | ns |
| Frontal Lobe GM | 1.19 | 1 | 21 | 0.2879 | ns | 0.4031 | ns | 1.61 | 1 | 22 | 0.2184 | ns | 0.2730 | ns |
| Parietal Lobe GM | 2.53 | 1 | 21 | 0.1269 | ns | 0.2961 | ns | 3.98 | 1 | 22 | 0.0585 | ns | 0.1915 | ns |
| Occipital Lobe GM | 3.13 | 1 | 21 | 0.0915 | ns | 0.2646 | ns | 2.61 | 1 | 22 | 0.1204 | ns | 0.1915 | ns |
| Cingulate Lobe GM | 0.45 | 1 | 21 | 0.5097 | ns | 0.5709 | ns | 3.36 | 1 | 22 | 0.0802 | ns | 0.1915 | ns |
| Insular Lobe GM | 0.98 | 1 | 21 | 0.3340 | ns | 0.4251 | ns | 2.61 | 1 | 22 | 0.1202 | ns | 0.1915 | ns |
| **WM segments** | | | | | | | | | | | | | | |
| Temporal Lobe WM | 2.63 | 1 | 21 | 0.1201 | ns | 0.2961 | ns | 4.93 | 1 | 22 | 0.0370 | * | 0.1915 | ns |
| Frontal Lobe WM | 0.79 | 1 | 21 | 0.3845 | ns | 0.4544 | ns | 1.01 | 1 | 22 | 0.3253 | ns | 0.3697 | ns |
| Parietal Lobe WM | 5.63 | 1 | 21 | 0.0273 | * | 0.2548 | ns | - | - | - | - | - | - | - |
| Occipital Lobe WM | 18.21 | 1 | 21 | 0.0003 | *** | 0.0084 | ** | - | - | - | - | - | - | - |
| Cingulate Lobe WM | 0.77 | 1 | 21 | 0.3895 | ns | 0.4544 | ns | 2.52 | 1 | 22 | 0.1265 | ns | 0.1915 | ns |
| Insular Lobe WM | 0.35 | 1 | 21 | 0.5583 | ns | 0.5790 | ns | 4.89 | 1 | 22 | 0.0378 | * | 0.1915 | ns |
| **Deep GM & Other** | | | | | | | | | | | | | | |
| Hippocampus | 1.73 | 1 | 21 | 0.2032 | ns | 0.3161 | ns | 2.37 | 1 | 22 | 0.1379 | ns | 0.1915 | ns |
| Amygdala | 3.32 | 1 | 21 | 0.0826 | ns | 0.2646 | ns | 1.08 | 1 | 22 | 0.3106 | ns | 0.3697 | ns |
| Caudate Nucleus | 4.10 | 1 | 21 | 0.0557 | ns | 0.2646 | ns | 8.09 | 1 | 22 | 0.0095 | ** | 0.1915 | ns |
| Lentiform Nucleus | 1.84 | 1 | 21 | 0.1895 | ns | 0.3121 | ns | 0.63 | 1 | 22 | 0.4360 | ns | 0.4542 | ns |
| Thalamus | 3.18 | 1 | 21 | 0.0892 | ns | 0.2646 | ns | 2.96 | 1 | 22 | 0.0992 | ns | 0.1915 | ns |

### **Table S11:** Table of results for Spearman's rank correlation tests for CHD+ and CHD- neonates with DS (using absolute z-scores).

Table of results for Spearman's rank correlation tests assessing the correlation of absolute z-scores and PMA at scan (in weeks) for CHD+ and CHD- neonates with DS. All corresponding simple linear regression plots can be found in Figure S4. The uncorrected P-value, as well as the FDR-corrected P-value are shown. A cell highlighted in green indicates a P-value < 0.05. The $R^2$ value is provided to assess linear model goodness of fit.

| | ***Correlation of absolute z-scores and PMA at scan*** | | | | | | | | | | | |
|---|---|---|---|---|---|---|---|---|---|---|---|---|
| | **CHD+ (n = 13)** | | | | | | **CHD- (n = 12)** | | | | | |
| | **Spearman's Rho** | **uncorrected P value** | **Sig** | **pFDR** | **Sig** | **R squared** | **Spearman's Rho** | **uncorrected P value** | **Sig** | **pFDR** | **Sig** | **R squared** |
| **Whole Brain Volumes** | | | | | | | | | | | | |
| ICV | -0.67 | 0.0149 | * | 0.0503 | ns | 0.48 | -0.40 | 0.1929 | ns | 0.6931 | ns | 0.15 |
| TTV | -0.73 | 0.0059 | ** | 0.0503 | ns | 0.48 | -0.18 | 0.5832 | ns | 0.8506 | ns | 0.05 |
| **Main Tissue Volumes** | | | | | | | | | | | | |
| Cortical GM | -0.69 | 0.0110 | * | 0.0503 | ns | 0.41 | -0.12 | 0.7111 | ns | 0.8932 | ns | 0.05 |
| Deep GM | -0.40 | 0.1706 | ns | 0.2063 | ns | 0.23 | 0.04 | 0.9076 | ns | 0.9602 | ns | 0.00 |
| WM | -0.63 | 0.0238 | * | 0.0663 | ns | 0.42 | -0.19 | 0.5601 | ns | 0.8506 | ns | 0.02 |
| Cerebellum | -0.71 | 0.0080 | ** | 0.0503 | ns | 0.48 | -0.43 | 0.1655 | ns | 0.6931 | ns | 0.26 |
| Brainstem | -0.39 | 0.1834 | ns | 0.2063 | ns | 0.22 | -0.37 | 0.2373 | ns | 0.6931 | ns | 0.20 |
| eCSF | -0.39 | 0.1834 | ns | 0.2063 | ns | 0.28 | -0.44 | 0.1581 | ns | 0.6931 | ns | 0.21 |
| Lateral Ventricles | -0.51 | 0.0781 | ns | 0.1172 | ns | 0.11 | -0.39 | 0.2145 | ns | 0.6931 | ns | 0.22 |
| **Cortical GM segments** | | | | | | | | | | | | |
| Temporal Lobe GM | -0.54 | 0.0615 | ns | 0.1107 | ns | 0.34 | 0.19 | 0.5450 | ns | 0.8506 | ns | 0.00 |
| Frontal Lobe GM | -0.62 | 0.0261 | * | 0.0663 | ns | 0.37 | -0.27 | 0.3926 | ns | 0.8154 | ns | 0.14 |
| Parietal Lobe GM | -0.59 | 0.0351 | * | 0.0752 | ns | 0.40 | 0.06 | 0.8470 | ns | 0.9529 | ns | 0.01 |
| Occipital Lobe GM | -0.67 | 0.0138 | * | 0.0503 | ns | 0.49 | -0.32 | 0.3150 | ns | 0.7603 | ns | 0.13 |
| Cingulate Lobe GM | -0.52 | 0.0711 | ns | 0.1129 | ns | 0.20 | -0.55 | 0.0687 | ns | 0.6931 | ns | 0.24 |
| Insular Lobe GM | -0.22 | 0.4614 | ns | 0.4614 | ns | 0.09 | 0.45 | 0.1406 | ns | 0.6931 | ns | 0.00 |
| **WM segments** | | | | | | | | | | | | |
| Temporal Lobe WM | -0.55 | 0.0557 | ns | 0.1074 | ns | 0.38 | 0.02 | 0.9602 | ns | 0.9602 | ns | 0.00 |
| Frontal Lobe WM | -0.41 | 0.1645 | ns | 0.2063 | ns | 0.21 | -0.44 | 0.1581 | ns | 0.6931 | ns | 0.14 |
| Parietal Lobe WM | -0.69 | 0.0119 | * | 0.0503 | ns | 0.44 | 0.11 | 0.7278 | ns | 0.8932 | ns | 0.01 |
| Occipital Lobe WM | -0.89 | <0.0001 | **** | 0.0027 | ** | 0.78 | 0.15 | 0.6301 | ns | 0.8506 | ns | 0.05 |
| Cingulate Lobe WM | -0.46 | 0.1110 | ns | 0.1499 | ns | 0.12 | -0.02 | 0.9426 | ns | 0.9602 | ns | 0.02 |
| Insular Lobe WM | -0.28 | 0.3456 | ns | 0.3589 | ns | 0.10 | -0.30 | 0.3379 | ns | 0.7603 | ns | 0.10 |
| **Deep GM & Other** | | | | | | | | | | | | |
| Hippocampus | -0.62 | 0.0270 | * | 0.0663 | ns | 0.46 | -0.35 | 0.2567 | ns | 0.6931 | ns | 0.17 |
| Amygdala | -0.50 | 0.0873 | ns | 0.1241 | ns | 0.34 | 0.24 | 0.4588 | ns | 0.8506 | ns | 0.01 |
| Caudate Nucleus | -0.52 | 0.0711 | ns | 0.1129 | ns | 0.32 | -0.20 | 0.5303 | ns | 0.8506 | ns | 0.00 |
| Lentiform Nucleus | -0.37 | 0.2180 | ns | 0.2354 | ns | 0.12 | 0.17 | 0.5987 | ns | 0.8506 | ns | 0.04 |
| Thalamus | -0.59 | 0.0362 | * | 0.0752 | ns | 0.35 | -0.09 | 0.7696 | ns | 0.9034 | ns | 0.03 |

### SUPPLEMENTARY FIGURES

**Figure S1:** GPR plots for all segments.

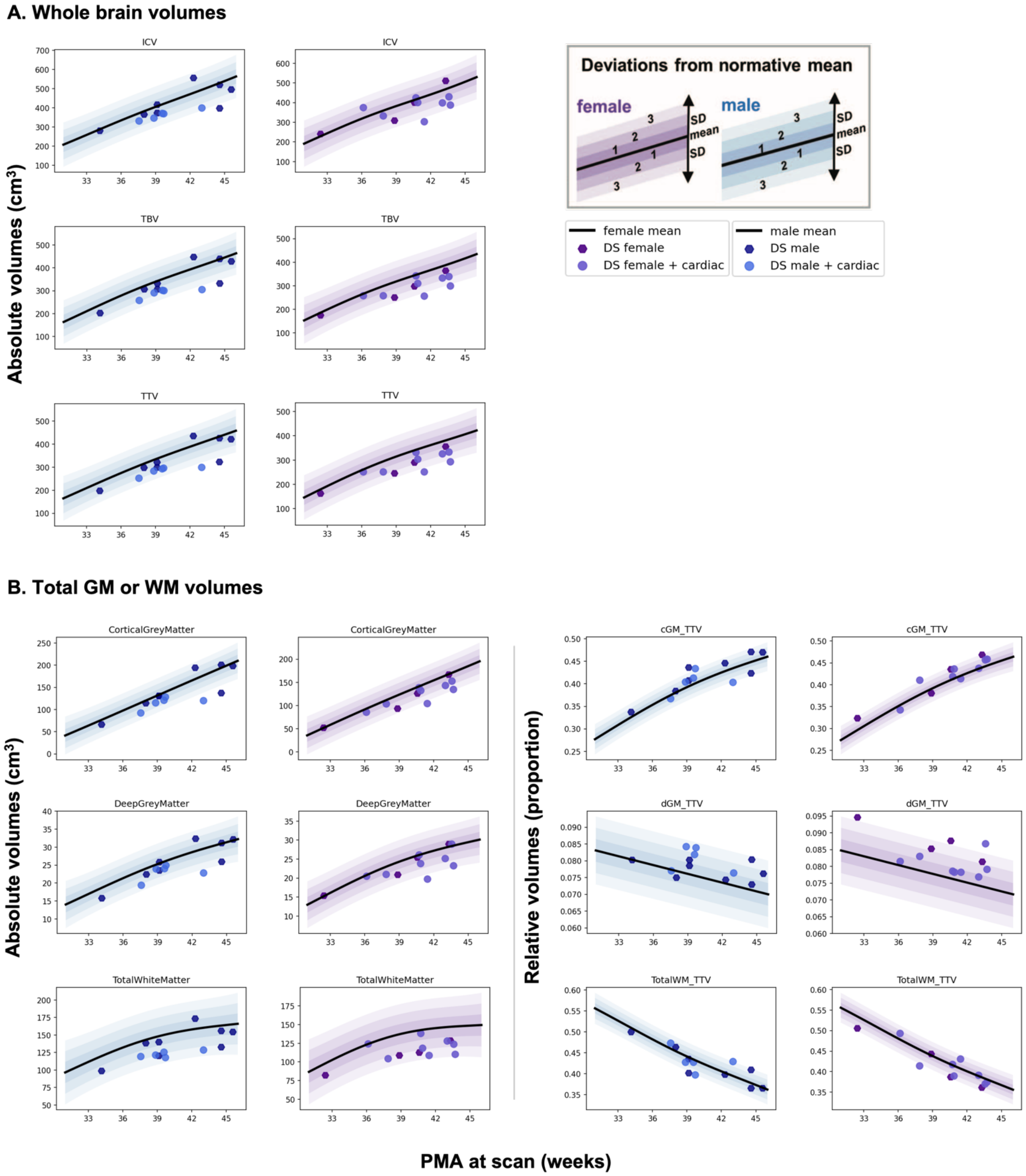

### C. Regional volumes

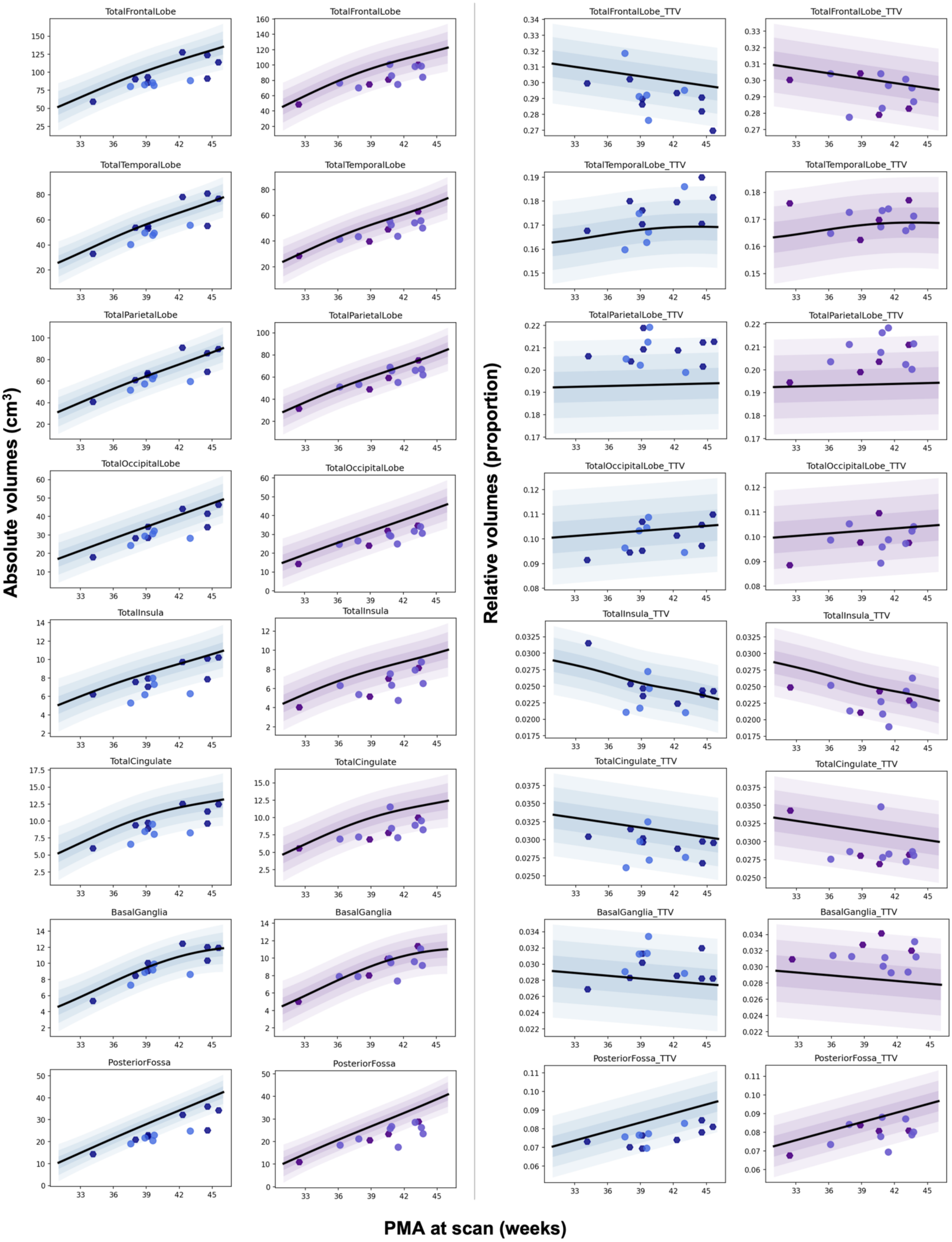

### D.1. Specific tissue volumes – cortical GM segments

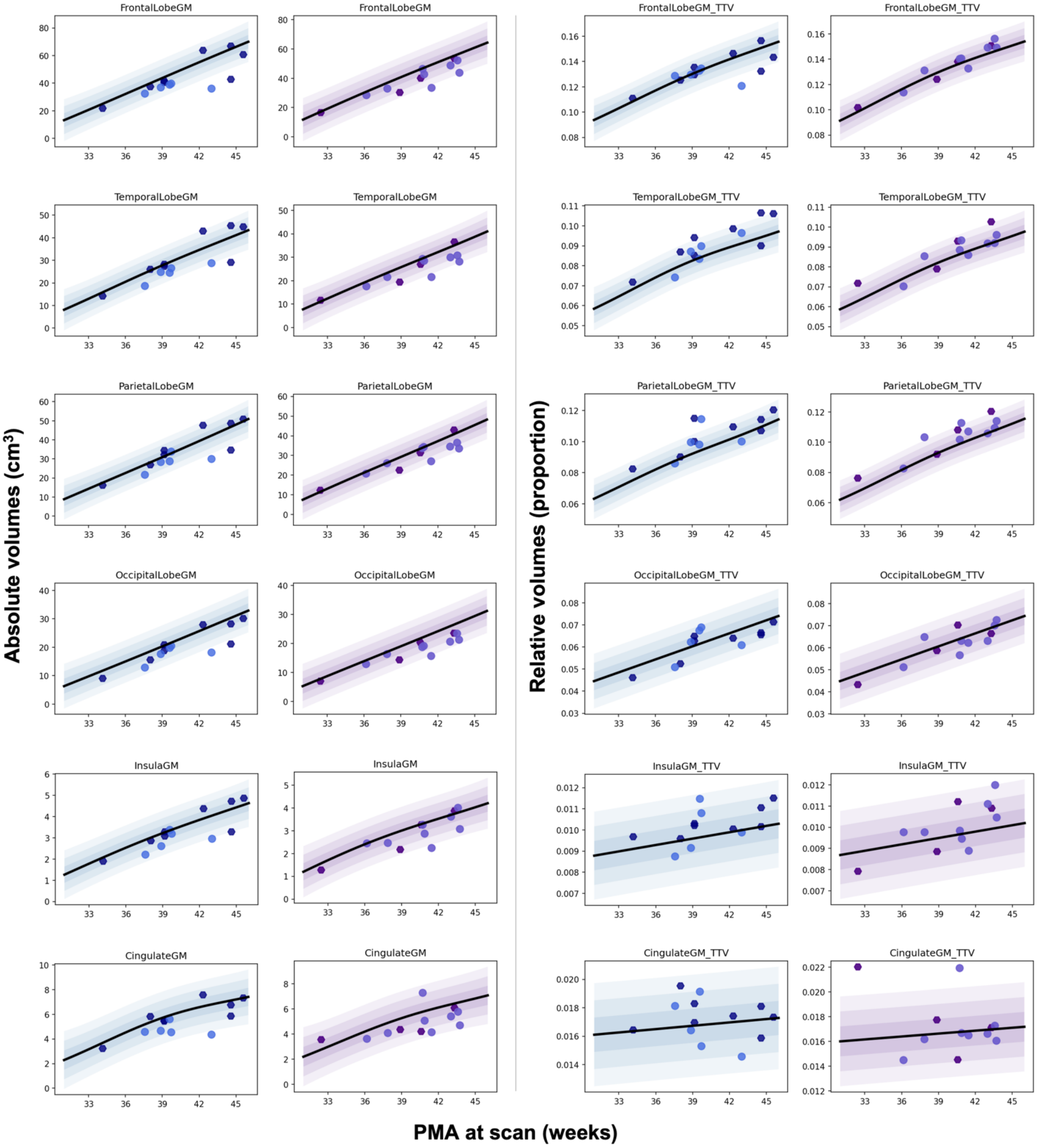

#### D.2. Specific tissue volumes – WM segments

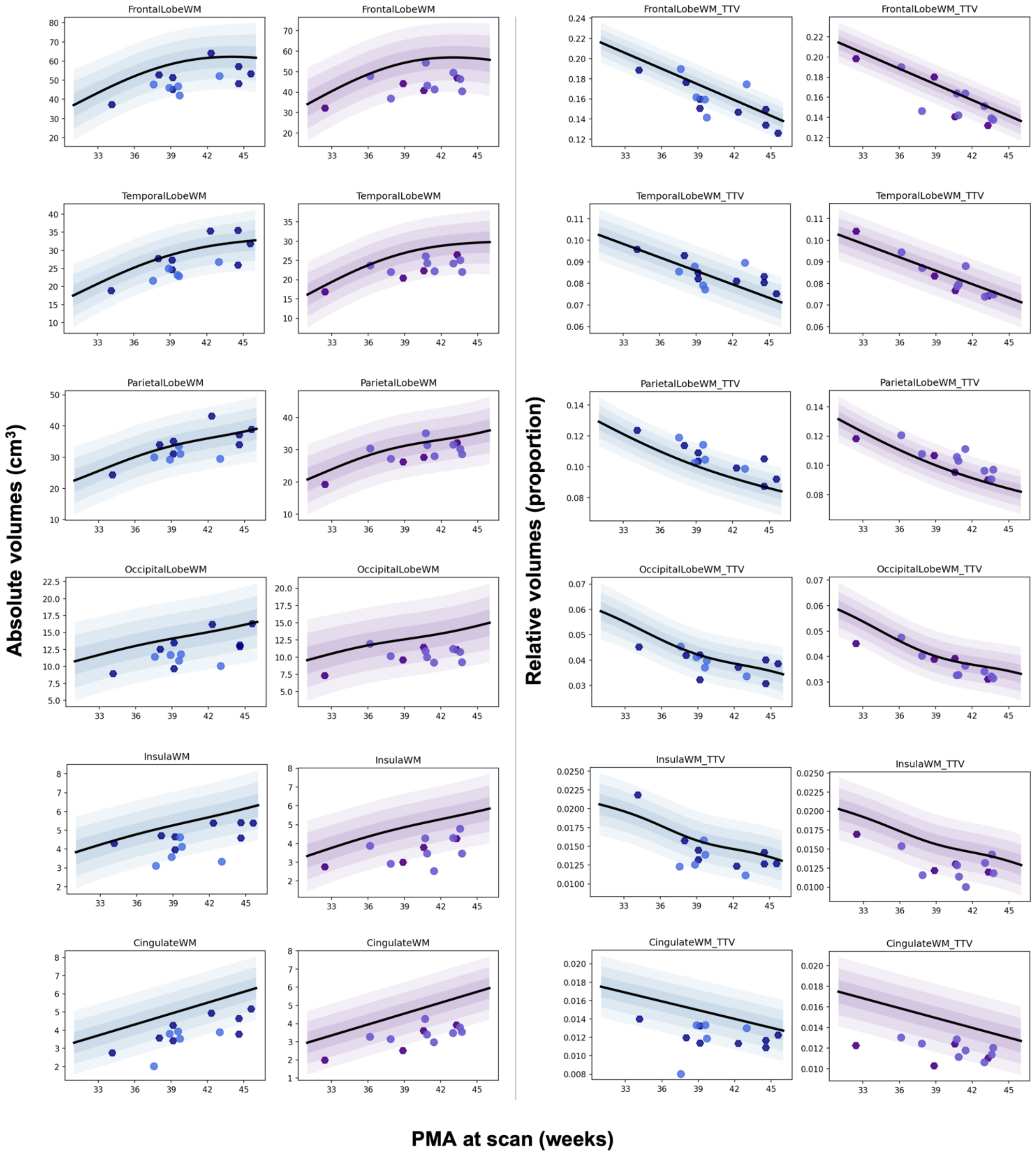

### D.3. Specific tissue volumes – deep GM and other segments

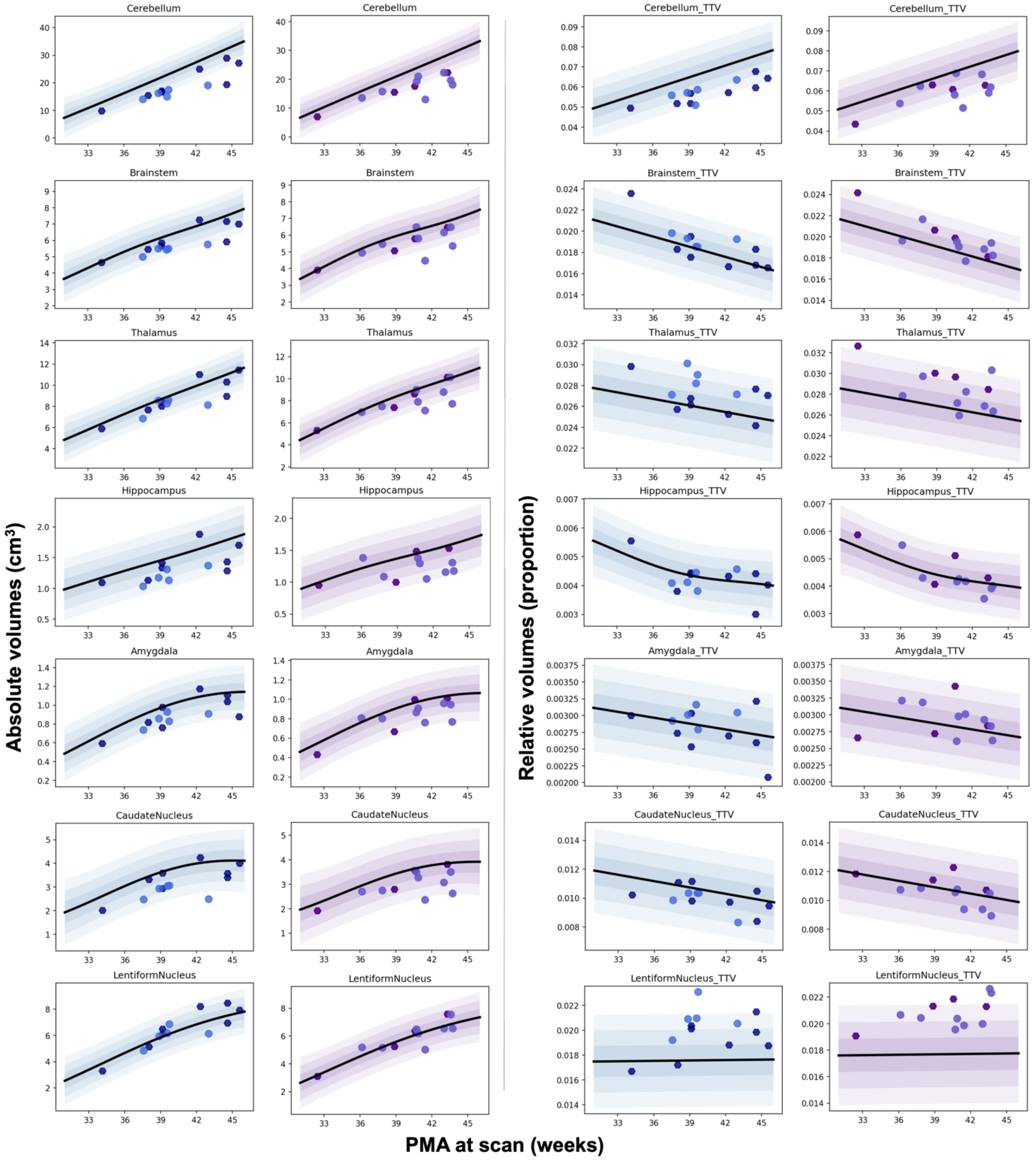

**Figure S2:** Volumetric brain development in control cohort from 32 to < 46 weeks PMA.

Scatter plots of absolute (in $cm^3$) or relative volumes (proportion) from 32 to < 46 weeks PMA for n = 493 preterm to term-born control neonates (females and males consolidated). Data was fitted with a Gaussian curve (bolded curve) and 95% confidence intervals (dotted lines). Plots for **A)** whole brain volumes (in $cm^3$), **B)** main tissue classes (in $cm^3$) and **C)** main tissue classes (in relative volume), **D)** cortical GM segments (in $cm^3$), **E)** WM segments (in $cm^3$), **F)** Deep GM and other segments (in $cm^3$), **G)** cortical GM segments (in relative volume), **H)** WM segments (in relative volume) and **I)** Deep GM and other segments (in relative volume).

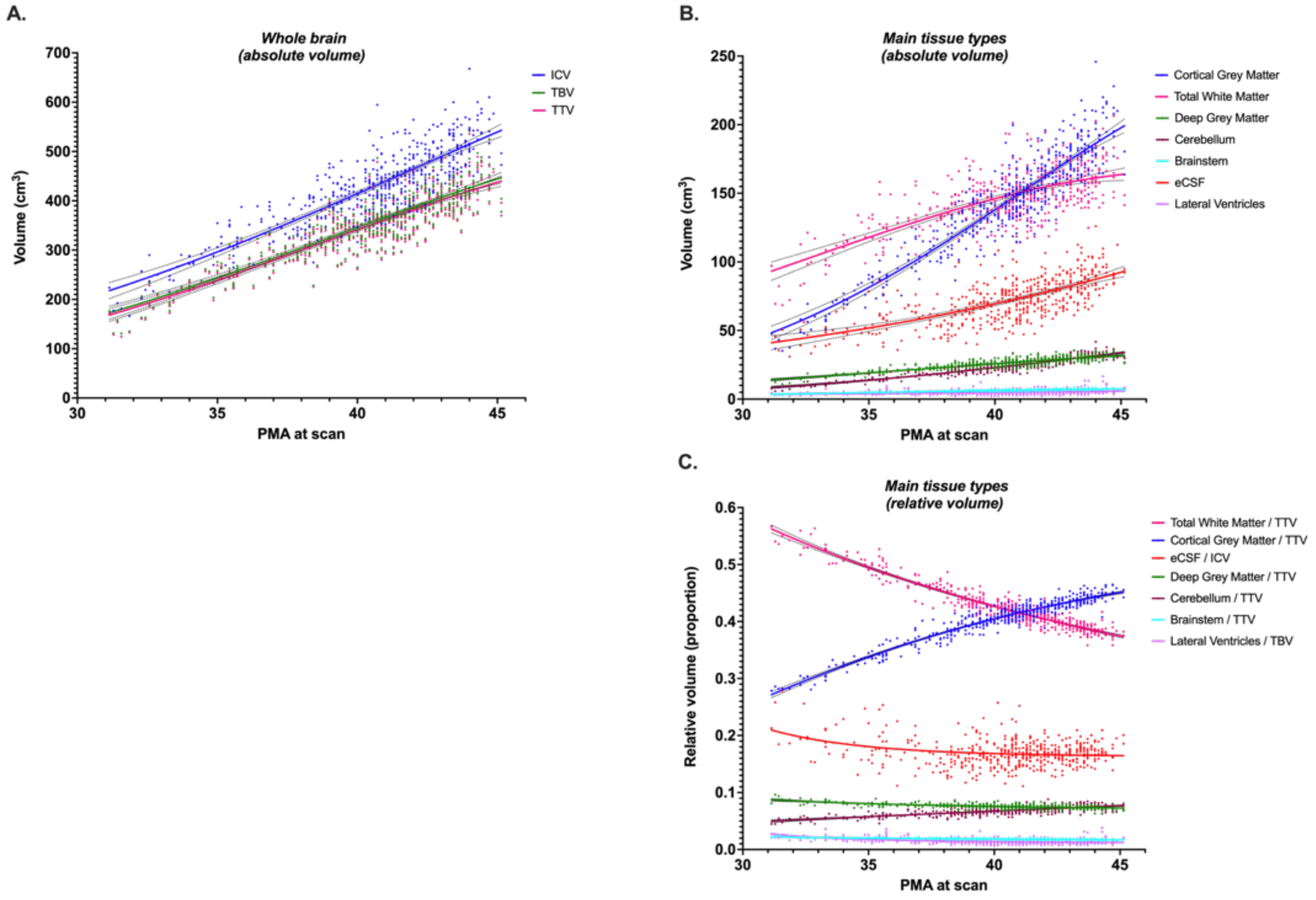

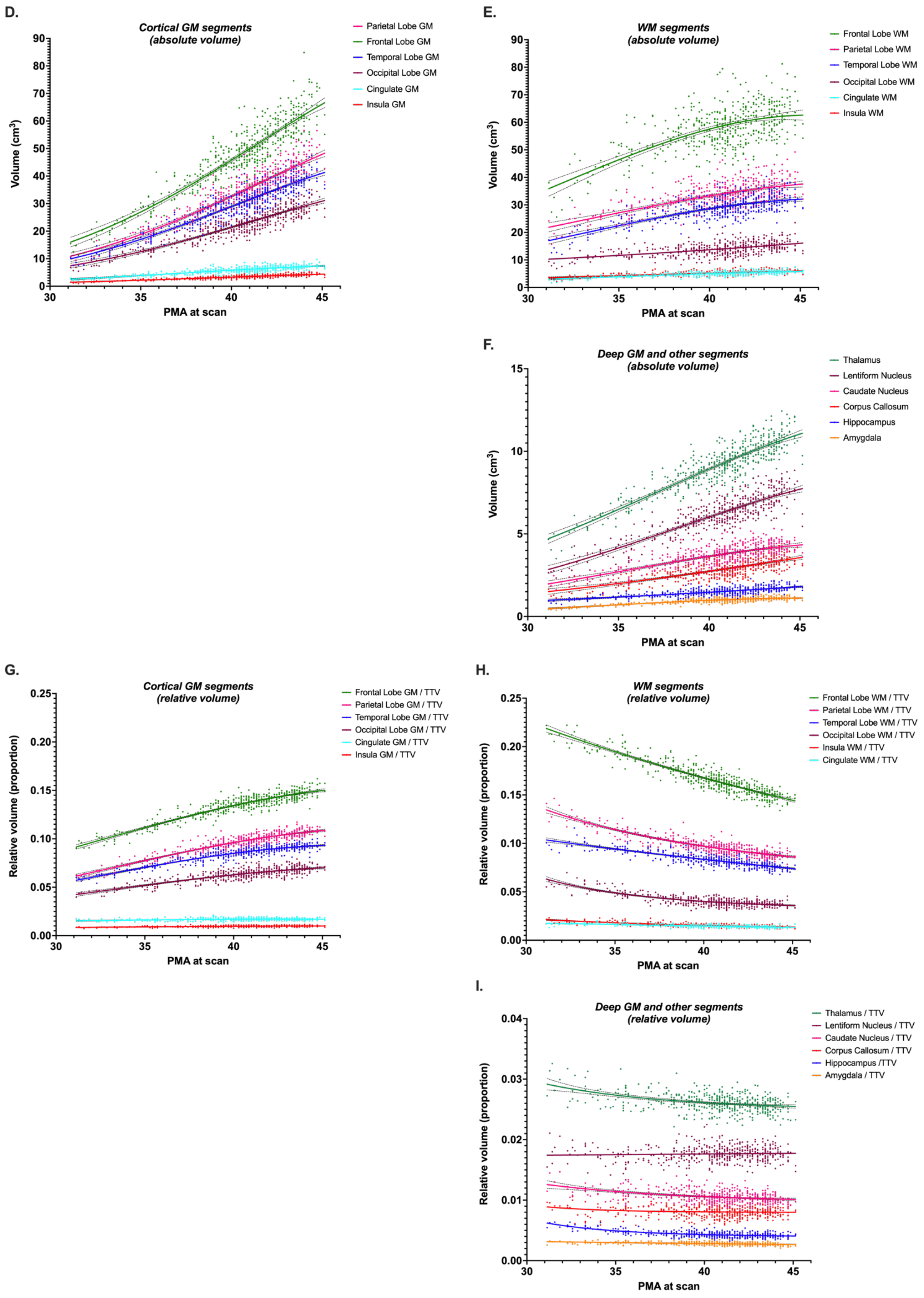
D.
Cortical GM segments
(absolute volume)
Parietal Lobe GM
Frontal Lobe GM
Temporal Lobe GM
Occipital Lobe GM
Cingulate GM
Insula GM
Volume (cm³)
PMA at scan
E.
WM segments
(absolute volume)
Frontal Lobe WM
Parietal Lobe WM
Temporal Lobe WM
Occipital Lobe WM
Cingulate WM
Insula WM
Volume (cm³)
PMA at scan
F.
Deep GM and other segments
(absolute volume)
Thalamus
Lentiform Nucleus
Caudate Nucleus
Corpus Callosum
Hippocampus
Amygdala
Volume (cm³)
PMA at scan
G.
Cortical GM segments
(relative volume)
Frontal Lobe GM / TTV
Parietal Lobe GM / TTV
Temporal Lobe GM / TTV
Occipital Lobe GM / TTV
Cingulate GM / TTV
Insula GM / TTV
Relative volume (proportion)
PMA at scan
H.
WM segments
(relative volume)
Frontal Lobe WM / TTV
Parietal Lobe WM / TTV
Temporal Lobe WM / TTV
Occipital Lobe WM / TTV
Insula WM / TTV
Cingulate WM / TTV
Relative volume (proportion)
PMA at scan
I.
Deep GM and other segments
(relative volume)
Thalamus / TTV
Lentiform Nucleus / TTV
Caudate Nucleus / TTV
Corpus Callosum / TTV
Hippocampus /TTV
Amygdala / TTV
Relative volume (proportion)
PMA at scan

**<u>Figure S3:</u>** <u>DS and control simple linear regressions by brain segment (using absolute z-scores).</u>

Simple linear regression plots of absolute z-scores against PMA at scan (in weeks) appear as coloured lines with 95% confidence intervals as dotted lines for neonates with DS (n = 25, males & females). Dots for individual control neonates (n = 493, males & females) appear in grey, and simple linear regressions appear as flat black lines at z = 0 with 95% confidence intervals as dotted lines. Plots for **A.** whole brain, **B.** cortical GM segments, **C.** WM segments, **D.** the total deep GM and other segments. Main tissue volumes can be found in Figure 6 of main text**).** A table of results for F-tests can be found in Table S5 and Spearman's correlation in Table S6.

A. Whole brain volumes

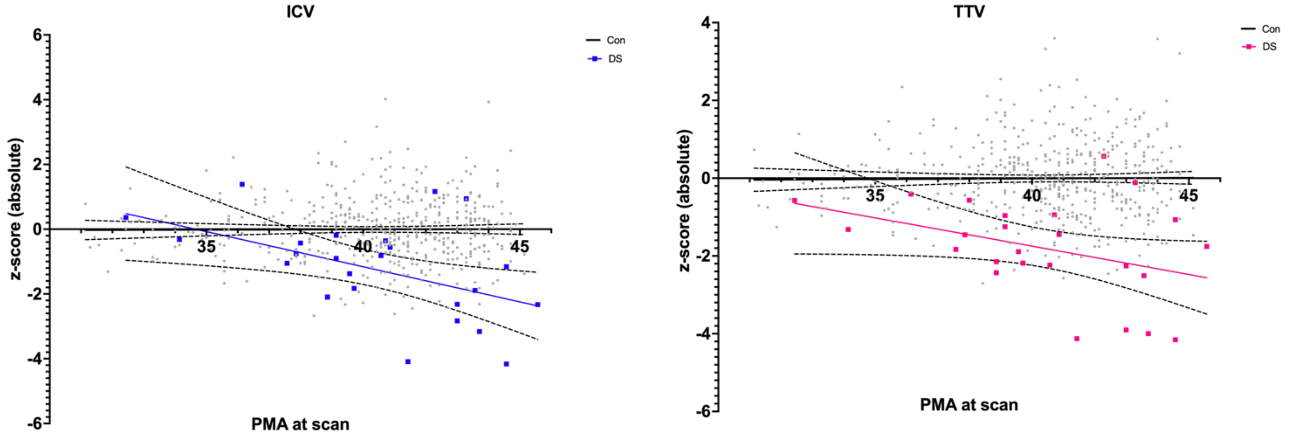

**(See next pages for B, C, D).**

### B. Cortical GM segments

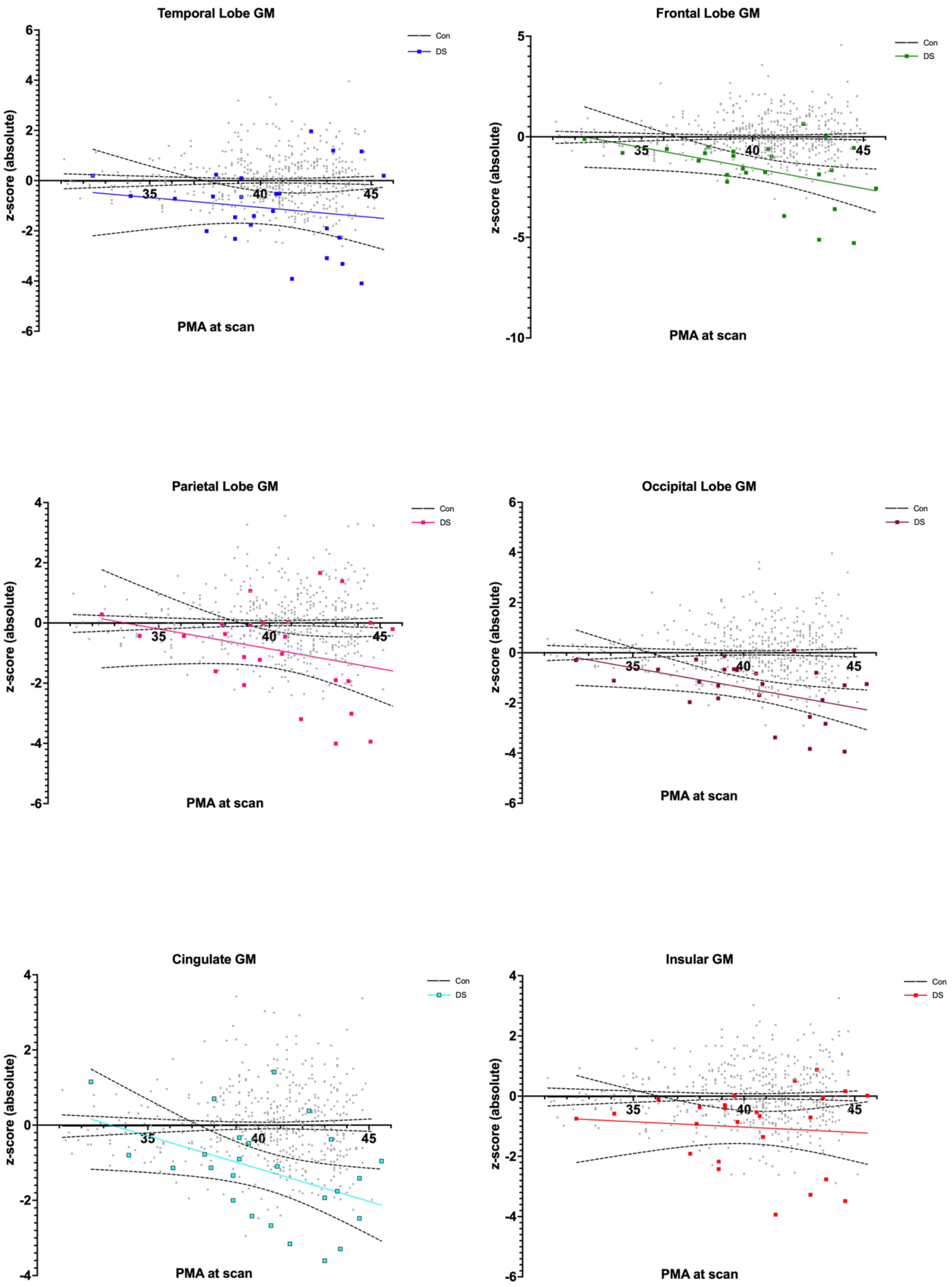

### C. WM segments

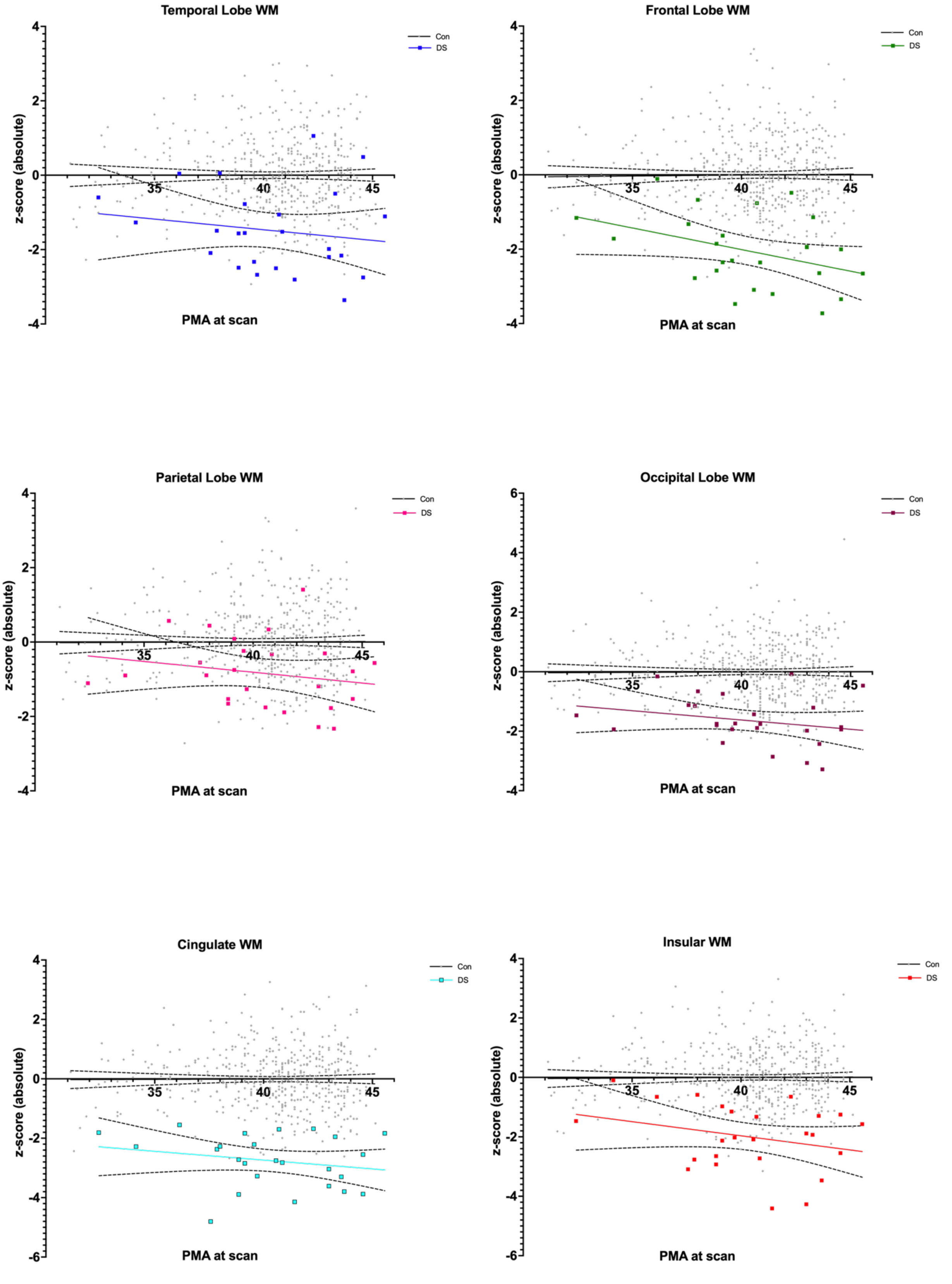

### D. Deep GM and Other segments

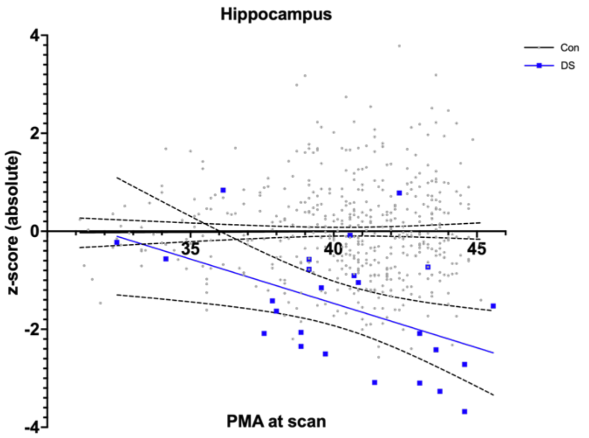

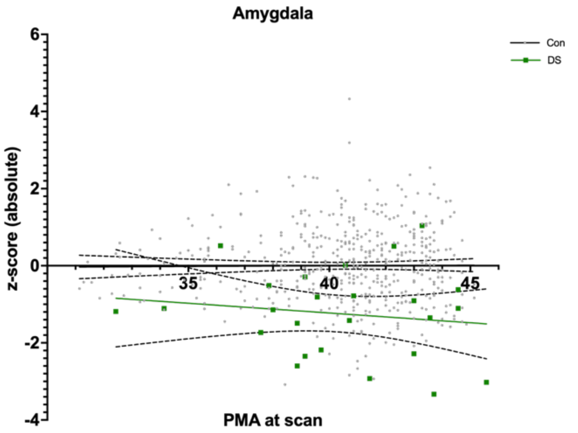

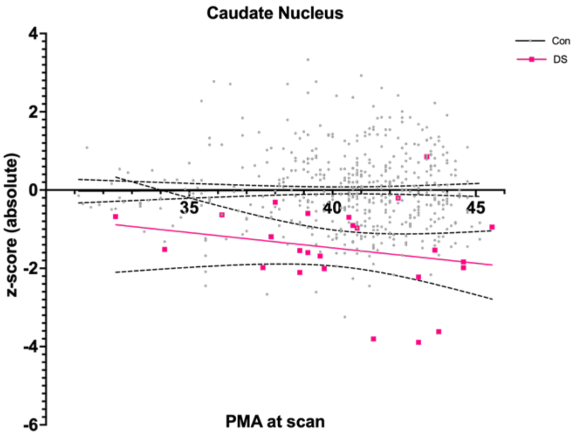

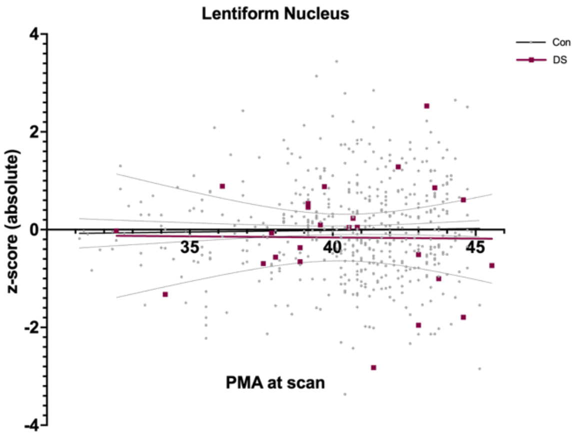

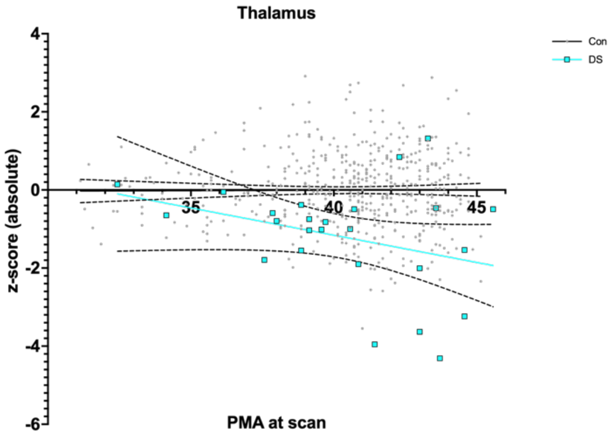

### **Figure S4:** CHD+ and CHD- simple linear regressions by brain segment (using absolute z-scores).

Simple linear regression plots of absolute z-scores against PMA at scan (in weeks) for CHD+ (n = 13, in pink) and CHD- (n = 12, in purple) neonates with DS. Simple linear regressions appear as coloured lines with 95% confidence intervals as dotted lines. Dots for individual control neonates (n = 493, males & females) appear in grey, and simple linear regressions appear as flat black lines at z = 0 with 95% confidence intervals as dotted lines. Plots for **A.** whole brain, **B.** cortical GM segments, **C.** WM segments, **D.** the total deep GM and other segments. **(**Main tissue volumes can be found in Figure 7 of main text**).** A table of results for F-tests can be found in Table S10 and Spearman's correlation in Table S11.

**A. Whole brain volumes (CHD+ and CHD-)**

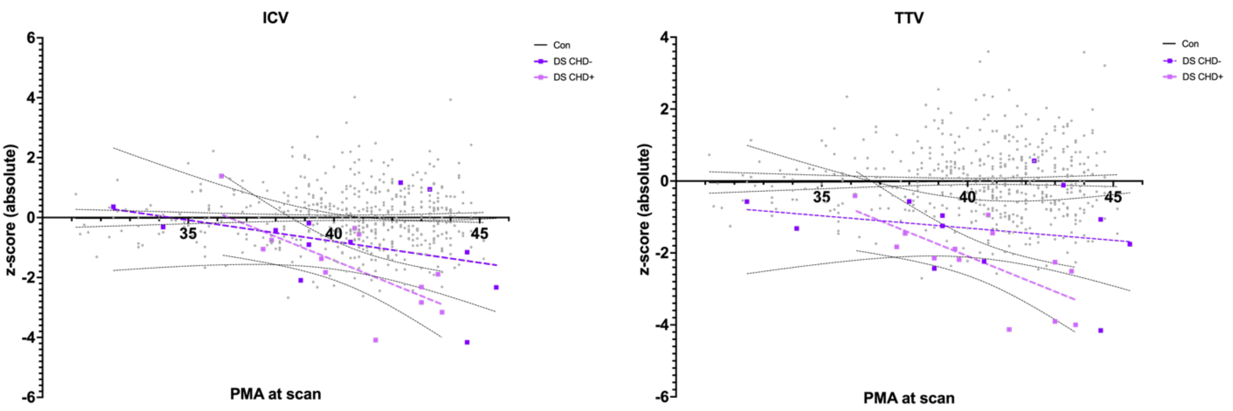

**(See next pages for B, C, D).**

### B. Cortical GM segments (CHD+ and CHD-)

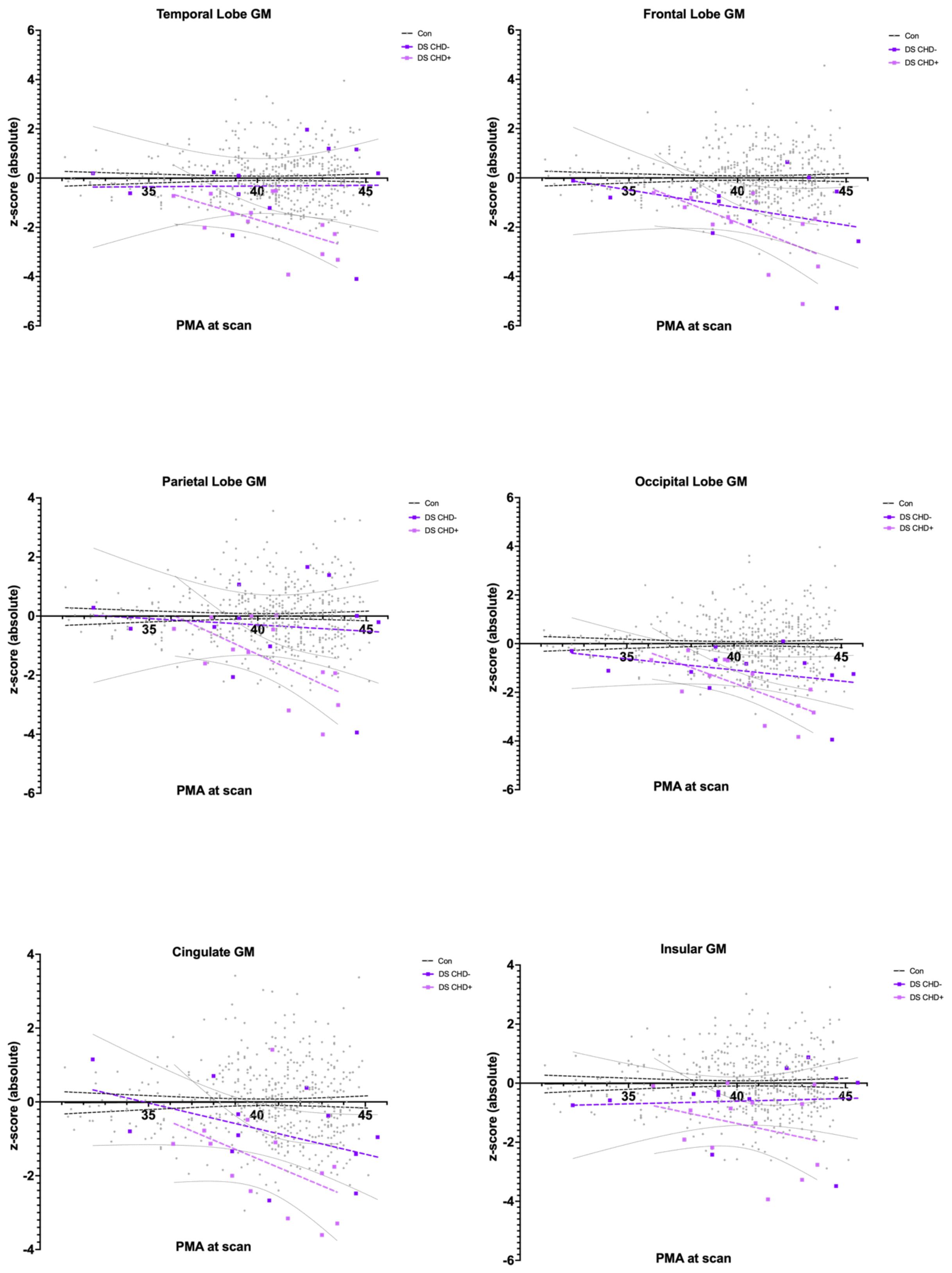

### C. WM segments (CHD+ and CHD-)

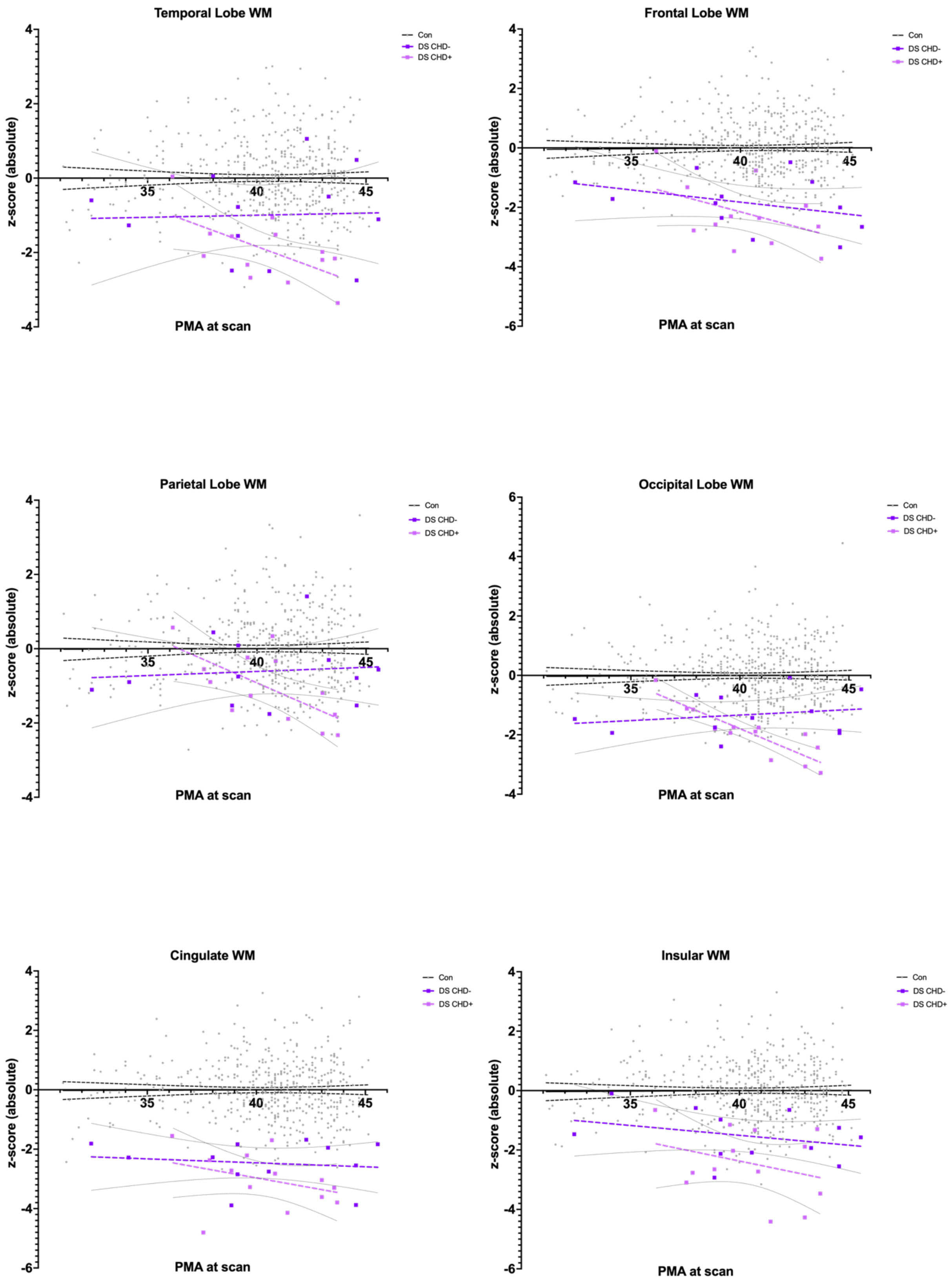

**D. Deep GM and other segments (CHD+ and CHD-)**

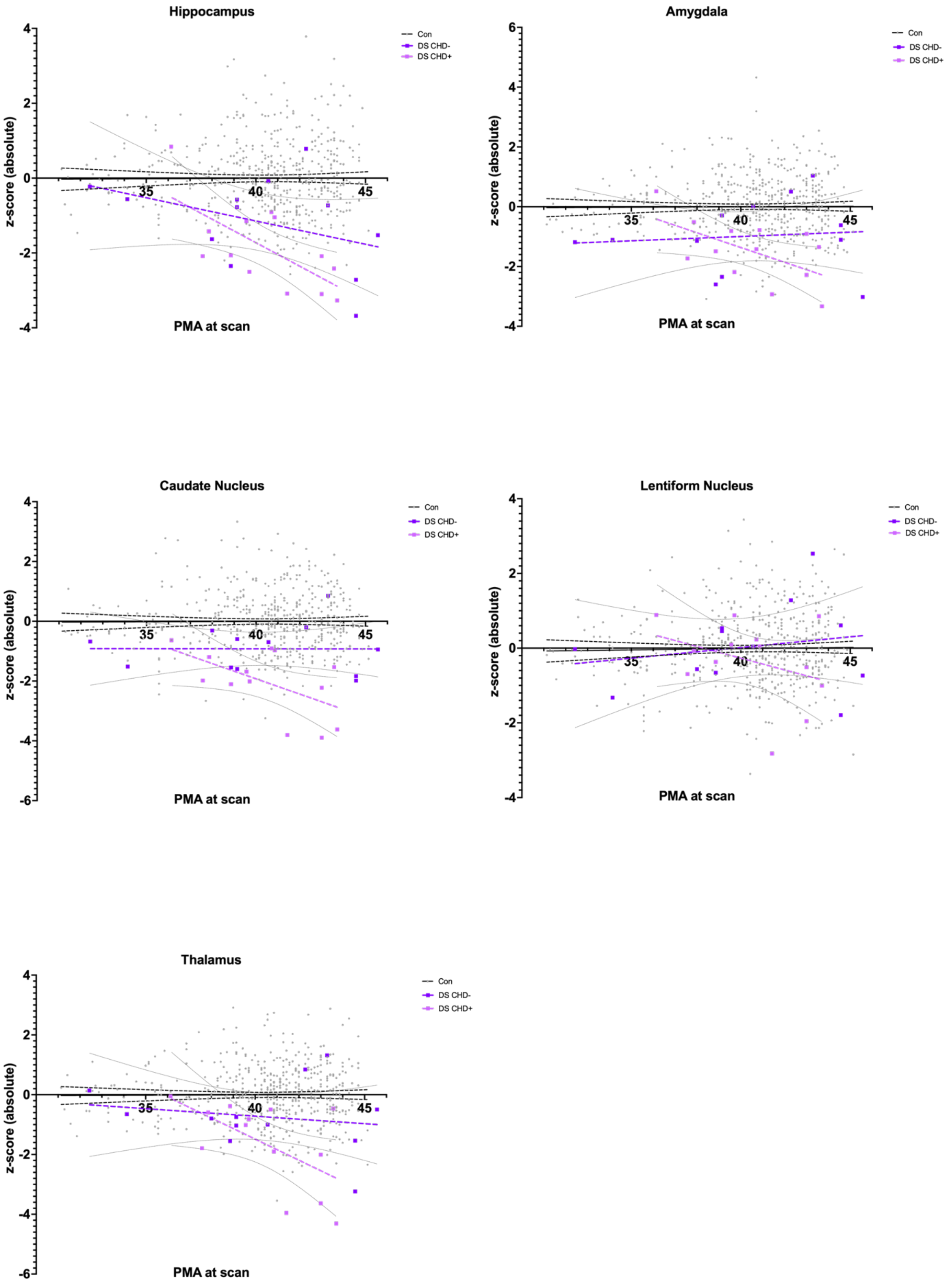

### **Figure S5:** DS and control simple linear regressions by brain segment (using relative z-scores).

Simple linear regression plots of relative z-scores against PMA at scan (in weeks) appear as coloured lines with 95% confidence intervals as dotted lines for neonates with DS (n = 25, males & females). Dots for individual control neonates (n = 493, males & females) appear in grey, and simple linear regressions appear as flat black lines at z = 0 with 95% confidence intervals as dotted lines. Plots for **A.** main tissue volumes **B.** cortical GM segments, **C.** WM segments, **D.** the total deep GM and other segments**.** A table of results for F-tests can be found in Table S7 and Spearman's correlation in Table S8.

**(See next pages).**

### A. Main Tissue Volumes

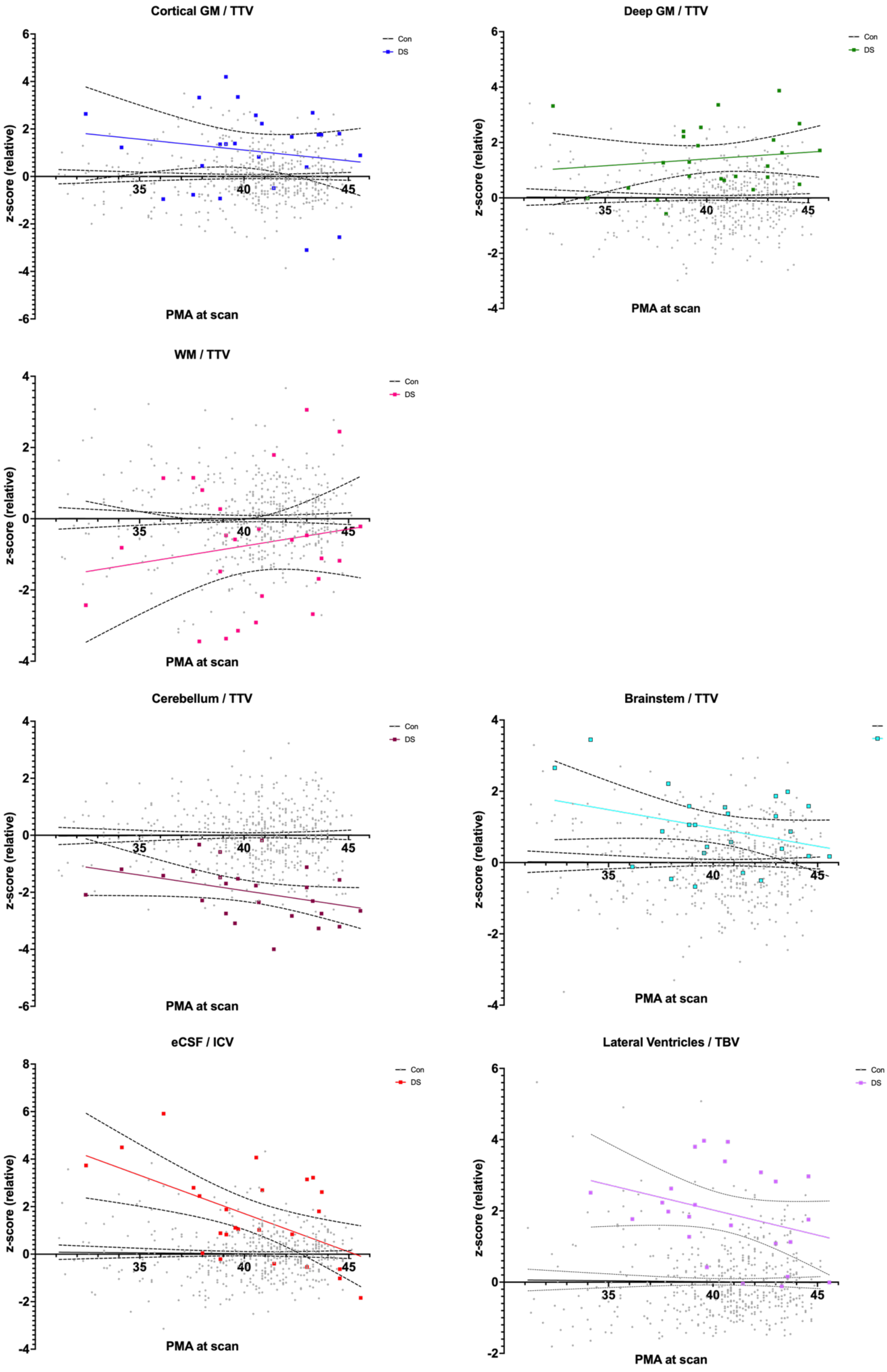

### B. Cortical GM segments

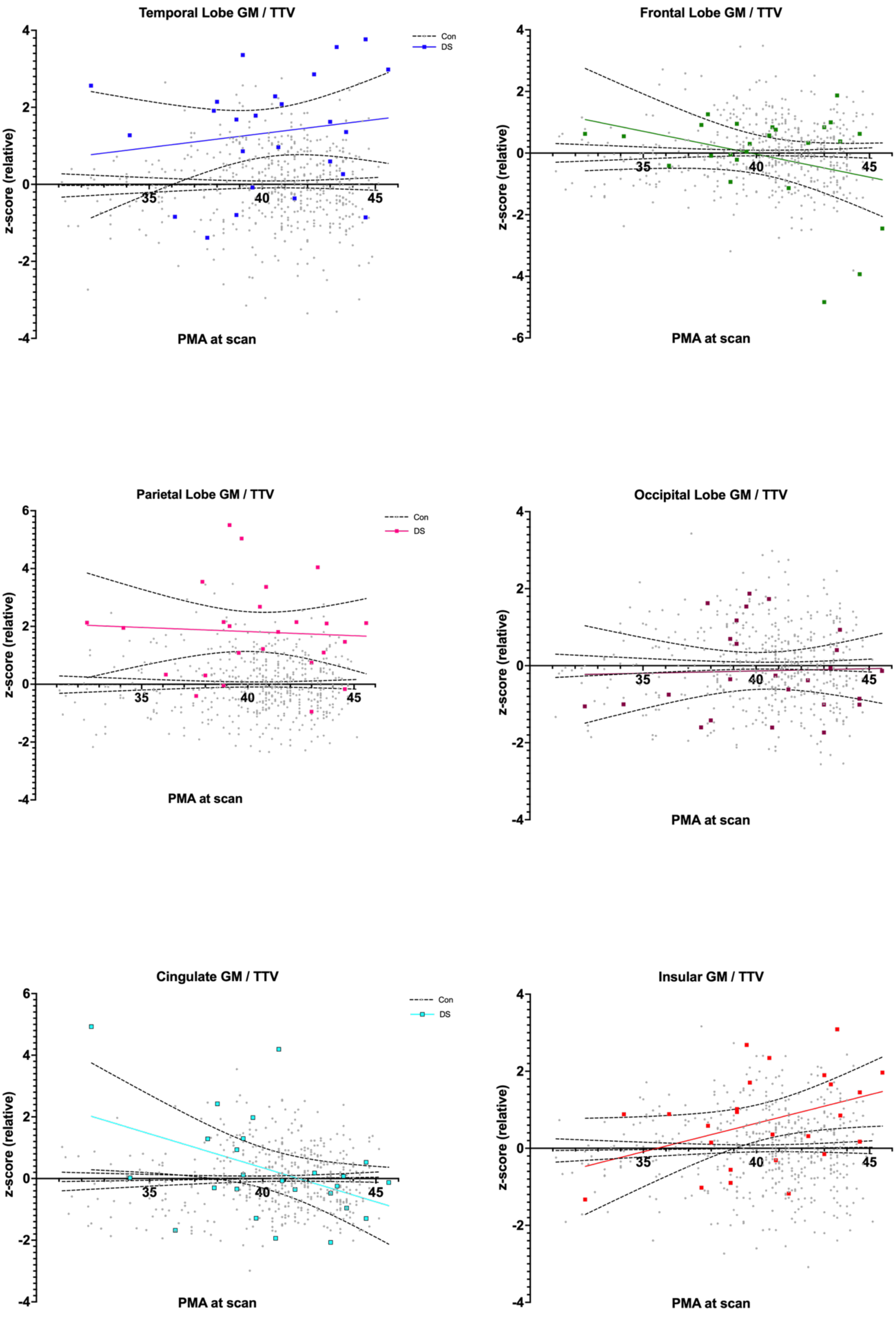

### C. WM segments

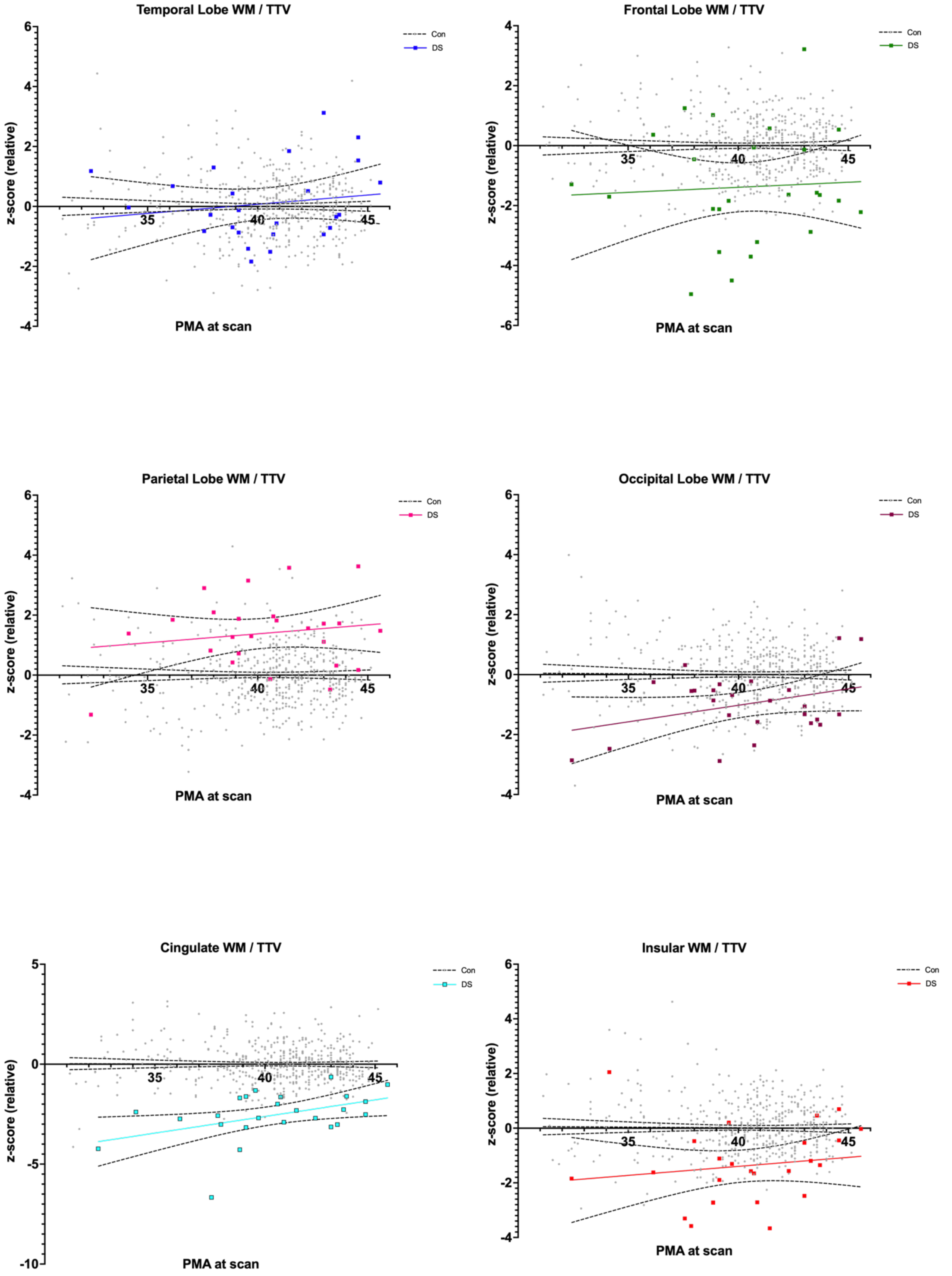

### D. Deep GM and other segments

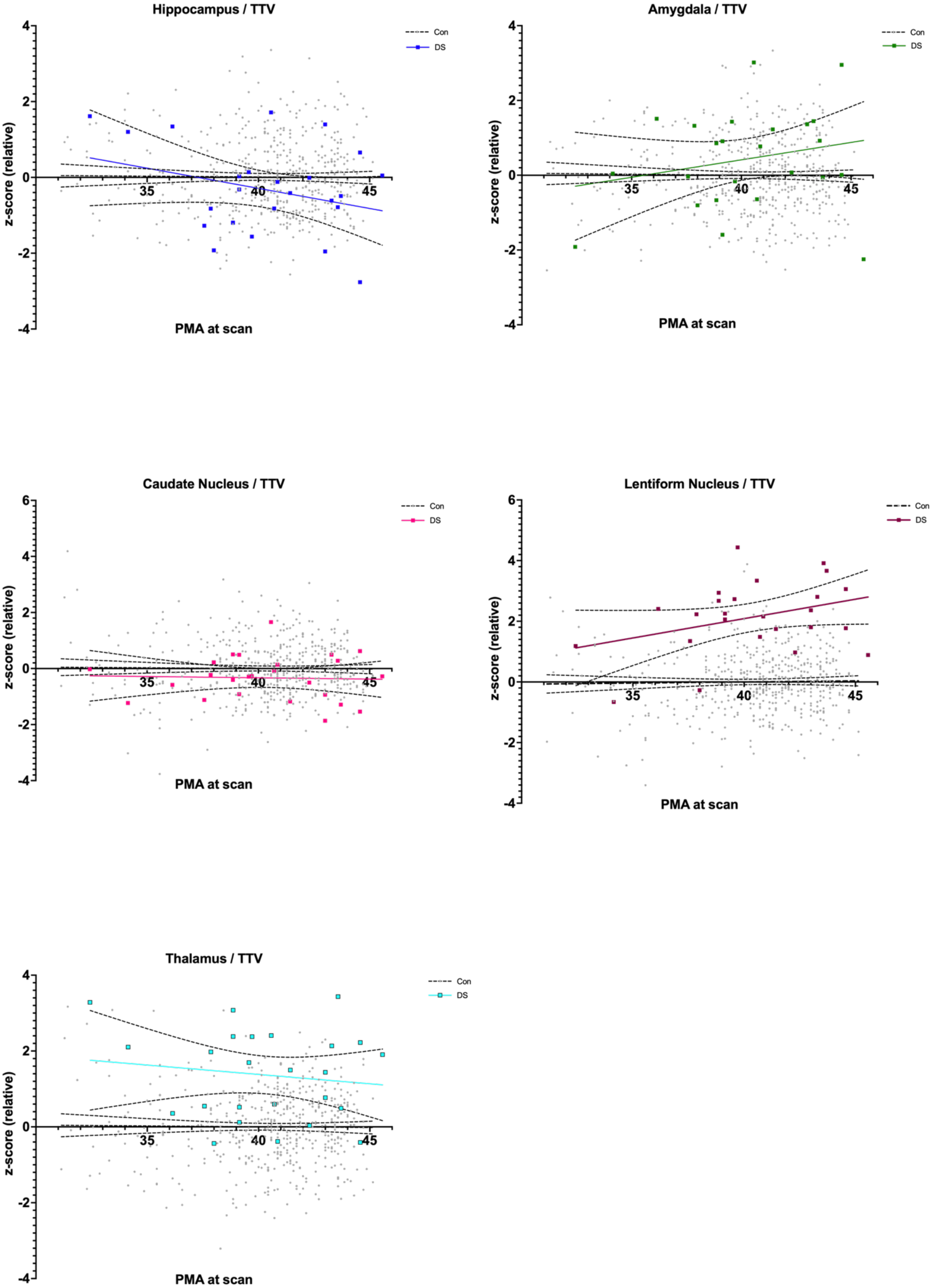
